# A single-cell fixed RNA profiling of liver fibrosis progression and regression reveals SEMA4D and LMCD1 as key mediators of fibrogenesis

**DOI:** 10.1101/2025.06.12.657772

**Authors:** Pham Minh Duc, Le Thi Thanh Thuy, Hoang Hai, Nguyen Thi Ha, Pham Tuan Anh, Hiroko Ikenaga, Hideki Fuji, Hideto Yuasa, Tsutomu Matsubara, Vu Thuong Huyen, Yi Cheng, Ryota Yamagishi, Naoko Ohtani, Daisuke Oikawa, Fuminori Tokunaga, Tatiana Kisseleva, David Brenner, Yasuko Iwakiri, Jordi Gracia-Sancho, Norifumi Kawada

## Abstract

Liver fibrosis progression and regression are dynamic processes involving diverse hepatic and immune cell populations. Here, we utilize single-cell fixed RNA profiling (FLEX) of a TAA-induced mouse liver cirrhosis model, with and without a recovery phase, to depict the cellular landscape and molecular mechanisms of fibrosis resolution. The regression phase was characterized by the emergence of pericentral hepatocytes enriched in detoxification and antioxidant genes (e.g., Cyp2e1, Txn1), which secreted Rarres2 to modulate hepatic stellate cell (HSC) function. This was accompanied by the upregulation of scar-resolving genes (Mmp14, Ctsl), restoration of fenestrae in liver sinusoidal endothelial cells, anti-inflammatory phenotypes of Kupffer cells, a decrease in fibrogenic cholangiocyte subsets, and recovery-associated signatures in NK/T cells, B cells, and neutrophils. In contrast, SEMA4D secreted by monocyte-derived macrophages during fibrosis progression activated Plxnb2⁺ HSCs, and its blockade attenuated fibrosis in vivo. Furthermore, LMCD1 was identified as a novel marker for HSC activation and regulation. This single-cell atlas reveals key transcriptional programs and intercellular signaling pathways dependent on the fibrotic condition, offering new therapeutic targets for liver cirrhosis.

## Introduction

Cirrhosis is a prevalent pathological condition associated with advanced chronic liver diseases induced by hepatitis virus infection, alcohol abuse and metabolic dysfunction-associated steatotic liver disease, often leading to decompensated cirrhosis and hepatocellular carcinoma (HCC)^1^. Over the past four decades, remarkable advancements have been made in understanding the biological mechanisms underlying the progression of liver fibrosis to cirrhosis; however, urgent unmet medical needs have remained to yield effective therapies for liver fibrosis and cirrhosis resolution to improve clinical outcomes for patients^2^.

A critical aspect of both clinical and experimental liver cirrhosis is the potential for disease regression. Landmark studies from the 1970s documented instances of regression of liver fibrosis and cirrhosis in various animal models and human liver diseases. Today, the reversal, at least partially, of human cirrhosis has been demonstrated in clinical practice, supported by serial biopsy assessments in patients eradicated by hepatitis virus C infection^3^. Despite these knowledge, the mechanisms driving cirrhosis regression remain largely elusive, partly due to the limitations of preclinical rodent models, which exhibit easy regeneration and rapid reverse and hardly reflect human pathological process^4^. While our understanding of the cellular mechanisms involved in liver fibrosis progression and regression has expanded, the focus of research in this field has predominantly centered on the functional analyses of hepatic stellate cells (HSCs)^5–10^ and the comprehensive understanding on the role of cell-cell interaction among hepatic constituent cells, i.e. hepatocytes (HEPs), liver sinusoidal endothelial cells (LSECs), macrophages (MACs), hepatic stellate cells (HSCs), cholangiocytes (CHOLs), natural killer T (NK&T) cells, neutrophils (NEUs), dendritic cells (DCs), B cells and mesothelial cells remains largely unknown.

Previous reviews have highlighted key molecular mechanisms underlying the reversibility of liver fibrosis, such as apoptosis and inactivation of HSCs, crosstalk between HSCs and bone marrow-derived MACs, and the interactions among these cell types that induce apoptosis in myofibroblasts^11–13^. Understanding these cellular and molecular mechanisms has sparked extensive research into new pharmacological strategies aimed at enhancing these processes. Current pharmacological therapies are typically prophylactic during the preclinical induction of chronic liver injury or administered shortly after the establishment of mild disease in mice. However, in clinical trials, drugs need to be developed that can be administered to patients with already established stage 3 or 4 fibrosis and have the pharmacological effect of eliminating fibrosis bundles.

Recent advancements have opened new avenues for a more nuanced understanding of molecular mechanism of cirrhosis^14^. Notably, single-cell RNA sequencing (scRNA-seq) has been employed to create comprehensive atlases of hepatic constituent cells in both human and mouse models of intact and diseased livers. These studies have typically focused on total liver cells, CD45^+^ liver cells, non-parenchymal liver cells, or HSCs^15–18^. However, sample quality remains a critical factor influencing sequencing outcomes and data analysis. Enzymatic digestion of whole liver tissue necessitates careful perfusion of solutions at specific flow rates, as improper perfusion can lead to tissue damage and alterations in gene expression profile^19–21^. One notable limitation of single nucleus RNA-seq (snRNA-seq) is its inability to sequence cytoplasmic RNA, as it exclusively collects and analyzes nuclear transcripts^22^. snRNA-seq often required more tissue to compensate for loss of material and more additional steps compared to scRNA-seq. On the contrary, single-cell fixed RNA sequencing (FLEX) represents an innovative approach for isolating cells from frozen tissues, preserving RNA through paraformaldehyde (PFA) fixation. This method yields a higher number of unique molecular identifiers (UMIs) and genes, while reducing the percentage of mitochondrial genes compared to methanol fixation. Fixed droplet RNA sequencing is particularly advantageous for studying rare cell subpopulations, offering researchers increased flexibility in high-throughput scRNA-seq applications^23^.

To gain comprehensive insights into the phenotypic alterations of liver cell types and their interactions contributing to cirrhosis development, we are the first group employed this FLEX method on frozen liver tissues from normal, thioacetamide (TAA)-induced cirrhosis, and its regression phases in mice. Our analysis revealed the phenotypic alterations in ten primary hepatic cells, i.e., HEPs, LSECs, MACs, HSCs, CHOLs, NK&T cells, NEUs, DCs, B cells and mesothelial cells alongside their interactions during cirrhosis development and regression. Obtained results from this research would contribute to the development of effective antifibrotic therapies in the future targeted to LIM and cysteine rich domains 1 (LMCD1) and semaphorin-4D (SEMA4D).

## Results

### FLEX unveiled hepatic cell populations atlas comprehensively during progression and regression of liver cirrhosis in the thioacetamide-induced mouse model

To elucidate the cellular landscape of liver cirrhosis progression and regression, we employed a thioacetamide (TAA)-induced mouse model of advanced liver fibrosis. TAA administration for 10 weeks (Supplementary Fig. 1a) led to significant liver injury (Supplementary Fig. 1b), characterized by bridging fibrosis between portal and central veins accompanied by increase in ACTA2^+^ activated HSCs, marked CD68^+^ macrophage infiltration and proliferation of Ki67^+^ hepatocytes (Supplementary Fig. 1c). In contrast, these changes improved when TAA administration was discontinued and livers undergoing spontaneous fibrosis regression demonstrated a significant reduction in apoptosis as evidenced by the reduction of p21 and BAX, and xenobiotics metabolism capacity by increasing the level of CYP2E1 (Supplementary Fig. 1a–e). Cytokine and chemokine profiling from mouse serum revealed elevated levels of regenerative cytokines inclusing IL4, IGFBP1 and M-CSF in fibrosis regression group (Supplementary Fig. 1f).

To dissect cell-type-specific dynamics and intercellular interactions during cirrhosis progression and regression, we performed single-cell fixed RNA profiling (FLEX) using the 10X Chromium platform, analyzing n = 3 biological replicates per group (Fig. 1a). Following integration via Seurat’s reciprocal PCA method, and normalization at the gene and cell levels, we obtained transcriptomic data from 38,136 cells - 13,948 from the control, 14,986 from the cirrhosis, and 9,202 from the regression group - with an average sequencing depth of 4,426 genes per cell (Supplementary Data 1). Based on canonical markers, we annotated 10 major hepatic cell types (Fig. 1b–d), including HEPs expressing Alb, Ttr, Trf, Serpina1a1, Hnf4a, and Aox3; LSECs expressing Kdr, Aqp1, Pecam1, and Eng; MACs including Kupffer cells expressing Vsig4, Adgre1, Clec4f, Lgals3, Ccr2, and Cd68; HSCs expressing Rgs5, Colec11, Lrat, Dcn, Cygb, and Des; CHOLs expressing Epcam, Krt7, Spp1, Krt19; B cells expressing Cd79a, Cd79b, Ebf1, and Ms4a1; NK&T cells expressing Cd3d, Il7r, Nkg7, and Gzma; mesothelial cells expressing Msln and Upk3b; NEUs expressing Cxcr2 and S100a9; and DCs expressing Clec9a, Siglech, and Cox6a2 (Fig. 1e and Supplementary Data 1). Comparative analysis of cell-type proportions across the three groups revealed that cirrhotic livers displayed an increased proportion of MACs and immune cells (B cells, NK&T cells, NEUs, and DCs), CHOLs, and HSCs, alongside reduced abundance of HEPs and LSECs compared to the control and regression groups (Fig. 1f and Supplementary Data 1).

**Fig. 1.**
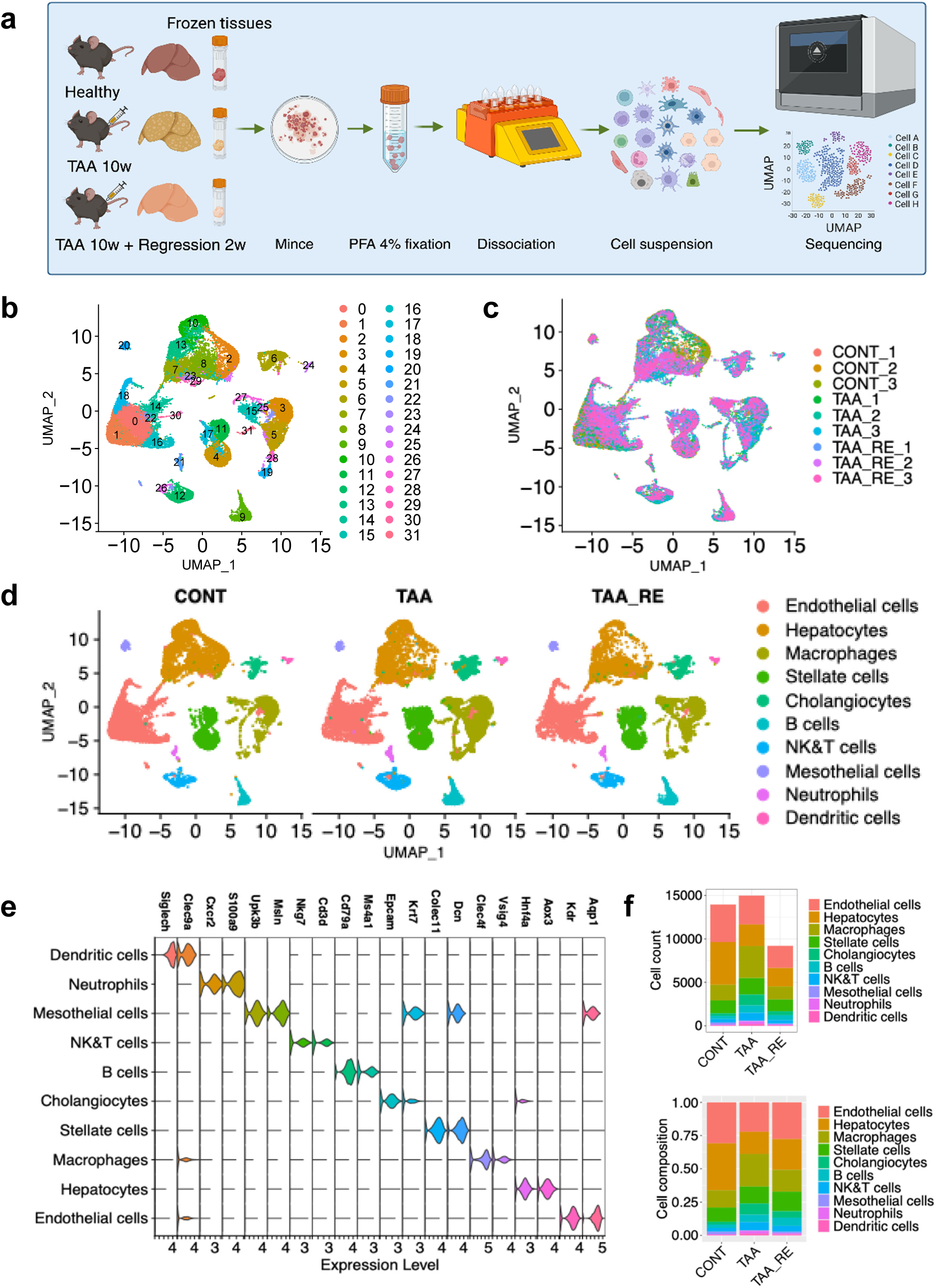
Single cell fixed RNA profiling (FLEX) of mouse livers from control, TAA treated and recovered groups. **a** Single cell fixed RNAseq workflow using the 10X Genomics Chromium Flex platform. Created in BioRender. Duc, P. (2025) https://BioRender.com/zo1s4v7. **b** Uniformed Manifold Approximation and Projection (UMAP) showing 32 cell populations of integrated data. Each dot represents a single cell. Cells from different clusters are marked by colors **c** UMAP showing 9 datasets integration from the control, cirrhosis, and regression group (n=3, each group). Colors denoted samples. **d** UMAP plots of 13948 cells from the control group, 14986 cells from the cirrhosis group, and 9202 from the regression group; showing 10 annotated cell populations of integrated data. The identity of each cluster was determined by matching the expression profiles of clusters with established cell-specific marker genes of different hepatic cells. Each cluster was shown in different color. Rpca package was used to correct batch effect and constructed one UMAP based on all cells from nine datasets, and then split cells by three groups. **e** Stacked violin plots showing the average expression of specific markers in indicated cell types. **f** Cell counts (upper panel) and cell composition (lower panel) of 10 major cell types across three groups (control, cirrhosis, regression). Colors indicated different cell types

### RecoHEPs^Cyp2e^^1, Cyp1a2, Glul, Oat, Mgst1, and Txn1^ define a hepatocyte subpopulation with enhanced xenobiotic and antioxidant functions during fibrosis regression

To further resolve HEP heterogeneity, we reclustered HEPs from all three conditions, identifying 20 transcriptionally distinct subclusters (Fig. 2a-b, Supplemetary Data 2) distributed in three phases of the cell cycle with increased G2/M phase in fibrotic hepatocytes (TAA treated group) (Fig. 2c) which was consistent with dominant Ki67 expression shown in Supplementary Figure 1c. Subclusters S1, 3, 5, 7, 9, 11, 13, 14, and 17 were predominantly derived from control livers, while subclusters S2, 4, 6, 12, 18, and 19 were enriched in the cirrhosis group. Notably, clusters S0, 8, and 16 were composed primarily of HEPs from the regression group (Supplementary Fig. 2a). Given that hepatocyte zonation, the spatial arrangement of different metabolic functions within liver lobules, is crucial for maintaining overall liver function and can be implicated in various liver diseases^24, 25^, we have analysed HEP zonation based on its pericentral or periportal signature score using AddModuleScore (Fig. 2d, Supplementary Fig. 2b). Gene expression heatmaps and gene set enrichment analysis (GSEA) demonstrated that the transcriptional profiles of pericentral HEPs in regression phase resembled those of healthy controls (Fig. 2e-f, and Supplemetary Data 2). Among these, a distinct subcluster, which was termed recoHEPs^Cyp2e1, Cyp1a2, Glul, Oat, Mgst1, Txn1^, exhibited high gene expression of xenobiotic metabolism (Cyp2e1, Cyp1a2), Glutamine synthetase (Glul), Ornithine aminotransferase (Oat), and the antioxidants such as Microsomal glutathione s-transferase 1 (Mgst1) and Thioredoxin 1 (Txn1) (Fig. 2e, g, and Supplementary Data 2), being predominantly localized in the pericentral region. These genes were significantly upregulated in the regression group compared to the cirrhosis group based on differential gene expression (Supplementary Fig. 2c). In contrast, periportal HEPs displayed minimal changes between fibrotic and regressed livers (Supplementary Fig. 2d). Immunohistochemical staining for E-cadherin, a cell adhesion molecule primarily expressed in periportal hepatocytes within the liver lobules^26^, clearly highlighted the periportal region (zone 1) in the healthy mouse liver group, but zone 1-E-cadherin positivity was lost in the fibrotic liver, and did not appear to be reconstructed during the recovery phase (Supplementary Fig. 2f). Meanwhile, immunohistochemical staining for GLUL and CYP2E1 demonstrated a marked recovery of these two proteins in the pericentral region (Fig. 2h). The fact that recovery occurs in the pericentral rather than the periportal region may represent the initial stage of liver functional restoration or fibrosis regression, as observed above.

**Fig. 2.**
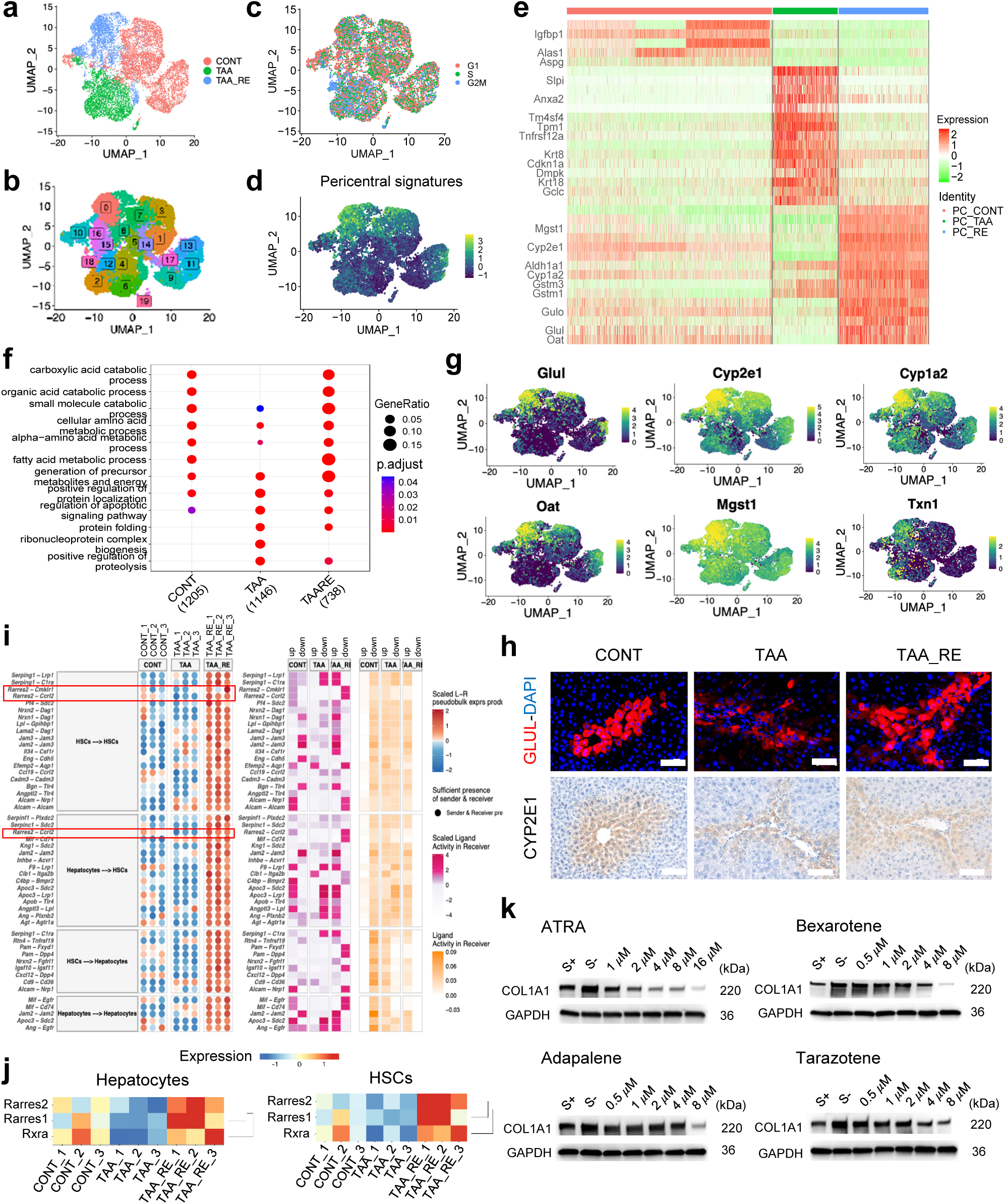
Characteristic of hepatocytes with enhanced xenobiotic and antioxidant functions during fibrosis regression. **a** UMAP visualization hepatocytes from control, cirrhosis, and regression group, colored by group. **b** UMAP of 20 hepatocytes subpopulations, colored and labeled by cluster identity. **c** UMAP showing cell cycle analysis of hepatocytes subpopulations, colour by cell cycle phase prediction. **d** UMAP showing pericentral signature score of hepatocytes subpopulations. **e** Heatmap showing representative genes expressed by pericentral hepatocytes in the control, cirrhosis, and regression groups. PC: Pericentral hepatocytes **f** Dot plot represented pathways distinguishing each group. Dot sizes corresponded to the ratio of pathway genes found in the dataset. Colors corresponded to p-value adjustment. Genes sets used for the analysis were taken from the Gene Ontology (GO) dataset. **g** Feature plots showing the relative distributions of Glul, Cyp2e1, Cyp1a2, Oat, Mgst1, and Tnx1 expression among all subpopulations from lowest (dark green) to highest (yellow). **h** Immunofluorescence staining with Glutamine synthetase (red) and DAPI (green) (upper). IHC staining shows positive area of CYP2E1 staining. Scale bar 50 𝜇𝜇m. **i** Bubble plot showing the top 50 ligand-receptor (L-R) interactions between hepatocytes and HSCs in the regression phase. Bubble colors and sizes represent the per-sample scaled product of normalized ligand and receptor pseudo-bulk expression and sufficient presence L-R. Heatmap colors indicated the ligand activity and scaled ligand activity in the receiver. **j** Heatmap showing the relative expression of Rarres1, Rarres2, and Rxra among groups in hepatocytes and HSCs, colored by gene expression **k** Representative immunoblotting showing expression of COL1A1 in HHSteCs treated dose-dependently with retinoic receptor agonists (ATRA, Bexarotene, Adapalene, and Tarazotene). GAPDH was used as an internal control.

This prompted us to firstly investigate the interactions between HEPs and HSCs involved in liver fibrogenesis using the MultiNicheNet analytical approach. The results revealed that among the top 50 statistically significant receptor-ligand interactions, many occurred from HEPs to HSCs or between HSCs themselves (Fig. 2i). Notably, interactions such as Retinoic acid receptor responder protein 2 (Rarres2)–Cmklr1 and Rarres2–Ccrl2 were strongly enriched during the recovery phase (Fig. 2i), and Rarres2–Ccrl2 was only found from HEPs to HSCs at recovered group (Supplementary Fig.3a).

Heatmap analysis also showed a prominent upregulation of transcriptional Rarres2 in both HEPs and HSCs in the recovery group, whereas its expression was downregulated in the fibrotic liver group (Fig. 2j), consistent with its transcriptional expression (Supplementary Fig.3b) and its protein level increased in the serum of recovered mice found in Supplementary Figure 1f.

Given that Rarres2 expresssion increased in regenerated hepatocytes and regressed HSCs (Fig. 2j), modulating Rarres2 could be protective against fibrosis progression by mitigating HSCs activation, we treated human HSCs (HHSteCs) with one first-generation Rarres2 agonist, all-trans retinoic acid (ATRA) and three third-generation agonists: Bexarotene, Adapalene, and Tazarotene^27^. At the same concentration of 5 µM, ATRA demonstrated the strongest ability to induce RARRES2 expression in HHSteCs (Supplementary Fig.3c). Moreover, dose-dependent treatment with ATRA led to a marked inverse correlation with COL1A1 expression, indicating a significantly stronger inhibitory effect on collagen synthesis compared to the other three agonists (Fig. 2k, Supplementary Fig.3d). Collectively, our findings indicate that subpopulation of pericentral hepatocytes may contribute to drive hepatocyte regeneration via enhancing xenobiotics metabolism and antioxidant capacity and Rarres2 could be one of potential biomarkers of resolution phase of cirrhosis.

### Remodeling of liver sinusoidal endothelial cells during cirrhosis regression

While previous studies have reported zonation-specific changes in LSECs under normal and fibrotic conditions^15, 18, 28^, their phenotypic dynamics during cirrhosis regression remain poorly defined. To address this, we reclustered LSECs from control, cirrhotic, and regression groups into nine transcriptionally distinct subclusters (Fig. 3a–b). Heatmap analysis revealed distinct gene expression profiles (Supplementary Fig. 4a), with subclusters S2–S4 enriched in cells from healthy livers; S1 and S5 from cirrhotic livers; S0 predominantly from the regression group; S6 from a mix of regression and healthy samples; and S7 and S8 comprising cells from all three conditions (Fig. 3c). According to zonation-specific markers^15, 18, 28^, subcluster S7 expressing Adgrg6, Vegfc, Jag1, and Npr3, corresponds to reported portal LSECs, whereas S8 expressing Rspo3, Wnt9b, Bmp4, and Selp, corresponds to reported central LSECs (Fig. 3d). Spatial mapping suggested S2 and S4 localized to the periportal and midzonal regions and were marked by Adam23, Msr1, Cd36, and Lyve1, while S3, expressing Wnt2, Thbd, and Kit, was positioned in the pericentral zone. Gene set enrichment analysis (GSEA) revealed that healthy LSEC clusters (S2–S4) were enriched in small GTPase signaling, Ras pathways, cell adhesion, metal ion response, and lipid and small molecule metabolism (Fig. 3e). By contrast, cirrhotic clusters S1 and S5 showed reduced expression of these pathways and upregulation of genes related to angiogenesis, endothelial development, migration, and junction assembly. Notably, S1 was enriched in inflammatory and apoptotic genes, including Cdkn1a, Bax, Dmpk, Il1a, Hic1, and Tnfrsf12a (Supplementary Fig. 4b; Supplementary Data 3) while S5 showed elevated ECM signatures, including Col4a1, Col4a2, Col1a1, Col1a2, Bcam, Ednrb (Supplementary Fig. 4c; Supplementary Data 3). Human cirrhotic LSECs in public dataset showed similarity with our current analysis (Supplementary Fig. 4d).

**Fig. 3.**
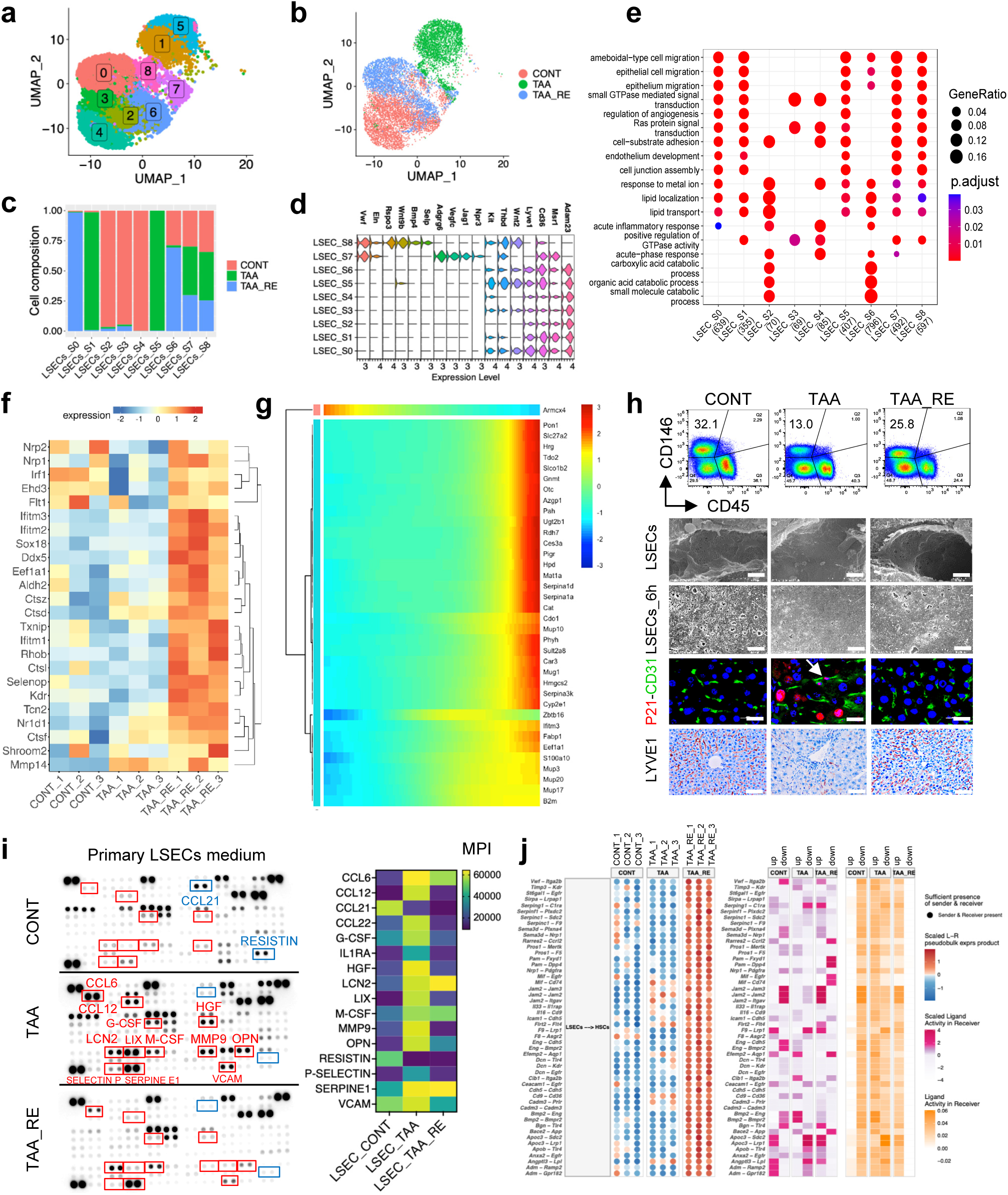
Remodeling of liver sinusoidal endothelial cells during cirrhosis regression. **a** UMAP showing 9 subpopulations of LSECs from control, cirrhosis, and the regression group, colored by cluster. **b** UMAP showing LSECs from control, cirrhosis, and regression group, colored by group. **c** Barplot showing group composition breakdown per cluster (fraction of total cell count per cluster) **d** Stacked violin plot showing markers of LSECs landscape and mixed clusters **e** Dot plot represented pathways distinguishing each LSECs subpopulation by ClusterProFiler. Dot sizes corresponded to the ratio of pathway genes found in the dataset. Colors corresponded to p-value adjustment. Genes sets used for the analysis were taken from the Gene Ontology (GO) dataset. **f** Heatmap displayed up-regulation of ECM-degrading enzymes (Ctsl, Ctsd, Ctsf, Ctsz, and Mmp14), interferon-stimulated genes (Ifitm1–3), and antioxidant genes (Selenop, Tcn2, and Txnip) in the regression group. **g** Heatmap of the top genes that were differentially expressed along the pseudotime, colored by gene expression. **h** Flow cytometry showing CD146 and CD45 positive cells among control, cirrhosis and the regression group (upper panel). Scanning electron microscopy (SEM) images showed morphology and fenestration of LSECs in liver tissues from control, cirrhosis and regression. Scale bar 2.5 𝜇𝜇m. SEM image showed morphology and fenestration of isolated LSECs from control, cirrhosis and the regression group after culturing 6 hours in medium. Scale bar 2.5 𝜇𝜇m. Double IF staining for P21 (red), CD31 (green), and DAPI (blue) for nuclear staining confirms colocalized of P21-CD31 in cirrhosis livers (white arrow). IHC staining showing expression of Lymphatic vessel endothelial receptor 1 (LYVE1) in healthy, fibrosis and regressed group (lower panel). Scale bar 50 𝜇𝜇m. **i** Cytokine array membrane of soluble factors in cultured medium of LSECs isolated from control, cirrhosis and the regression group (left panel) and heatmap showing the quantification of mean pixel density (MPI, right panel). The soluble factors with predominant change in TAA were marked with red boxes, the soluble factors with predominant change in healthy control (CONT) were marked with blue boxes. **j** Bubble colors and sizes represent the per-sample scaled product of normalized ligand and receptor pseudo-bulk expression and sufficient presence L-R. Heatmap colors indicated the ligand activity and scaled ligand activity in the receiver.

LSECs in recovery phase showed heterogeneous states. Subcluster S0 remained similar to S1 and S5, suggesting persistent injury, whereas S6 resembled healthy S2 with partial restoration of homeostatic signaling. Regression-associated subpopulations exhibited increased expression of ECM-degrading enzymes (Ctsl, Ctsd, Ctsf, Ctsz, and Mmp14), interferon-stimulated genes (Ifitm1–3), and antioxidant genes (Selenop, Tcn2, and Txnip) (Fig. 3f). Trajectory analysis using Monocle 2 revealed that LSEC differentiation dynamics (Supplementary Fig. 4e) with hierarchical clustering highlighting transcriptional transitions during recovery (Fig. 3g). Recovered LSECs upregulated genes supporting antioxidant defense Pon1^29^, lipid metabolism, immune modulation, and nitrogen clearance such as Slc27a2, Tdo2, Gnmt, and Otc. To examine the phenotype of LSECs at recovery state, we isolated primary liver non-parenchymal cells from the three groups. Flow cytometry analysis using CD146, a transmembrane protein expressed on endothelial cells, and CD45 gating which helps distinguish LSECs from other cell types like leukocytes, reveals that CD146⁺ LSECs populations decreased in fibrotic liver compared with controls (32.1% vs 13%) and regained in recovery phase (25.8%) (Fig. 3h, top). SEM imaging confirmed restoration of LSEC fenestration and immunostaining showed reduced senescence (P21) and re-expression of Lyve1 in regressed livers (Fig. 3h, bottom). Cytokine arrays of LSEC-conditioned media demonstrated elevated angiokines and inflammatory mediators including CCL6, CCL12, CCL22, IL1RA, HGF, LCN2, LIX, MMP9, OPN, G-CSF, M-CSF, P-SELECTIN, and VCAM in cirrhotic LSECs (red boxes), which were markedly reduced upon regression as well as control cells; meanwhile CCL21 and Resistin secreted into the medium were reduced in cirrhotic LSECs compared with control ones (blue boxes) (Fig. 3i, left panel) and their mean pixel density calculated heatmap (Fig. 3i, right panel).

Finally, predicted ligand–receptor interactions between LSECs and HSCs in the regression phase identified Vwf–Itga2b, Timp3–Kdr, and St6gal1–Egfr as top signaling axes (Fig. 3j). The von Willebrand factor (VWF) and integrin αIIbβ3 (Itga2b) axis plays a crucial role in LSECs by facilitating platelet adhesion and thrombus formation in the liver, which is important for hemostasis^30, 31^. The TIMP3-KDR axis functions by TIMP3, a tissue inhibitor of metalloproteinases, inhibiting the binding of vascular endothelial growth factor (VEGF) to its receptor, KDR (VEGFR2). This interaction is crucial for regulating angiogenesis^32^. ST6Gal1-mediated EGFR regulation was recently shown to foster elevated integrin forces, inferring a role in migration^33^. Together, these findings provide a comprehensive map of LSECs remodeling during cirrhosis regression.

### Sema4d+ macrophages are key drivers of liver inflammation and fibrosis

To investigate the role of macrophage subpopulations in liver cirrhosis progression and regression, we performed clustering of macrophages from control, cirrhotic, and regression-phase livers and identified 12 distinct subclusters (Fig. 4a). Based on canonical markers^34, 35^, subclusters S1–S5 and S7 expressed Mrc1, Adgre1, Cd5l, Clec4f, Cd163, Vsig4, Marco, Folr2, and Timd4, consistent with resident Kupffer cells (KC1). Subclusters S0, S6, and S8, enriched in Trem2, Gpnmb, and Cd63 with reduced Folr2 and Cd163, were defined as KC2. S10 cells expressing KC1 markers along with Mki67 and Top2a were classified as proliferating Kupffer cells (KC3). In contrast, subclusters S9 and S11 expressed Itgam and Fcgr2b, and were identified as monocyte-derived macrophages (MoMAC) (Fig. 4b, Supplementary Fig. 5a–b and Supplementary Data 4). Flow cytometric analysis of isolated hepatic macrophages confirmed these classifications. KC1 (CD45⁺Adgre1⁺CD11b⁻MHCII⁺Folr2⁺) dominated in healthy livers but declined in cirrhosis and rebounded during regression. In contrast, KC2 (CD45⁺Adgre1⁺CD11b⁻MHCII⁺Folr2⁻) and Momacs (CD45⁺Adgre1⁻CD11b⁺Ly6G⁻) expanded during fibrosis and declined upon regression (Fig. 4c–d, Supplementary Fig. 5c–d, and Supplementary Data 4). Due to the intracellular nature of Mki67, KC1 and KC3 could not be separated by flow cytometry and were analyzed together. Neutrophils (CD45⁺F4/80⁻CD11b⁺Ly6G⁺) followed a similar trend (Fig. 4d, bottom). Notably, in culture, Momacs displayed heterogeneous morphology with enlarged cytoplasm (Supplementary Fig. 5e), suggesting functional divergence among macrophage subsets. Gene set enrichment analysis (GSEA) revealed that while KC1 was associated with metabolic homeostasis, KC2, and Momacs were enriched in tumor necrosis factor superfamily signaling, leukocyte adhesion, and inflammatory pathways, KC3 enriched in nuclear division, mitotic nuclear division (Supplementary Fig. 5f). Cytokine profiling of primary sorted macrophages further showed that KC2 and Momacs secreted significantly higher levels of inflammatory mediators than KC1 (Supplementary Fig. 5g). Quantitative data showed that KC2 and Momacs displayed a highly inflammatory secretome (red and magenta boxes, respectively), including MCP-1, MIP1, IL-1β, TNF𝛼𝛼, CXCL16, CCL6, CCL12, CCL22, MMP9, OPN, SERPINE1, CD14, CHI3L1, CXCL10, IL10, GDF15, and MPO, whereas KC1 secreted non-inflammatory factors such as DPP4 (CD26, blue box) (Supplementary Fig. 5g-h). Interestingly, KC2 highly expressed matrix metalloproteinase signature (MMP-related genes and ADAM-related genes) and cathepsin signature (Supplementary Fig.5i). During regression, macrophages from recovered livers upregulated complement and degradative phagosome-associated phagocytic genes (Supplementary Fig. 5j–k).

**Fig. 4.**
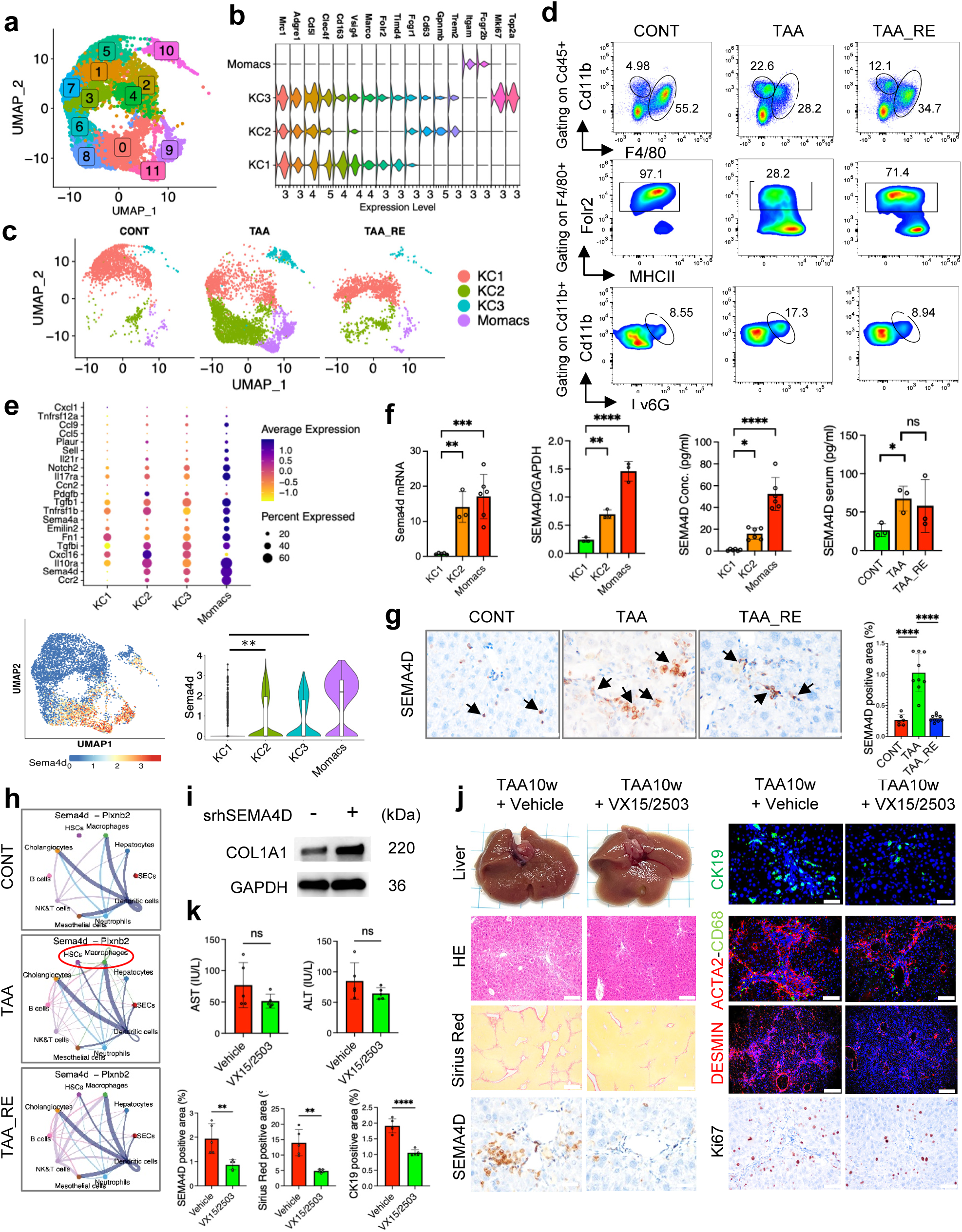
Sema4d⁺ Macrophages are key drivers of hepatic inflammation and fibrosis. **a** UMAP showing 12 subpopulations of macrophages, colored by cluster. **b** Stacked violin plot showing classified markers of macrophages. **c** UMAP showing macrophages from control, cirrhosis, and regression group, colored by specific macrophages subpopulations. **d** Gating strategy for identifying macrophages subpopulation in control, cirrhosis and in the regression phase. **e** Dotplot showing secretomes of macrophages subpopulations, colour by average expression (upper panel). UMAP showing expression of Sema4d alongside subpopulations (lower panel-left). Violin plot showing expression of Sema4d among KC1, KC2, Momacs and KC3 (lower panel-right) **f** Barplot shows expression of quantitative RT-PCR and immunoblotting of Sema4d in KC1, KC2, and Momacs (2-panel left). GAPDH was used as an internal control. Sema4d concentration in media of KC1, KC2, and Momacs (middle panel), and Sema4d concentration in serum of healthy, fibrosis, and regression group (right panel). **g** IHC showing expression of SEMA4D (black arrow) in livers section of control, fibrosis and regression group and its quantification. Scale bar 50 𝜇𝜇m. **h** Chord diagram showing Sema4d-Plxnb2 interaction among all cell types in healthy, fibrosis, and regressed livers. **i** Immunoblotting showing COL1A1 expression of HHSteCs treated with srhSema4D. GAPDH was used as internal control. **j** Representative images of livers, H&E staining; Sirius Red staining; IHC staining of SEMA4D, KI67; IF staining of CK19 (green) – DAPI (blue), DESMIN (red) – DAPI (blue); and double IF staining of ACTA2 (red) - CD68 (green) – DAPI (blue) of TAA 10 weeks mouse model treated with Vehicle or VX15/2503 at dose of 5 mg/kg tail vein injection twice a weeks within last 5 weeks (n = 5, each group). Scale bar 100 𝜇𝜇m (H&E, Sirus Red) and 20 𝜇𝜇m (SEMA4D, CK19, ACTA2-CD68, DESMIN, KI67) **k** Serum level of AST, and ALT (UI/l, upper panel). Quantification of positive area of SEMA4D, Sirius Red, and CK19 of TAA 10 weeks mouse model treated with Vehicle or VX15/2503 at dose of 5 mg/kg within last 5 weeks.. For **f-g-k**, Data represent the means ± SD from at least three separate experiments. *p-*values were analyzed by two-tailed Student’s t-test or one-way ANOVA followed by Tukey’s multiple comparison test. \**p* < 0.05, \*\**p* < 0.01, \*\*\**p* < 0.001, \*\*\*\**p* < 0.001.

Single-cell secretome profiling identified Sema4d as one of the most prominent cytokine transcripts in Momacs (Fig. 4e), and trajectory analysis implicated Sema4d as a key regulator of Momacs activation (Supplementary Fig. 6a–b). Sema4d expression was significantly elevated at the mRNA and protein levels in KC2 and Momacs (Fig. 4f, two left panels; Supplementary Fig. 6c), and in their conditioned media and serum (Fig. 4f, two right panels). Hepatic SEMA4D expression increased markedly in fibrotic livers and declined during regression (Fig. 4g), and was similarly elevated in other fibrosis models, including CCl₄ and bile duct ligation (Supplementary Fig. 6d). In human samples, SEMA4D was strongly expressed in advanced liver fibrosis (Supplementary Fig. 6e), was detectable in serum from MASLD patients, and correlated significantly with fibrosis stage (Supplementary Fig. 6f).

To explore how SEMA4D secreted from macrophages play a role in liver fibrosis, we analyzed cell–cell communication using CellChat. Sema4 signaling (Supplementary Fig.7a) markedly upregulated in liver fibrosis via its interaction with Plexin B1 and 2 (Plxnb1, 2) on HSCs (Fig. 4h). Although Plexin B1, 2 expression can be found at many cell types (Supplementary Fig.7b), MultiNicheNet analysis ranked Sema4d–Plxnb2 as the top macrophage–HSC interaction in TAA-induced fibrosis (Supplementary Fig. 7c). Plxnb2 expression in HSCs was induced under fibrotic conditions and absent in controls and reduced in regressed livers (Supplementary Fig. 8a). In vitro, PLXNB2 protein expression increased in primary mouse HSCs during spontaneous activation similar to ACTA2 and COL1A1 at days 4 and 7 compared to day 1 (Supplementary Fig. 8b). Treatment with soluble recombinant human SEMA4D (srhSEMA4D) protein dose dependently induced COL1A1 expression in LX2 (Supplementary Fig. 8c, Fig. 4i). In vivo blockade of SEMA4D using the humanized IgG4 monoclonal antibody against SEMA4D, VX15/2503, suppressed fibrosis progression. Mice treated with 0.5 or 1 mg/kg VX15/2503 during the final 3 weeks of a 6-week TAA model (Supplementary Fig. 8d–f), or with 5 mg/kg during the last 5 weeks of a 10-week advanced fibrosis model (Fig. 4j–k), showed reduced liver fibrosis as shown by Sirius Red staining, ductular reactions as shown by CK19 positivity, and HSC activation markers such as ACTA2 and Desmin and inflammation by CD68 staining, and cell proliferation Ki67.

Together, these findings reveal that SEMA4D derived from Momacs may be one of the key drivers of hepatic inflammation and fibrosis. Their targeted inhibition represents a promising therapeutic strategy for reversing liver fibrosis.

### Mmp7+ Cholangiocytes contributes to fibrosis progression via cytokine production

TAA treatment significantly increased the number of CK19⁺ CHOLs compared to healthy livers, which declined following two weeks of regression (Supplementary Fig. 9a). Reclustering identified six CHOL subclusters across control, cirrhotic, and regression groups (Fig. 5a–b), all of which express Krt7, Krt19, Epcam, Cd24a, Sox9, and Spp1 (Fig. 5d). CHOLs in control were mainly in subcluster 1, while those from cirrhotic livers were distributed across subclusters 0, 2, and 4, with subcluster 2 (Fig. 5c) enriched in pro-fibrotic and pro-inflammatory genes such as Lcn2, Scube3, Thbs1, Tgfb2, Pdgfrb, Cxcl16, Tnfrsf12a, Icam1, and Vcam1 (Fig. 5d), along with cytokine-mediated signaling pathways (Fig. 5e, left panel). CHOLs in recovered groups were mainly in subcluster 3 and 5 (Fig. 5c) enriched in negative regulation of programmed cell death and triglyceride metabolic process (Fig. 5e, middle and right panels, Supplementary Data 5). Trajectory analysis over pseudotime from TAA treatment to regression (Fig. 5f) revealed that Mmp7 was the top gene downregulated (Fig. 5g). The volcano plot analysis of differentially expressed genes also revealed that Mmp7 was markedly upregulated in cirrhotic livers compared to both control (Fig. 5h, upper panel) or recovery (Fig. 5h, lower panel), and confirmed at mRNA or serum (Fig. 5i). Serum level of Mmp7 showed signigicantly increased in all other mouse models of liver fibrosis induced by CCL4, or liver cholestasis induced by DDC, BDL, or high fat diet-induced steatohepatitis (Supplementary Fig. 9b). Transcription factor analysis revealed that CHOLs S2 upregulated inflammatory intracellular signals including Nfkb1-2, Nfkbiz, Rela, Relb, Ikbkb, Ikbke, Ikbkg, Traf1-5, Tnfaip1-3, Tradd, Fadd, Ripk1-4, Irak1-2 (Fig.5j). We further investigated the CHOL-specific secretomes by cytokine profiling from primary isolated and cultured Epcam⁺CHOLs (Fig. 5k) and found that cirrhotic CHOLs secreted various cytokines and chemokines (red boxes) including IGFBP6 and Osteoprotegerin (OPG, encoded by Tnfrsf11b gene) in comparison with other groups (Fig. 5k, circle boxes). Interestingly, the increased expression of MMP7, IGFBP6, and TNFRSF11B were also found in human PSC compared with healthy group via spatial transciptomics analysis from published database GSE245620 and GSE240029 (Supplementary Fig. 9c).

**Fig. 5.**
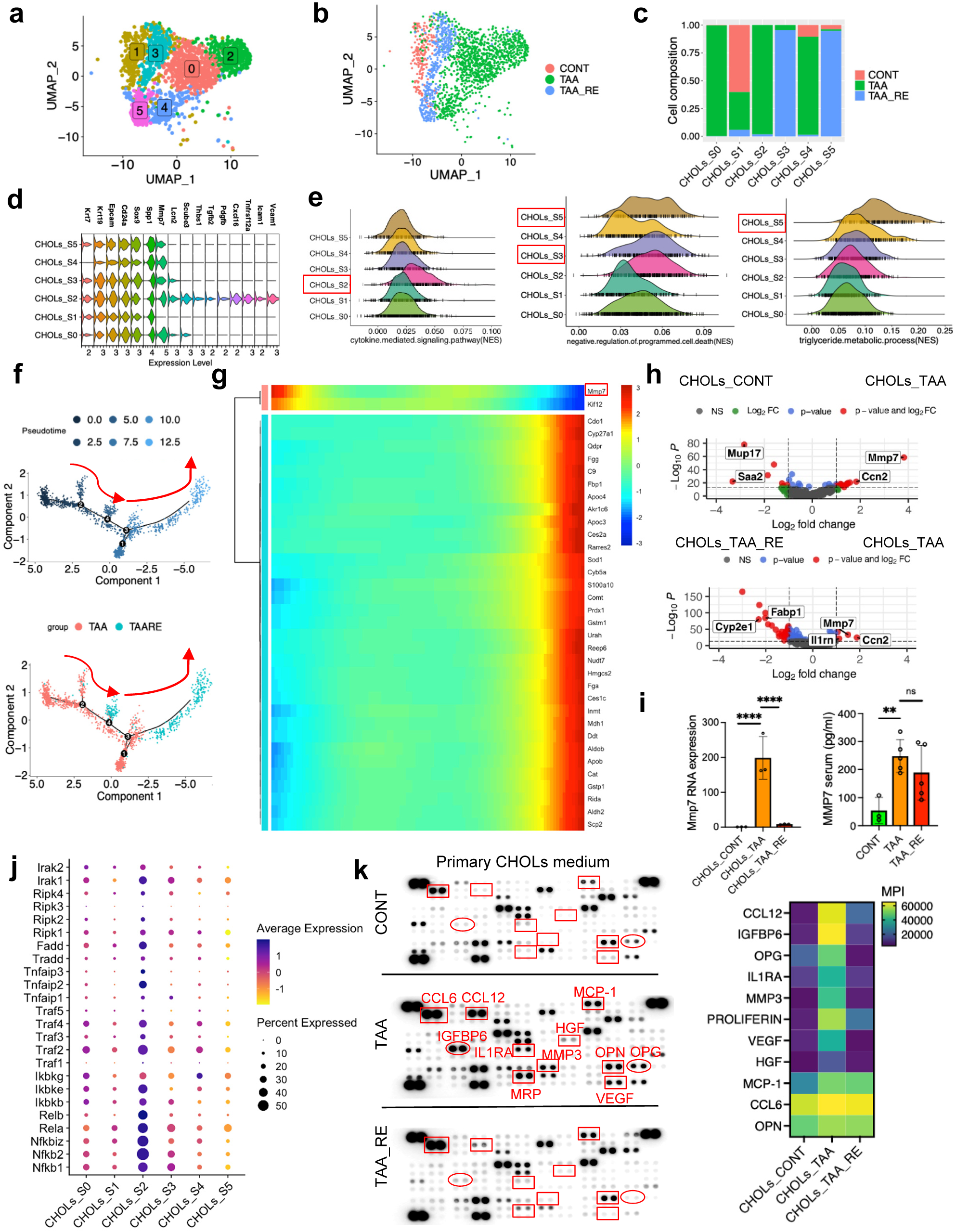
Mmp7+ Cholangiocytes contributes to fibrosis progression via cytokine production. **a** UMAP showing 6 subpopulations of CHOLs, colored by cluster. **b** UMAP showing CHOLs from control, cirrhosis, and regression group, colored by group. **c** Barplot shows group composition breakdown per cluster (fraction of total cell count per cluster), colored by group. **d** Stacked violin plot showing markers of cholangiocytes and mixed clusters. **e** Multi-ridge plots show the comparison of normalized enrichment score (NES) of selected gene set by scGSVA, colored by CHOLs subpopulation. **f, g** Pseudotime trajectory mapped to cholangiocytes UMAP coordinates. Direction of arrows indicates predicted cell state progression from fibrosis to regression phase (f) and heatmap (g) showing the dynamics of differentially expressed genes between two cell states. Genes (rows) and cells (columns) are ordered according to the pseudo-time trajectory. **h** Volcano plot showing differential gene expression of cholangiocytes in control and fibrosis group (upper panel), regressed and fibrosis group (lower panel). Gene upregulated or downregulated by more than 2-fold and significant to p < 0.05 is shown in red. **i** Quantification of Mmp7 expression by qRT-PCR in CHOLs isolated from control, fibrosis and regression group (left panel). Gapdh was used as internal control. Quantification of Mmp7 concentration in serum of control, cirrhosis, and regression group (right panel). Data represent the means ± SD, *p-*values were analyzed by one-way ANOVA followed by Tukey’s multiple comparison test. \**p* < 0.05, \*\**p* < 0.01, \*\*\**p* < 0.001, \*\*\*\**p* < 0.001. **j** Dot plot showing inflammatory-related genes of CHOLs cells subpopulation, colored by gene expression. The size of the dot corresponds to the percentage of cells expressing the gene in each CHOLs subpopulation. **k** Cytokine array of soluble factors in cultured media from cholangiocytes isolated from control, cirrhosis, and in the regression group, the soluble factors with predominant change were marked with red boxes or circles (left panel). Heatmap showing the soluble factors expression of each group by calculating mean pixel density (MPI), colored by MPI (right panel).

Collectively, CHOLs from regressed livers showed enrichment in the negative regulation of programmed cell death and triglyceride metabolic process while those from cirrhotic livers secreted inflammatory cytokines.

### LMCD1+ HSCs promotes fibrosis progression via modulating AKT-4EBP1

To examine the heterogeneity of HSCs across healthy, cirrhotic, and regressed livers, we subclustered HSCs into nine subsets (S0–S8; Fig. 6a–c), using a validated set of murine HSC markers^8^. In healthy livers, HSCs were predominantly found in subclusters S2, S3, and S8, expressing Reln, Tnxb, Colec10, Colec11, Fga, Fgb, and Vipr1, and enriched in pathways related to cell–cell junction assembly (S2), fibrinolysis and plasminogen activation (S3), and peptide hormone response and lipid metabolism regulation (S8) (Fig. 6c-e). In contrast, cirrhotic HSCs were highly represented in subclusters S0 and S4, characterized by elevated expression of canonical fibrogenic genes (Col1a1, Col1a2, Col3a1, Eln, Lox, Col15a1). S0 was enriched in extracellular matrix (ECM) organization, while S4 upregulated genes involved in integrin-mediated signaling, including Lama1, Lamb1, Steap4, and Notch3 (Fig. 6c–e; Supplementary Data 6). In regressed livers, subclusters S5 and S6 downregulated fibrogenic markers but expressed quiescence-related genes such as Cadm3, Agtr1a, Ehd3, Tmem56, Alcam, Emp1, and Tcf21, with enrichment in vasculogenesis, blood vessel morphogenesis, and ECM organization (Fig. 6d–e). Subclusters S1 and S7 included HSCs from all three conditions and were enriched for physiological HSC functions, including triglyceride metabolism (S1) and elastic fiber assembly (S7). Trajectory analysis revealed a shift from activated to inactivated phenotypes during regression, with upregulation of the inactivation gene module and suppression of the activation module (Fig. 6f–g). Among the genes modulated during this transition, members of the LIM domain protein family including Fhl2, Fhl3, Pdlim2, Pdlim7, Limk2, Lima1, and Lmcd1 were significantly upregulated in fibrotic HSCs but downregulated during regression (Fig. 6h). These findings were validated in primary murine HSCs, where expression of these genes increased at days 4 and 7 of culture compared to day 1 at both mRNA (Supplementary Fig. 10a) and protein levels (Fig. 6i). In human HHSteCs, both spontaneous activation (by shifting culture condition from with supplement (S+) to without supplement (S-) and TGFβ1 (2 ng/mL) stimulation significantly upregulated LMCD1, LIMK2, PDLIM5, and PDLIM7 at transcript and protein levels (Supplementary Fig. 10b–d). In a public single-cell dataset from healthy and NASH patients^36^, LIM domain gene expression, including LMCD1, was elevated in HSCs from fibrotic livers (GSE212837; Supplementary Fig. 11a–d).

**Fig. 6.**
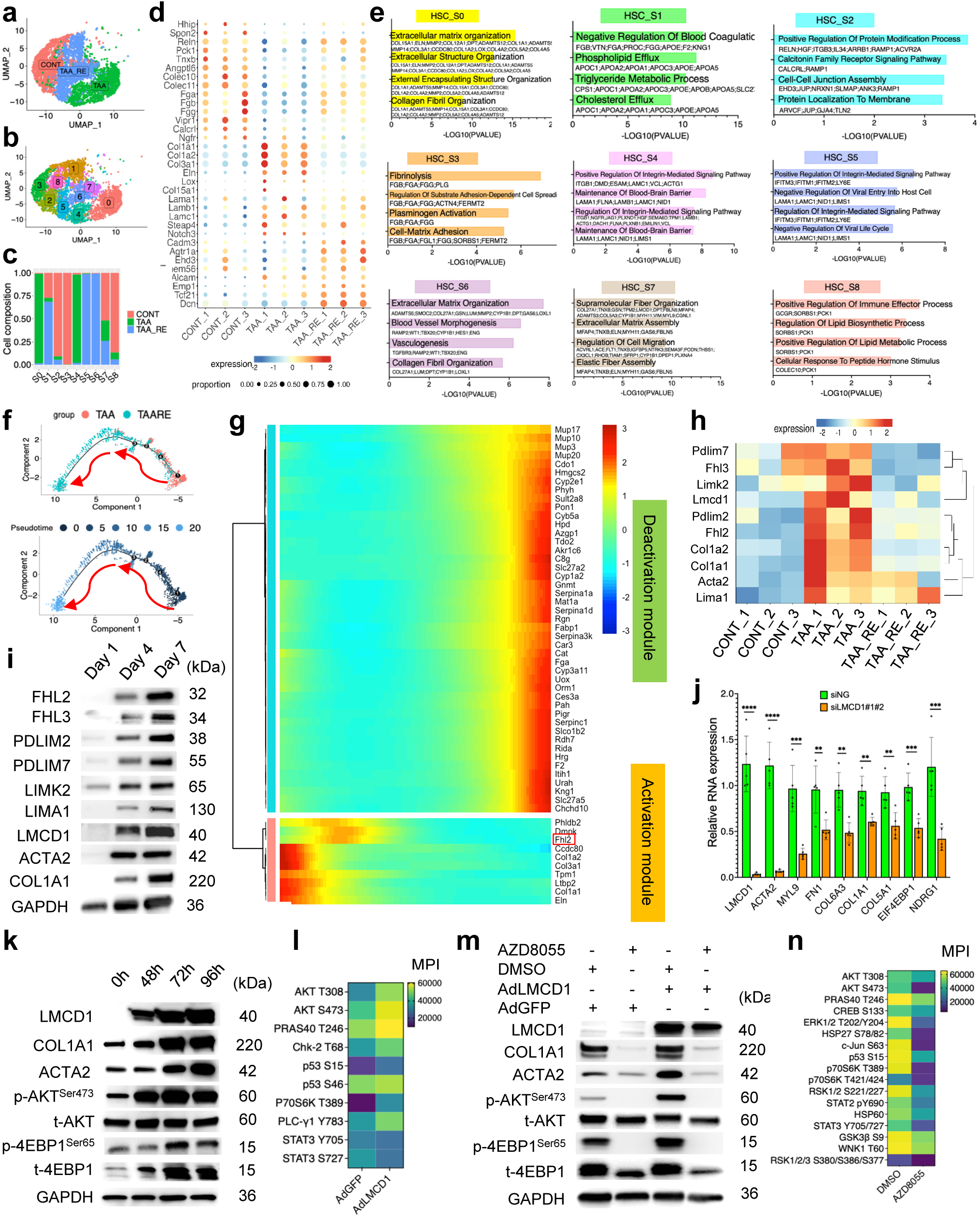
LMCD1+ HSCs promoted fibrosis progression via modulating AKT-4EBP1. **a** UMAP showing HSCs from control, cirrhosis, and regression group, colored by group. **b** UMAP showing 9 subpopulations of HSCs, colored by cluster. **c** Barplot shows group composition breakdown per cluster (fraction of total cell count per cluster), colored by group. **d** Dot plot showing group-specific gene expression in HSCs, colored by expression **e** Barplot showing top 4 GO pathways analyzed by EnrichR of each HSCs subclusters, ordered by -log10(p-value) of each pathway. **f, g** Trajectory reconstruction of all HSCs reveals one branch of cell fate (g). Cells are colored by pseudo-time and group, respectively. Heatmap (h) showing the dynamics of differentially expressed genes between two cell states. Genes (rows) and cells (columns) are ordered according to the pseudo-time trajectory. **h** Heatmap showing expression of LIM family related genes among group, colored by gene expression **i** Immunoblotting showing the expression of LIM family protein of mHSCs at day 1, day 4, day 7 after isolation. **j** Quantitative RT-PCR showing RNA expression of LMCD1, ACTA2, MYL9, FN1, COL6A3, COL1A1, COL5A1, EIF4EBP1, NDGR1 in HHSteCs under silencing negative control (siNG) or LMCD1 (siLMCD1). 18S was used as internal control. Data represent the means ± SD. *p*-values were analyzed by two-tailed Student’s t-test. \**p* < 0.05, \*\**p* < 0.01, \*\*\**p* < 0.001, \*\*\*\**p* < 0.0001. **k** Immunoblotting showing protein expression of LMCD1, COL1A1, ACTA2, p-AKT^Ser473^, t-AKT, p-4EBP1^Ser65^, t-4EBP1 in HHSteCs under overexpression of LMCD1 time-dependently. **l** Heatmap showing quantitative expression of phosphoproteins expression of overexpression of GFP (AdGFP) or LMCD1 (AdLMCD1) by phosphokinase array, colored by mean pixel density (MPI). **m** Immunoblotting showing protein expression of LMCD1, COL1A1, ACTA2, p-AKT^Ser473^, t-AKT, p-4EBP1^Ser65^, t-4EBP1 in HHSteCs under AdGFP or AdLMCD1 treated with and without AZD8055 1 𝜇𝜇M. **n** Heatmap showing quantitative expression of phosphoproteins expression by phosphokinase array of HHSteCs under TGF𝛽𝛽1 treatment (2 ng/ml) treated with DMSO or AZD8055 1 𝜇𝜇M, colored by MPI. For **i, k, m.** GAPDH was used as an internal control.

We next focused on LMCD1 as a potential fibrosis marker. In our MASLD cohort, LMCD1 was highly expressed in fibrotic septa (Supplementary Fig. 12a) and positively correlated with fibrosis stage (AUROC = 0.92, p = 0.006; Supplementary Fig. 12b). Immunofluorescence confirmed LMCD1 colocalization with ACTA2 and Vimentin, markers of activated HSCs (Supplementary Fig. 12c). LMCD1 was also expressed in cancer-associated fibroblasts in HCC (Supplementary Fig. 12d) and prominently localized along fibrotic septa in HCV-HCC tissues (Supplementary Fig. 12e–f). To assess functional relevance, we silenced LMCD1 in HHSteCs using two siRNAs, achieving 95% (mixture of siLMCD1#1 and siLMCD1#2) knockdown (Fig. 6j) resulting in suppressed COL1A1, COL5A1, COL6A3, FN1, ACTA2, MYL9, EIF4EBP1, and NDGR1 expression at transcript levels (Fig. 6j). Conversely, overexpression of LMCD1 by adenoviral transduction (AdLMCD1) induced HSCs activation via increased expression of fibrosis related genes (Supplementary Fig. 13a), and collagen synthesis (Fig. 6k), and enhanced phosphorylation of AKT and 4EBP1, all known pro-fibrotic signaling molecules^37^. A phosphokinase array further confirmed reciprocal regulation of these pathways in LMCD1-overexpressed cells, particularly within the AKT signal (Fig. 6l; Supplementary Fig. 13b). Pharmacological inhibition of this axis using AZD8055 in LMCD1-overexpressing HHSteCs suppressed COL1A1, ACTA2, and LMCD1 expression, and downregulated 16 out of 37 phosphoproteins including CREB, c-JUN, TP53, WNK1, GSK3B, ERK1/2, P70S6K, and RSK1/2/3 (Fig. 6m–n; Supplementary Fig. 13c–d). In culture condition, treatement of AZD8055 also mitigated spontaneous and TGFβ1–induced HSC activation in both murine (Supplementary Fig. 14a–b) and human (Supplementary Fig. 14c-f) in a dose-dependent manner. Moreover, in vivo application of AZD8055 therapy by i.p. injection twice week for the last 4 weeks of TAA-8-week treatment (Supplementary Fig. 15a-d) demonstrated its role in suppression of fibrosis development by reducing collagen formation indicated by Sirius-red staining, HSC activation indicated by ACTA2 and Desmin staining, and also reduced inflammatory indicated by CD68 and SEMA4D staining (Supplementary Fig. 15b-d). Collectively, these results identify LMCD1 as a novel activation marker of HSCs, with functional relevance in liver fibrogenesis via the AKT–4EBP1 pathway. Targeting LMCD1 represents a promising therapeutic strategy to suppress HSC activation and halt fibrosis progression.

### A subset of immune cells may contributes to liver fibrosis regression

Although immune cells are known to play roles in liver fibrosis progression, their contribution to regression remains underexplored. We investigated the roles of NK&T cells, neutrophils, and B cells in fibrosis resolution. Clustering of NK&T cells from control, TAA-treated, and regression-phase livers revealed eight subclusters (Fig. 7a–c). Cluster 0, 2, 3, 4, 5, 6, 7, and 8 which expressed Cd3d, Cd3e, and Cd3g were annotated as T cells. Cluster 1 expressing Gzma, and Ncr1 was annotated as NK cells. Notably, subclusters S5 and S6 were predominantly derived from regressed livers. S6, expressing Cd3d, Cd3e, Cd3g, Cd4, Cd8a, Sell, and S1pr1 but lacking Cd44, was annotated as naïve T cells^12^, whereas S5 positive for Cd8 and Cd44, but negative for Sell and S1pr1, represented CD8⁺ memory T cells^12^ (Fig. 7d). Gene set enrichment analysis (GSEA) showed that S6 upregulated pathways related to T cell activation, endothelial barrier establishment, and cytokine response, while S5 enriched antigen receptor and NK cell chemotaxis signaling (Fig. 7e). Cytokine-associated genes such as Il17ra, Il6ra, Il7r, Cxcr6, Ifngr1, Ccl5, and Gzmk were upregulated in S5 and S6, paralleling T cell subcluster S2 (Fig. 7f). Transcription factors Mef2a, Mef2b, and Mef2c, involved in mitochondrial fatty acid oxidation and T cell differentiation^38, 39^, were also upregulated S6 (Fig. 7g). Cell–cell interaction analysis revealed upregulated CCL and CXCL network communication in recovered livers compared to fibrotic ones (Fig. 7h). Flow cytometry of primary lymphocytes showed restoration fraction of CD4⁺, CD8⁺, and NK1.1⁺ cells, which indicated naiive T and Cd8 memory T cells, while Foxp3⁺ regulatory T cells remained stable (Fig. 7i).

**Fig. 7.**
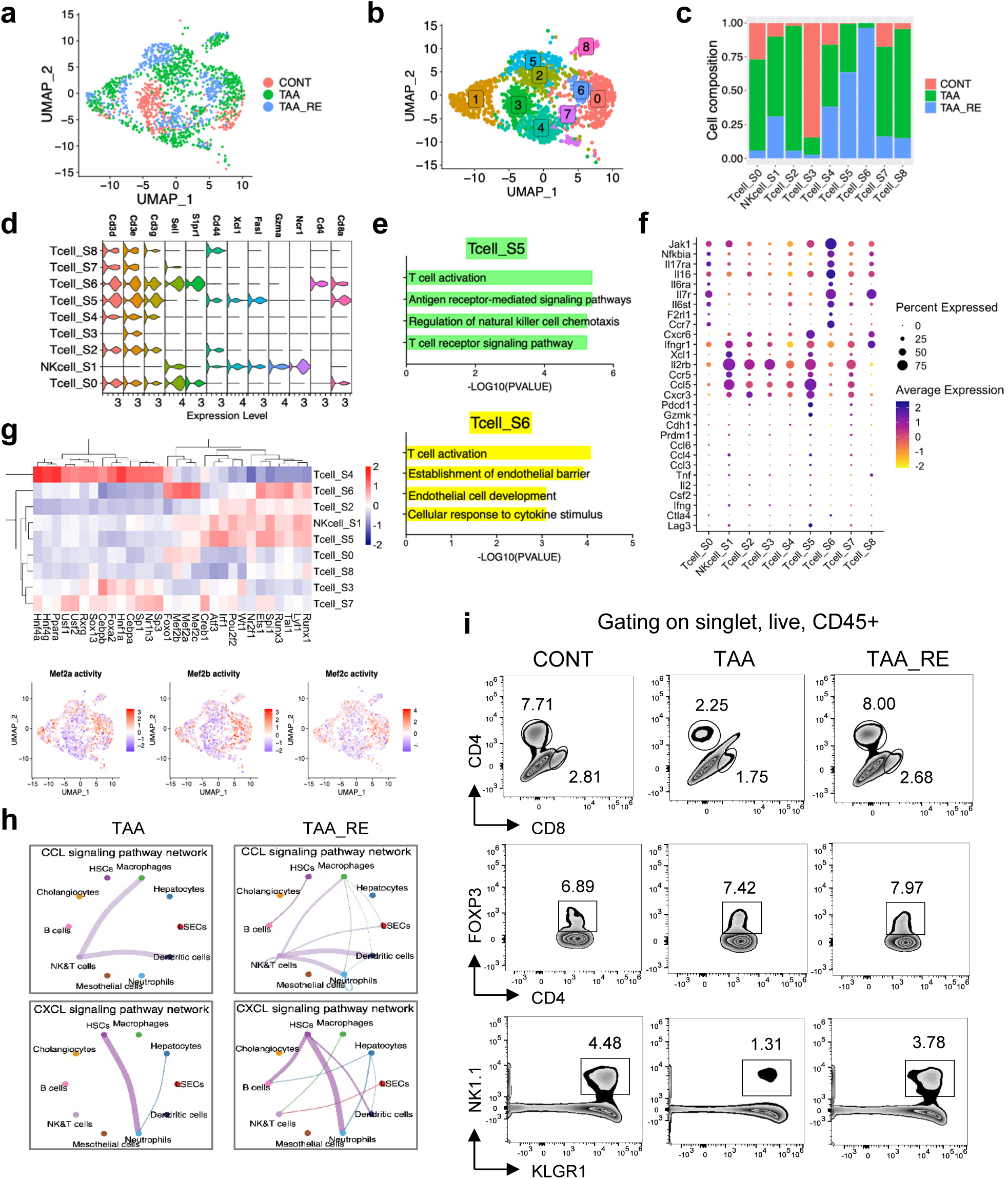
A subset of immune cell may contribute to fibrosis regression. **a** UMAP showing NK&T cells from control, cirrhosis, and regression group, colored by group. **b** UMAP showing 9 subpopulations of NK&T cells, colored by cluster. **c** Barplot shows group composition breakdown per cluster (fraction of total cell count per cluster), colored by group. **d** Stacked violin plot showing expression markers of NK&T cells. **e** Barplot showing top 4 GO gene set enrichment analysis of NK&T subclusters S5, S6; compared by -log10(PVAL) of each pathway. **f** Dot plot showing cytokine-related genes of NK&T cells subpopulation, colored by gene expression. The size of the dot corresponds to the percentage of cells expressing the gene in each cluster. **g** Transcription factor analysis by Dorothea, colored by average expression (upper panel); Transcription factor activity of Mef2a, Mef2b, Mef2c in NT&T cells subpopulation, colored by gene expression. **h** Interaction showing CCL (upper panel) and CXCL (lower panel) signaling pathways network interaction among all cell types in fibrosis and regressed livers **i** Gating strategy for NK&T cell subpopulations in control, cirrhosis, and in the regression group with CD45+CD4+CD8+FOXP3+NK1.1+KLRG1+.

B cell analysis identified six subclusters (S0–S5; Supplementary Fig. 15a–c), with S0 primarily derived from regressed livers. GSEA showed enrichment in pathways related to cytoskeletal organization, intracellular transport, BCR signaling, glucose response, and mRNA splicing (Supplementary Fig. 15d, Supplementary Data 7). Transcription factor analysis revealed upregulation of Ets1 and Spi1 in subcluster S0 of recovered B cells, essential for B cell development and BCR signaling^40, 41^ (Supplementary Fig. 15e–f) may reflect the recovery process.

NEUs clustered into four subpopulations (S0–S3; Supplementary Fig. 16a–c), with S3 comprising cells from all stages, but enriched in recovered livers. GSEA revealed activation of regulatory pathways related to Toll-like receptor signaling, IL-12, and ROS metabolism (Supplementary Fig. 16d). Stat3, Stat5b, Spi1, and Cebpa were the most upregulated transcription factors in S3 (Supplementary Fig. 16e–f) of NEUs from recovered group.

Collectively, although we have not investigated the detail functioning of each cluster of immune cells here, but our data revealed that in immune subpopulations recovered NK&T cells showed upregulated pathways related to T cell activation, endothelial barrier establishment, and cytokine response while recovered B cells demonstrated enrichement of pathways related to cytoskeletal organization, intracellular transport, BCR signaling, glucose response, and mRNA splicing. Finally, recovered neutrophils showed activation of regulatory pathways related to Toll-like receptor signaling, IL-12, and ROS metabolism regain their regulatory and protective functions during liver fibrosis regression.

### Cell–cell communication networks driving fibrosis regression

To characterize intercellular interactions associated with fibrosis progression and regression, we performed a global ligand–receptor network analysis using CellChat. We focused on interactions from all other cell types to HSCs. Bubble plot analysis comparing communication probability of ligand-receptors interaction between TAA and control showed that in healthy condition, the ligand-receptor reactions (red font) in order from strongest one coming from HSCs, mesothelial cells, CHOLs, LSECs, HEPs to HSCs, while other cell types including MACs, NEUs, NK&T, B cells, DCs showing no significant interactions with HSCs (Fig. 8a). In contrast, TAA treated HSCs received strong incoming signals (interactions in blue font) from all other cell types with the strongest one from autocrine interaction (Fig. 8a). Figure 8b showed that interaction strengths were diminished during regression phase (interactions in blue font).

**Fig. 8.**
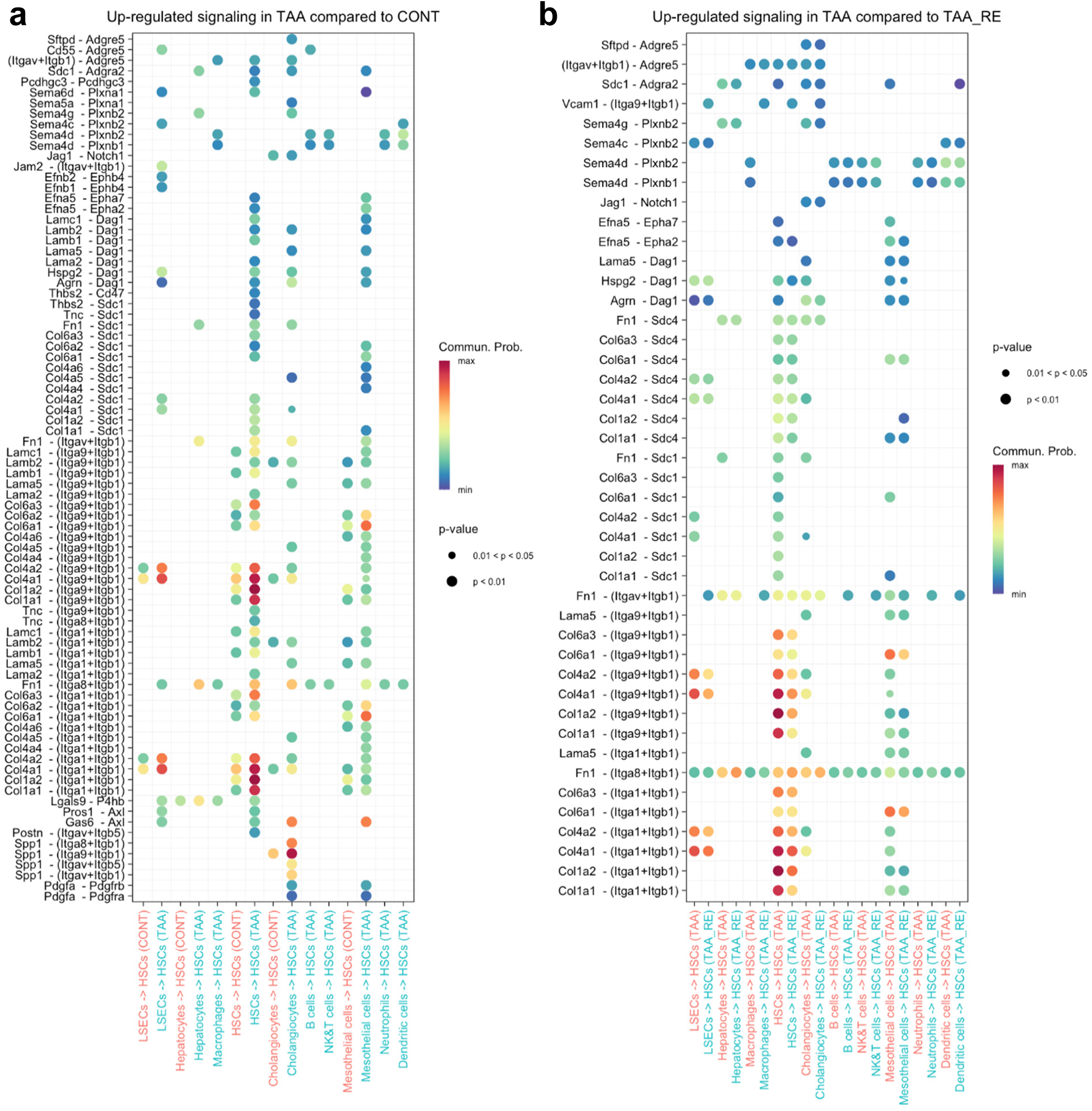
Cell–cell communication networks in fibrosis progression and regression. **a** Bubble plot showing upregulated ligand-receptor pairs interaction from 10 major cell types to HSCs in fibrosis progression compared to healthy control group, colored by communication probability. **b** Bubble plot showing upregulated ligand-receptor pairs interaction from 10 major cell types to HSCs in fibrosis progression compared to regression group, colored by communication probability. For **(a)** and **(b)** bubble color and size represent the level of communication probability and p-value, respectively, computed using CellChat.

To further interrogate altered intercellular signaling, CellChat was used to plot predicted relative information flow of significantly altered pathways in TAA vs control (Supplementary Fig. 18a) and in TAA vs recovered liver cells (Supplementary Fig. 18b). The pathways that were significantly regulated only in healthy liver but not in TAA treated liver cells such as FGF, IL1, CEACAM (Supplementary Fig. 18a), were also found in recovered liver cells (black arrows).

Interestingly, CellChat showed that the majority of outgoing signaling patterns among all groups are from HSCs, increased in fibrosis progression (Supplementary Fig. 19) meanwhile LSEC population demonstrating significant outgoing signaling in recovered cells. Exploring incoming signaling patterns, the results showed that macrophages became the main receiver cells in all groups (Supplementary Fig. 20), hepatocytes is the second receiver cells in healthy livers, and dendritic cells, neutrophils and LSECs are those contributing to receive the signaling in recovered cells.

In summary, using fixed single-cell RNA sequencing, we constructed a high-resolution cellular atlas and their interaction across healthy, fibrotic, and regressed mouse livers. Our integrative analyses identified key subpopulations and their interactions that orchestrate both progression and resolution of liver fibrosis. Notably, we pinpoint SEMA4D (from Momacs) and LMCD1 (from HSCs) as novel regulators of fibrosis, representing promising therapeutic targets.

## Discussion

In this study, we applied single-cell fixed RNA sequencing (FLEX) to investigate the cellular and molecular changes during liver fibrosis regression. By profiling over 38,000 cells from control, TAA-induced cirrhotic, and regression-phase mouse livers, we generated a comprehensive cell atlas of these conditions. Importantly, we identified novel subpopulations and molecular mediators, the macrophage-derived factor SEMA4D and the HSC-specific transcriptional regulator LMCD1 as promising targets for antifibrotic therapy.

### Single-cell Fixed RNAseq: A New Application in Liver Fibrosis Research

Our study is one of the first to use FLEX, 10x Genomics in liver fibrosis models, demonstrating the feasibility and advantages of this approach in preserving in vivo transcriptomic profiles during sample preparation. Traditional single-cell RNAseq methods using fresh or enzymatically dissociated cells can introduce stress-induced artifacts^42^, especially in fragile hepatic cell types like LSECs and HEPs^21^. By contrast, FLEX allows immediate tissue fixation, minimizing transcriptional changes during processing and enabling the capture of a broader range of cell states, including those from inflamed or necrotic regions often lost during standard dissociation^21^.

This technical advancement facilitated the identification of rare but functionally important populations in both fibrotic and regressed livers, including recoHEPs, restorative LSECs, pro-resolution macrophages, and inactivated HSCs. Furthermore, the preservation of cell morphology and integrity also enabled successful integration with flow cytometry and spatial analyses.

### Cellular Mechanisms of Fibrosis Regression: New Insights Across Cell Types

Our single-cell profiling uncovered significant remodeling across multiple cell types during fibrosis regression. In HEPs, we identified a peri-central subpopulation (recoHEPs) enriched in antioxidant and xenobiotic metabolism genes (e.g., Cyp2e1, Txn1) that reappeared during regression. Previous work by Ben-Moshe et al.^43^ characterized hepatocyte zonation changes during injury, but our study uniquely shows the regeneration of functionally competent peri-central hepatocytes during regression, indicating not just structural but functional restoration.

These recoHEPs and regressed HSCs higly expressed Rarres2 which we propose plays a role in modulating HSC activity. Our study showed that induction of RARRES2 in HHsteCs using retinoic acid receptor agonist reduced activation of HSCs. Recent study directly investigates the effects of chemerin isoforms (huChem-157, huChem-156, huChem-155) on human HSCs, using the LX-2 cell line and primary HSCs. It found that huChem-156 up-regulates IL-6, IL-8, and galectin-3 in LX-2 cells, indicating its role in promoting inflammatory and fibrotic responses^44^. Sebastian et al. reports that CCRL2, a receptor for chemerin (Rarres2), is expressed in HSCs, LSECs, and Kupffer cells. CCRL2 mRNA levels in HSCs are induced by TNF and positively correlate with inflammation, fibrosis, and NASH scores in 85 NAFLD patients compared to 33 controls. It also correlates with CMKLR1 mRNA, suggesting a coordinated chemerin receptor system in HSCs^45^. These evidences leans toward chemerin interacting with HSC receptors like CMKLR1 and CCRL2, supporting its role in liver fibrosis, and more research is needed to clarify the exact mechanisms and therapeutic potential.

In LSECs, regression was marked by the re-emergence of fenestrated subtypes expressing matrix-degrading enzymes such as Mmp14 and Ctsl, and antioxidant genes including Pon1 and Tcn2. These observations align partially with Kantari et al.^46^, who described myeloid cell– derived VEGF as a critical regulator of fibrosis resolution via Mmp2/14 secreted by LSECs. Our data further extend these findings by showing that recovered LSECs possess pro-resolving transcriptional programs and morphological features (e.g., restored fenestration), reinforcing their active role in scar remodeling.

The macrophage compartment also exhibited dynamic changes. We identified three major populations-KC1, KC2, and Momacs-with distinct roles in fibrosis progression and resolution. In agreement with Ramachandran et al.^15^ and Krenkel et al.^47^, we found that Momacs dominate during fibrogenesis. However, our data uniquely highlight a shift toward anti-inflammatory, phagocytic KC1 cells during regression, characterized by enhanced expression of complement and antioxidant genes. Importantly, SEMA4D’s role in liver fibrosis is further elucidated by its ability to enhance HSC activation through Plxnb2 signaling, with knockout studies demonstrating reduced fibrosis via suppression of AOX1 in retinol metabolism and modulation of T cell subsets^48^. The therapeutic promise of SEMA4D blockade is evident from preclinical trials where VX15/2503, a monoclonal antibody, significantly reduced fibrosis, inflammation, and HSC activation in TAA-induced mouse models, positioning it as a targeted anti-fibrotic strategy. In human MASLD, elevated SEMA4D serum levels correlate with advanced fibrosis stages, suggesting its potential as a biomarker for disease severity^49^. SEMA4D’s profibrotic effects are not limited to toxic models but extend to parasitic liver fibrosis, such as schistosomiasis, where its inhibition by Sja-miR-71a mitigates fibrosis, indicating a broad role in hepatic fibrogenesis^50^.These findings highlight the need for further research into SEMA4D’s downstream pathways and its integration into combination therapies to enhance fibrosis resolution.

Cholangiocytes also showed plasticity during regression. We identified a fibrogenic subpopulation (CHOL_S2) during TAA-induced fibrosis marked by Mmp7, Tgfb2, and Pdgfrb expression, which declined in the regression phase (Fig.5d). Our findings are consistent with those of Pepe-Mooney et al.^51^, who described biliary expansion and remodeling in liver injury. However, our study adds new insights by linking cholangiocyte-derived cytokines, emphasizing their role in promoting liver fibrosis.

HSC analysis revealed multiple activation states, including two regression-specific clusters expressing inactivation-associated genes (Cadm3, Tcf21) and reduced collagen genes (Col1a1, Col3a1). Notably, we identified LMCD1, a LIM-domain containing protein, as a novel marker and act as transcription factor of HSCs activation. LMCD1 was highly expressed in fibrotic HSCs and downregulated in regression. Functional studies in HHSteCs confirmed its role in collagen synthesis via the AKT–4EBP1 pathway. While prior work has linked LMCD1 with scleroderma-associated lung fibrosis^52^ and kidney fibrosis^53^, LMCD1 has not previously been implicated in liver diseases, highlighting its novelty and potential clinical relevance. LMCD1’s role in fibrosis is not limited to the liver, as prior studies have established its involvement in other organs, providing a comparative framework for its hepatic functions. In renal fibrosis, LMCD1 promotes tubulointerstitial inflammation by activating NFATc1-mediated NLRP3 pathways, leading to increased collagen deposition in a unilateral ureteral obstruction (UUO) mouse model^54^. Silencing LMCD1 in this model reduced expression of TGFβ1, fibronectin, and collagens, suggesting a mechanism that may parallel its role in liver HSCs. Similarly, in scleroderma-associated interstitial lung disease (SSc-ILD), LMCD1 is upregulated in lung fibroblasts, contributing to their profibrotic phenotype by enhancing ECM production^52^. These studies indicate a conserved fibrogenic role for LMCD1 across tissues, likely mediated by its LIM domains, which facilitate protein-protein interactions in signaling pathways like ERK, NFATc1, and AKT. Our observation that LMCD1 is also expressed in HCC-associated fibroblasts (Supplementary Fig. 12d–e) suggests it may contribute to fibrotic microenvironments in liver cancer, a novel finding that warrants further exploration.

Our analysis also revealed dynamic immune cell remodeling during fibrosis regression, including subsets of NK&T cells, B cells, and neutrophils with restored functional profiles. A unique cluster of naïve T cells (Cd3d⁺, Cd4⁺, Cd8a⁺, Sell⁺, S1pr1⁺) appeared exclusively during the regression phase, enriched in pathways related to T cell activation and endothelial barrier function. Validated flow cytometry analysis confirming the regain fraction of NK, and TCD4/CD8 cells in recovered phase reported partial restoration of immune surveillance during fibrosis resolution. These data were consistent with clinical observations showed that a subset of CD56^dim^KLRG1+ NK cells decreased in the peripheral blood and liver of patients with chronic HBV infection at advanced stages of fibrosis^55^. These NK cells exhibited high level of IFN-γ expression and TRAIL-dependent cytotoxicity against HSCs, which could be further enhanced by IFN-γ and CD44 stimulations^55^. Our results also indicate a more distinct shift toward anti-inflammatory phenotypes—particularly in neutrophils and B cells, which upregulated regulators of ROS metabolism and B cell receptor signaling, respectively. These interesting results suggest the need for further studies examining immune cell plasticity across diverse etiologies and regression stages.

### Therapeutic Implications of SEMA4D and LMCD1 Targeting

The therapeutic potential of SEMA4D blockade using the humanized IgG4 monoclonal antibody VX15/2503 has shown promising results in suppressing liver fibrosis, as demonstrated in the TAA-induced mouse model. In vivo studies revealed that VX15/2503, administered at 0.5 or 1 mg/kg in a 6-week model or 5 mg/kg in a 10-week advanced fibrosis model, significantly reduced fibrosis, as evidenced by decreased Sirius Red staining, ductular reactions (CK19 positivity), HSC activation (ACTA2, Desmin), inflammation (CD68), and cell proliferation (Ki67). This aligns with SEMA4D’s role as a key driver of inflammation and fibrosis, particularly through Sema4d–Plxnb2 interactions between Momacs and HSCs. The blockade of SEMA4D disrupts this pro-fibrotic signaling, offering a targeted anti-inflammatory strategy to halt disease progression. Emerging anti-inflammatory therapies, such as those targeting TNF^56^ or IL-1β^57^, share similar goals of mitigating macrophage-driven inflammation, but SEMA4D blockade provides a more specific approach by addressing macrophage–HSC crosstalk. Furthermore, SEMA4D correlated with fibrosis stage in MASLD serum suggests to establish it as a biomarker for predicting severity of fibrosis (Supplementary Fig. 6f). The success of VX15/2503 in preclinical models parallels other monoclonal antibody therapies, such as anti-TGFβ, which also target fibrogenic pathways^58^. Combining SEMA4D blockade with existing anti-fibrotic agents, like nintedanib, could enhance efficacy by addressing multiple pathways^59^. Future clinical trials are warranted to evaluate VX15/2503’s safety and efficacy in human liver fibrosis, potentially positioning it as a novel therapeutic in the anti-inflammatory arsenal.These findings are in line with ongoing clinical trials testing anti-SEMA4D antibodies in Huntington’ diseases^60^, suggesting potential for rapid translation to liver fibrosis.

LMCD1, a nuclear co-regulator, appears to act cell-autonomously within HSCs. LMCD1 has emerged as a novel activation marker for HSCs, with significant upregulation in fibrotic livers and validation in human MASLD and HCC (Supplementary Fig. 12a–c). In single-cell datasets from MASLD patients, LMCD1 expression was elevated in fibrotic HSCs, correlating strongly with fibrosis stage (AUROC = 0.92, p = 0.006), highlighting its potential as a biomarker (Supplementary Fig. 12b). Functional studies demonstrated that LMCD1 regulates HSC activation through the AKT–4EBP1 signaling pathway, with its overexpression enhancing collagen production and silencing reducing COL1A1 and ACTA2 expression (Fig. 6j–k). This aligns with prior research on AKT signaling in HSC fibrogenesis, where its activation promotes extracellular matrix (ECM) deposition^37^. AZD8055, an AKT-4EBP1 inhibitor, showed promise as a therapeutic agent by suppressing LMCD1-driven HSC activation in both in vitro human HSC cultures and in vivo TAA-induced mouse models (Fig. 6m–n; Supplementary Fig. 15a-d). These findings are consistent with mTOR inhibitors’ anti-fibrotic effects in liver disease models, which target AKT-dependent pathways^61^. This suggests that LMCD1 may be both a biomarker and a driver of HSC-mediated fibrosis, with potential for targeted intervention through transcriptional or post-transcriptional approaches.

### Limitations

Several limitations should be acknowledged. First, our study used a chemical-induced fibrosis model (TAA), which may not fully represent the heterogeneity of human liver disease (e.g., MASH, viral hepatitis). Although we validated key findings using public human datasets, future studies should confirm SEMA4D and LMCD1 relevance in broader patient cohorts. Second, while our FLEX protocol minimized dissociation artifacts, it lacks the temporal resolution of live-cell tracking or functional assays in situ. Combining FLEX with spatial transcriptomics or lineage tracing could strengthen causal interpretations. Third, while in vitro assays demonstrated functional roles for SEMA4D and LMCD1, in vivo genetic knockdown or conditional knockout models would provide more definitive evidence.

In conclusion, our study provides a comprehensive, single-cell transcriptomic map of liver fibrosis progression and regression. By leveraging fixed RNAseq technology, we identified critical cell populations and signaling pathways that mediate fibrosis resolution. Notably, we uncovered SEMA4D as a macrophage-derived ligand promoting HSC activation and LMCD1 as a novel transcriptional driver of fibrogenesis. Both factors represent promising therapeutic targets for liver fibrosis. These insights offer a deeper understanding of the cell–cell interactions and molecular reprogramming that govern liver regeneration, opening avenues for precision antifibrotic therapy.

## Methods

### Human tissue specimens

Patients with biopsy-proven metabolic dysfunction-associated steatotic liver disease (MASLD), who were recruited at Osaka Metropolitan University Hospital (Osaka, Japan). Liver biopsy specimens were obtained using a 16-gause MAX-Core needle (Bard Biopsy System, AZ, USA). These samples were fixed with 10% formalin and used for HE, IHC, and IF staining. Each specimen was evaluated by an experienced pathologist who was blind to clinical findings. All patients gave written informed consent to participate in this study following the ethical guidelines of the 1975 Declaration of Helsinki, and according to the process approved by the ethical committee of Osaka Metropolitan University, Graduate School of Medicine, Osaka, Japan with number 1646, 3641, and 2024-046.

### Mice

Three groups of male C57BL/6 mice (SLC, Shizuoka, Japan), ten mice per group, were given an intraperitoneal (i.p.) injection of an escalating dose of thioacetamide (TAA) (Sigma, St Louis, MO, USA) twice a week for totally 10 weeks. For the first week, mice received an i.p. injection of 50 mg/kg body weight TAA for the first dose and then 100 mg/kg TAA for the second dose; for weeks 2–3, weeks 4–5, and weeks 6–10 they received 200, 300, and 400 mg/kg twice a week respectively (cirrhosis group). The control group (Vehicle group) was injected with normal saline instead of TAA. Subsequently, the regression group was discontinued with TAA injection for 2 weeks. After anesthetized by i.p. injection of pentobarbital (Somnopentyl, Kyoritsu, Tokyo, Japan) 70 mg/kg BW, mice were euthanized two days after the last injection. Blood and liver tissue were collected.

The 8-week-old male C57BL/6 mice received i.p. injections twice weekly of TAA or a similar volume of saline (Vehicle group) for 6 or 10 weeks. During the last 3 weeks of 6 weeks TAA injections, mice were administered with VX15/2503 (#HYP99407, MCE, USA) at 0.5 or 1 𝜇𝜇g/g diluted in saline twice a week or vehicle via tail-vein injection. During the last 5 weeks of 10 weeks TAA injections, mice were administered with VX15/2503 (#HYP99407, MCE, USA) at 5 𝜇𝜇g/g BW diluted in saline twice a week or vehicle via tail-vein injection. Mice were sacrificed 2 days after the last dose of TAA and the therapeutic injection.

The 8 week-old male C57BL/6 mice received i.p. injections twice weekly of TAA or a similar volume of saline (Control group) for 8 weeks. During the last 4 weeks of TAA injections, mice were administered with Vehicle (45% saline, 5% Tween-80, 40% PEG300, and 10% DMSO) or AZD8055 (#HY-10422, MCE, USA) at 1 𝜇𝜇g/g twice a week via i.p. injection. Control group was treated with AZD8055 at 1 𝜇𝜇g/g twice a week via i.p. injection during the last 4 weeks. Mice were sacrificed two days after the last dose of TAA and the therapeutic injection.

CCl4 6 weeks, BDL 2 weeks, BDL 24 hours, DDC 6 weeks, CDAA 32 weeks mouse model were performed in our laboratory in previous studies^62, 63^.

All mice received human care, according to Guide for the Care and Use of Laboratory Animals, National Institutes of Health. All protocols and experimental procedures were approved by the Institutional Animal Care and Use Committee of Osaka Metropolitan University and were performed following the guidelines of the National Institutes of Health for the use of animals in research. Mice were housed in a temperature-controlled (24 ± 1°C) environment, with humidity levels of 55 ± 5% and alternating 12-hour light/12-hour dark cycles. Mice had free access to water and a standard rodent diet.

### Single cell fixed RNA sequencing

#### Fixation and cell dissociation from frozen liver tissues

Mouse frozen liver tissues from control, TAA-induced cirrhosis, and regression groups (n=3 per group) were used for single-cell fixed RNA sequencing. Tissue dissociation and fixation were performed using the Chromium Next GEM RNA Profiling Sample Fixation Kit (10x Genomics, PN-1000414, CA, USA) according to the manufacturer’s protocol.

Briefly, tissues were weighed to calculate Fixation Buffer volume (1 mL per 25 mg tissue), placed on a pre-chilled glass dish on dry ice, finely minced, and incubated in Fixation Buffer at 4°C for 16–24 hours. Following centrifugation at 850 x g for 5 min at room temperature (RT), supernatants were discarded. The pellet was washed twice with chilled PBS and resuspended in Resuspension Buffer on ice. Dissociation Solution (10x Genomics) was pre-warmed at 37°C for 10 min, and 2 mL was added to each sample, which was then transferred to Miltenyi C tubes (Miltenyi Biotec, Cat#130-093-237) and processed using GentleMACS Octo Dissociator with Heater (Cat#130-096-427) as recommended. Resulting cell suspensions were filtered using MACS SmartStrainers 30 μm (Cat#130-098-458) and viable cells were counted using trypan blue or propidium iodide staining (Sigma, Cat#P4170). Single-cell suspensions were stored at –80°C in Quenching Buffer containing Enhancer and Glycerol.

#### Library Preparation and Sequencing

Gene expression profiling was conducted using the Chromium Fixed RNA Profiling Reagent Kit for Singleplexed Samples (10x Genomics, CG000477 Rev C), employing the Mouse Transcriptome Probe Set. Approximately 2,000,000–3,000,000 fixed cells were hybridized for 18 hours at 42 °C with whole-transcriptome probe pairs. After post-hybridization washes, cells were encapsulated in gel beads-in-emulsion (GEMs) using the Chromium X system and Chip Q. Within each GEM, ligated probe pairs were barcoded with 10x GEM Barcodes and Unique Molecular Identifiers (UMIs), followed by heat denaturation and recovery. The barcoded products underwent pre-amplification and purification using SPRIselect beads. Final libraries were generated via sample index PCR using the Dual Index Kit TS Set A and were further purified by size selection. Library quality was assessed using an Agilent Bioanalyzer, and concentrations were determined using the KAPA Library Quantification Kit (Roche). Libraries were sequenced on the DNBSEQ-G400RS platform (MGI Tech) in paired-end mode, using 28 bp for Read 1 (containing the 16 bp GEM barcode and 12 bp UMI) and 100 bp for Read 2 (capturing the probe insert). Demultiplexing and gene expression quantification were performed using Cell Ranger v7.1.0 (10x Genomics), aligning reads to the mm10 mouse reference genome and generating filtered feature-barcode matrices for downstream analyses.

#### Dataset Integration

Nine samples (control, cirrhosis, regression) were analyzed using Seurat^64^ (v4.3.0.1). Reciprocal PCA (rpca) was applied for integration following Filliol et al^65^. After identifying 3000 variable features (‘SelectIntegrationFeatures’), anchors were identified (‘FindIntegrationAnchors’, rpca method), and datasets were integrated (‘IntegrateData’). Downstream steps included scaling (‘ScaleData’), dimensionality reduction (‘RunPCA’, ‘ElbowPlot’), clustering (‘FindNeighbours’, ‘FindClusters’), and visualization (‘RunUMAP’). Manual sub-clustering was performed using ‘CellSelector’ when necessary, followed by re-clustering.

#### Cell type annotation

Cell-type identities were assigned based on differentially expressed genes (DEGs) within each cluster (‘FindMarkers’) using canonical markers and validation in PanglaoDB^66^ and CellMarker^67^ databases. For hepatocytes zonation analysis, pericentral and periportal signature score were calculated using AddModuleScore based on canonical markers of pericentral (Cyp2e1, Cyp1a2, and Glul) or periportal (Asl, Cyp2f2) hepatocytes.

#### Trajectory Analysis

Cell-state transitions were analyzed using Monocle2^68^ (v2.26.0) or Monocle3^69^ (v1.3.1). Seurat objects were converted to Monocle objects (‘as.cell_data_set’). Clustering (‘cluster_cells’), graph learning (‘learn_graph’), and pseudotime ordering (‘order_cell’) were conducted. Genes varying along trajectories were identified using ‘graph_test’.

#### Gene Set Enrichment Analysis (GSEA)

GO enrichment analysis was conducted using enrichR^70^ (v3.2) and ClusterProfiler^71^ (v4.6.2) against “GO_Biological_Process_2023,” “GO_Cellular_Component_2023,” and “GO_Molecular_Function_2023”. DEGs with average log2FC > 0.5 (max 100 genes) per cluster were used for input.

#### Transcription Factor (TF) Analysis

TF regulatory networks (regulons) were identified using SCENIC^72^ (v1.3.1) or Dorothea^73^ (v1.18.0). Regulon activity scores (AUC) were calculated and binarized using the BIC threshold. For clusters with > 50% active cells, median AUC was applied for thresholding.

#### Cell-Cell Interaction Analysis

CellChat: Cell-cell communication networks were inferred using CellChat^74^ (v1.6.1) with CellChatDB. Communication probability was estimated based on ligand-receptor expression levels, and interaction significance was assessed via permutation tests.

MultiNichNet^75–77:^ MultiNichNet was used to infer condition-specific ligand-receptor interactions and their downstream target genes. DE analysis and pseudo-bulk expression were used to calculate interaction specificity metrics, which were scaled and aggregated to rank ligand-receptor pairs across conditions.

#### External Datasets

Publicly available single-cell datasets from mouse and human liver tissues were obtained from GEO: Filliol et al.^65^ (GSM6508438, GSM6508440, GSM5257943), Wang et al.^36^ (GSE212837), Gribben et al.^78^ (GSE202379)

Spatial transcriptomics data from healthy human livers and PSC patients were obtained from MacParland lab^79^ (https://macparlandlab.shinyapps.io/healthylivermapspatialgui/ https://macparlandlab.shinyapps.io/shinypsc_spatialappgui/)

### Primary mouse liver cell isolation

Primary mouse liver non-parenchymal cells (NPCs) were isolated from the control group, mice subjected to TAA injection for 10 weeks, and mice with regression after 2 weeks as previously described protocol with some modifications^80^. Briefly, liver cell suspensions were obtained by in vivo pronase-collagenase perfusion, and then livers were surgically removed and cut into small pieces. Tissues were transferred in SC-2 containing 0.2% collagenase IV (0.5 mg mL^-1^, Cat#032-22364, Lot#WTL5857, Wako, Japan), pronase E (0.4 mg mL^-1^, Cat#9036-06-0, Merck, USA) and DNase I (0.1 mg mL^-1^, Cat#11284932001, Roche, Switzerland), and then incubated for in vitro digestion at 37 °C in a water bath for 30 minutes. Cell suspensions were centrifuged at 100 x g for 5 minutes to remove hepatocytes. The supernatants were pelleted at 300 x g for 10 minutes. Then cell suspensions were filtered using a 30 μm nylon cell strainer to remove clumps and cell debris. These non-parenchymal

Primary mouse hepatic stellate cells (mHSCs) were isolated from three groups in our laboratory using a pronase-collagenase digestion method as previously described with some modifications^81^. Primary mouse hepatocytes (mHEPs), liver sinusoid endothelial cells (mLSECs), Kupffer cells (mKCs), Monocytes-derived macrophages (mMomacs), and Cholangiocytes (mCHOLs) were isolated from three groups in combination with low-speed centrifugation and magnetic-activated cell sorting (MACS). Mouse livers were in situ perfused with 50 ml SC-1 solution and 50 ml 0.05% Collagenase IV solution. Then mHEPs were pelleted by centrifugation at 100 x g for 5 minutes, two times. The dead hepatocytes were removed by using Percol 45% solution in DMEM and were centrifuged at 1000 rpm for 10 minutes, 4℃. The supernatants were centrifuged at 300 x g for 10 minutes to collect NPCs. Primary mLSECs, mKCs, mMomacs, and mCHOLs were isolated from NPCs by using a 2-step protocol incubated with CD146 MicroBeads (#130-092-007, Miltenyi Biotec, Germany), or F4/80 MicroBeads UltraPure (#130-110-443, Miltenyi Biotec, Germany), or Cd11b MicroBeads (#130-097-142, Miltenyi Biotec, Germany), or Epcam MicroBeads (#130-105-958, Miltenyi Biotec, Germany), respectively; using the LS column (#130-042-40, Miltenyi Biotec, Germany). Then, primary mHSCs, mHEPs, mLSECs, mKCs, mMomacs and mCHOLs were cultured in the appropriate medium for 72 hours, cell lysates were harvested for quantitative real-time PCR analysis, immunoblotting analysis, and immunofluorescence staining. Conditioned media were harvested for ELISA and/or cytokine array experiment.

Primary liver mouse lymphocytes were isolated with previous publication with slight modification^12^. Livers were perfused through portal vein with PBS, and then minced finely. The minced tissues were mixed with liberase and DNase I and vibrated 30 minutes in water bath (37℃). The cell suspensions were filtered through cell strainer 70 𝜇𝜇m, and centrifuged at 450 x g, 5 minutes, 20 ℃. Cells were suspensed into 35% Percoll and centrifuged at 2000 rpm, 20 minutes, 4 ℃. The pellet was washed with 2% PBS-FBS and filtered through 30 𝜇𝜇m cell strainer. These cells were used for flow cytometry experiments.

### Treatment assay

HHSteCs were seeded into 12, 24, or 96-well plates (Greiner Bio-One, Tokyo, Japan), at 1 × 10^5^, 0.5 × 10^5^, 5.000 cells/well in a final volume of 1 mL, 0.5 mL, and 0.1 mL of the appropriate medium, respectively, for the performance of protein and mRNA analysis or other assays described below.

For the experiments in spontaneous activation, HHSteCs were seeded into Stellate Cell Medium (SteCM, #5301, ScienCell, CA, USA), supplemented with 2% FBS, 1% S (SteCGS, #5352), 1% P/S (S+) or with 2% FBS%, 1% P/S (S-). On the following day, recombinant human IGFBP6 (#876-B6, R&D Systems, Minneapolis, MN, USA), OPG (#6945-OS, R&D Systems, Minneapolis, MN, USA) were added, in a dose (0-1000 ng/ml)-dependent manner. In the experiment with TGFβ1–induced activation, HHSteCs were seeded into 24-well plates at 0.5 × 10^5^ cells/well in complete SteCM. On the next day, the medium was replaced with fresh 2% FBS/SteCM without the supplement. Recombinant human TGFβ1 2 ng/ml (rhTGFβ1, #7754-BH, R&D Systems, Minneapolis, MN, USA) were added simultaneously into medium and incubated for 24 hours.

LX2 were seeded into Dulbecco’s Modified Eagle Medium (DMEM, #11885084, Gibco, Grand Island, NY, USA), supplemented with 2% FBS, 1% P/S. On the following day, soluble recombinant human SEMA4D (#310-29, Peprotech, NY, USA) were added, in a dose (0-1000 ng/ml)-dependent manner for 48 hours. RNA and protein were isolated for molecular analysis.

HHSteCs were seeded into SteCM (#5301, ScienCell, USA) supplemented with 2% FBS, 1% S (S+), 1% P/S. On the following day, HHSteCs were changed to medium without supplement (S-). At day 3, HHSteCs were treated with AZD8055 (HY-10422) or All trans retinoic acid (HY14649) or Bexarotene (HY14171) or Adapalene (HY-B0091) or Tarazotene (HY-15388) (all from MedChem, Japan) dose-dependently diluted in medium. rhTGFβ1 were added after 4 hours simultaneously into medium and incubated for 24 hours. RNA and protein were isolated for molecular analysis.

### Quantitative real-time PCR (qRT-PCR) assay

Total RNA was extracted from cells or liver tissues using NucleoSpin RNA (MACHEREY-NAGEL, #Cat740955.250) according to the manufacturer’s instructions. cDNA was synthesized using ReverTra Ace kit (#FSQ-101, Toyobo, Osaka, Japan). Gene expression was measured by qRT-PCR using cDNA, THUNDERBIRD SYBR qPCR Mix Reagents (Toyobo, Osaka, Japan), and a set of gene-specific oligonucleotide primers and probes (Supplementary Data 8) using an Applied Biosystems Prism Fast 7500 system (Applied Biosystems, Tokyo, Japan). Glyceraldehyde-3-phosphate dehydrogenase (Gapdh) or 18S level was used to normalize relative mRNA abundance.

### Western blot

Proteins isolated from tissues (20 µg) or cells (4 – 6 µg) were subjected to SDS-PAGE and transferred to Immuno-Blot® PVDF membranes (Bio-Rad, CA, USA). After blocking with 5% skim milk, the membranes were incubated with the primary antibodies overnight at 4°C. Membranes were then labeled with horseradish peroxidase-conjugated secondary antibodies (1:2000, Dako, Agilent Technologies, Santa Clara, CA, USA). Immunoreactive bands were visualized by enhanced chemiluminescence using L-012 substrate (ImmunoStar LD, Wako, Osaka, Japan) and documented with a Fusion Solo S machine (VILBER, Marne-la-Vallée Cedex 3, France). GAPDH was used as the loading control. The quantitative density band was analyzed by ImageJ software. The primary antibodies used are listed in Supplementary Data 9.

### Hematoxylin-Eosin (H&E), Immunohistochemistry (IHC), and Immunofluorescence (IF) staining

H&E staining, IHC, and IF analysis were performed as described previously^62, 63^. The primary antibodies used are listed in Supplementary Data 9. SEMA4D IHC was stained using Takara Pod Conjungate Anti Rabbit (MK205, Takara, Japan). Double IHC was performed using ImmPRESS® Duet Double Staining Polymer Kit (MP-7724-15, VectorLabs, CA, USA).

For quantification of liver cirrhosis, paraffin liver sections (5 µm) were stained with Picro-Sirius Red Stain Kit (#PSR1, Logan, Utal, USA). Images were captured using a BZ-X700-All-in-One fluorescence microscope (Keyence Co., Osaka, Japan). Positively immune-stained cells were counted in number by taking 10 fields at magnification ×100, ×200 or ×400 without overlapping per section. Percentages of positive areas per corresponding whole lobe area were calculated using BZ-X Analyzer software.

### Measurement of AST and ALT

Aspartate transaminase (AST), and alanine transaminase (ALT) were measured in serum using a commercially available kit (Wako, Osaka, Japan) according to the manufacturer’s protocol.

### Proteome cytokine array

Secreted proteins in culture medium from isolated LSECs, macrophages, cholangiocytes and mouse serum were screened by using Proteome Mouse XL Cytokine Array Kit (#ARY028, R&D system, Minneapolis, MN, USA), which can detect 111 soluble proteins. Phosphoprotein in cell lysate from HHSteCs treated with DMSO or AZD8055 1 𝜇𝜇M in present of TGF𝛽𝛽1; GFP and LMCD1 overexpression HHSteCs lysates were assessed using human phosphokinase array (#ARY003C, R&D system, Minneapolis, MN, USA), which can detect 37 phosphokinase protein and 2 related total protein. Signal was obtained using a Fusion Solo S Machine (VILBER, Marne-la-Vallée Cedex 3, France). Spot analysis was performed by ImageJ.

### Multiplex flow cytometry and fluorescence-activated cell sorting (FACS)

Using multiple color staining, mouse non parenchymal cells or CD11b+macrophages or F4/80+macrophages were pre-incubated with unlabel CD16/32 mAb (2.4G2, Bioxel, Lebanon, USA) to avoid non-specific binding of antibody to Fc𝛾𝛾R. Dead cells were excluded using Zombie NIR fixable viability dye. Then cells were incubated with CD45, CD11B, F4/80, FOLR2, MHCII, Ly6G, NK1.1, CD4, CD8, KLGR1, FOXP3 antibody. Detailed antibody information is shown in Supplementary Data 9. Stained cells were analyzed using Attune NxT (Thermo Fisher, Tokyo, Japan) and data was analyzed by FlowJo software (Version 10.8.1).

Mouse macrophages cell sorting for culturing was performed on SH 800 Cell Sorter (SONY). Viable KC1 (CD45+F4/80+MHCII-Cd11b-Folr2+) and KC2 (CD45+F4/80+MHCII+CD11b-Folr2+) were isolated from F4/80+ macrophages of healthy control, cirrhosis and regression group (n=3, each group). Viable Momacs (Cd45+Cd11b+F4/80-Ly6G-) and neutrophils (Cd45+Cd11b+F4/80-Ly6G+) were isolated from CD11b+ macrophages of healthy control, cirrhosis, and regression group (n=3, each group). KC1, KC2, and Momacs were plated in 24-well plated in RMPI-1640 containing 10% fetal bovine serum and 1% penicillin/streptomycin for 72 hours. Conditional media were collected and centrifuged at 500 g for 10 minutes and stored at - 80°C.

### ELISA

Enzyme-linked immunosorbent assay kits specific for mouse SEMA4D (#OO9126, RayBiotech, USA), mouse MMP7 (#Q1O738, RayBiotech, USA), and human SEMA4D (#ELH-SEMA4D, RayBiotech, USA) were used according to the manufacturer’s protocol. Briefly, cell culture supernatants or human/mouse serum were added to the wells, followed by the biotinylated antibody. After incubation, streptavidin was added to the well. Next, wells were washed to remove unbound materials. Then, TMB One-Step Substrate Reagent was added to catalyze reactions. Finally, after adding stop solution, generated signals were measured at 450nm. The results were then calculated using the standard curve method.

### LMCD1 silence and overexpression in HHSteCs

HHSteCs were cultured into 24-well plates for 24 hours until they reached 70% confluence. Cells were then transfected with siRNAs using Lipofectamine RNAi MAX (#13778-150, Thermo Fisher Scientific). Briefly, Lipofectamine RNAiMAX was mixed with 5 pM negative control siRNA (siNC, #4390843, Thermo Fisher Scientific, Japan) or 5 pM LMCD1 siRNA (#134439 and #134440; Thermo Fisher Scientific, Japan) in a serum-free medium (Opti-MEM, #31985070, Thermo Fisher Scientific, Japan), for 15 minutes at room temperature to form complexes. The transfection mixture was added to each well. The cells were incubated at 37°C in a humidified CO2 incubator for 48 hours, followed by the addition of TGF𝛽𝛽 1 (2 ng/ml) into the culture medium for 24 hours before gene expression and immunoblotting analysis.

HHSteCs were cultured into 6-well plates for 24 hours until they reached 70% confluence in SteCM with Supplement (S+). Then, 10 𝜇𝜇L of LMCD1 adenovirus (#26901051, Abm, Canada) or GFP adenovirus (#000541A, Abm, Canada) were added to HHSteCs medium. LMCD1 and GFP transduction were evaluated at 48, 72, 96 hours after transfection at RNA and protein level.

### Statistical analysis

Statistical significance was determined using GraphPad Prism v.10.0. Data are expressed as the mean ± standard deviation. Statistical analyses were performed using two-tailed Student’s t-tests to compare differences between two unpaired groups. One-way analysis of variance (ANOVA), followed by Dunnett’s *post hoc* test, was performed to compare multiple groups with control. Two-way ANOVA, followed by Tukey’s HSD test, was performed to compare mean differences between groups. *P* < 0.05 was considered significant.

### Statistical analysis

Statistical significance was determined using GraphPad Prism v.10.0. Data are expressed as the mean ± standard deviation. Statistical analyses were performed using two-tailed Student’s t-tests to compare differences between two unpaired groups. One-way analysis of variance (ANOVA), followed by Dunnett’s *post hoc* test, was performed to compare multiple groups with control. Two-way ANOVA, followed by Tukey’s HSD test, was performed to compare mean differences between groups. *P* < 0.05 was considered significant.

## Data availablity

All data supporting the findings of this study are available within the article and associated supplementary information files. ScRNA-seq data supporting these finding have been deposited in the GEO database and can be accessed at [https://www.ncbi.nlm.nih.gov/geo/persistent-specific URL pending acceptance]. The liver atlas of integrated fixed single cell RNA sequencing data is available for exploration with our shiny app.

For all cell types mapping:

1. https://shinyapps.io/ persistent-specific URL pending acceptance

For subgroups mapping (Hepatocytes, Macrophages, Cholangiocytes, HSCs, LSECs, NK&T cells, Neutrophils, and B cells)

1. https://shinyapps.io/ persistent-specific URL pending acceptance
2. https://shinyapps.io/ persistent-specific URL pending acceptance
3. https://shinyapps.io/ persistent-specific URL pending acceptance
4. https://shinyapps.io/ persistent-specific URL pending acceptance
5. https://shinyapps.io/ persistent-specific URL pending acceptance
6. https://shinyapps.io/ persistent-specific URL pending acceptance
7. https://shinyapps.io/ persistent-specific URL pending acceptance
8. https://shinyapps.io/ persistent-specific URL pending acceptance
9. https://shinyapps.io/ persistent-specific URL pending acceptance

## Code availablity

Code used for the analysis of single cell fixed RNA profiling data are available in a public repository at https://github.com/ persistent-specific URL pending acceptance

## Acknowledgments

We thank Prof. Massimo Pinzani, and Prof. Nina Tirnitz-Parker for their helpful discussion. P.M.D was awarded the Japanese Government (MEXT) Scholarship for the Ph.D. couse. L.T.T.T. received a Grant-in-Aid for Scientific Research from the Japan Society for the Promotion of Science (JSPS Kaken C, 22K08083). N.K. received a Grant-in-Aid for Scientific Research from JSPS (J192640002), and a grant for a research program on hepatitis from the Japan Agency for Medical Research and Development (AMED-J202620103)

## Author Contributions

P.M.D and L.T.T.T. provided the concept and design, acquired data, analyzed and interpreted the data, performed all of the experiments, drafted the manuscript, and obtained funding. H.H., N.T.H., P.T.A, H.Y., T.M., V.T.H., Y.C., R.Y., D.O. performed animal studies and experiments. H.I., H.F.acquired human samples and data analysis. N.O., F. T., performed critical revisions of the manuscript for important intellectual content and obtained funding; T. K., D. B., Y. I., J.G-S. performed critical revisions of the manuscript for important intellectual content. N.K. and L.T.T.T. contributed to the study concept and design, drafted the manuscript, performed critical revisions of the manuscript for important intellectual content, obtained funding, and supervised the study. All authors had final approval of the submitted version.

## Additional information

Correspondence and requests for materials should be addressed to N. K and L.T.T.T.

---

**Supplementary Fig. 1.**
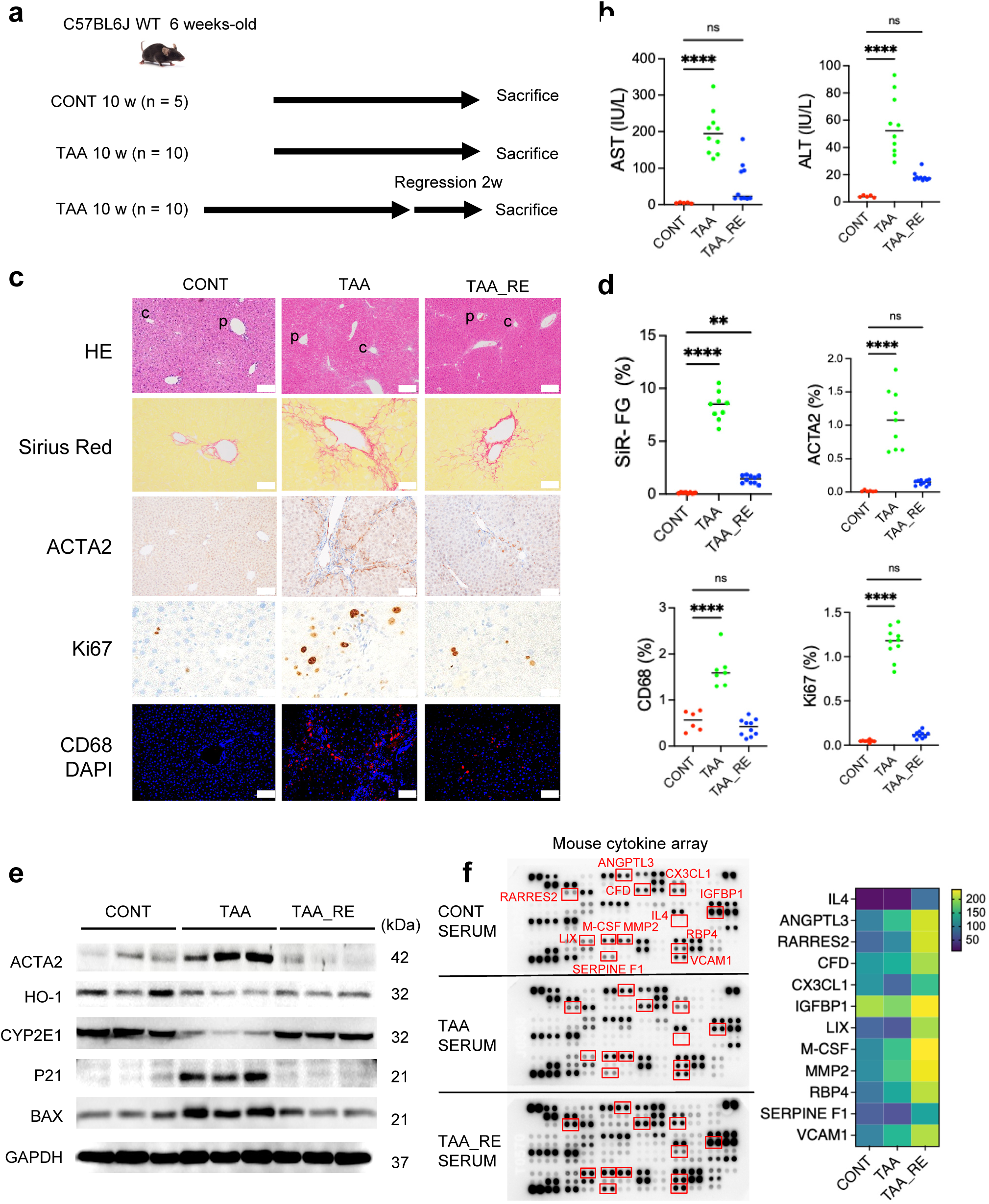
Animal model of advanced liver fibrosis progression and regression. **a.** Schematic diagram of TAA induced liver fibrosis with or without recovery state. **b.** Serum levels of AST and ALT. **c-d.** Representative liver images of H&E, Sirius Red, IHC staining of ACTA2, Ki67, IF staining of CD68 (CD68-red, DAPI-blue) and their quantifications, scale bar 50 𝜇m. **e.** Immunoblotting analysis for ACTA2, HO-1, CYP2E1, P21, BAX. GAPDH was used as internal control. **f.** Cytokine array of soluble factors in serum of control, cirrhosis, and in the regression group, the soluble factors with predominant change were marked with red boxes (left panel) and heatmap showing the quantification of mean pixel density (MPI), colored by MPI (right panel). For **b** and **d**, data represent the means ± SD. *p*-values were analyzed by one-way ANOVA followed by Tukey’s multiple comparison test. \**p* < 0.05, \*\**p* < 0.01, \*\*\**p* < 0.001, \*\*\*\**p* < 0.0001.

**Supplementary Fig. 2.**
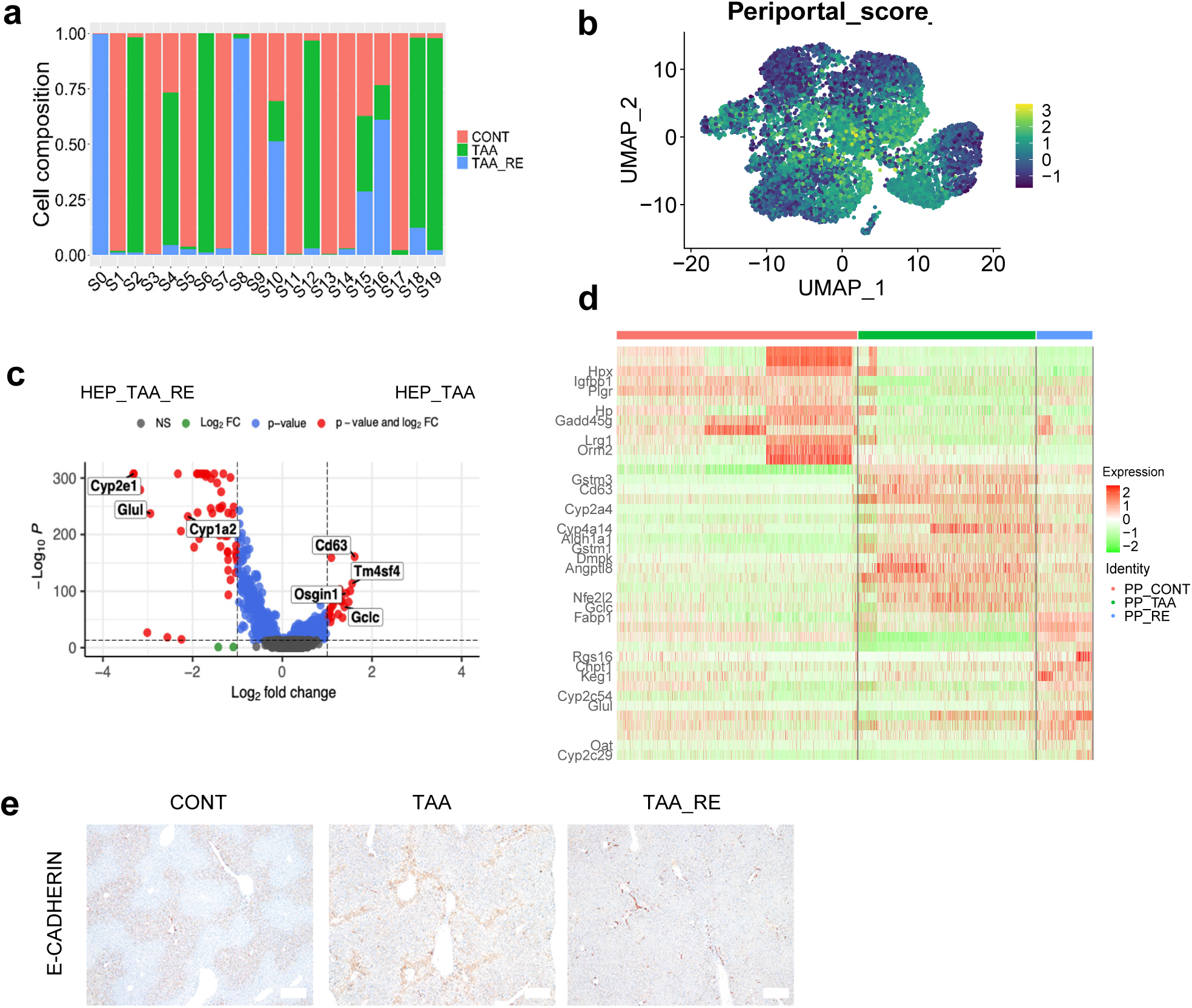
Periportal hepatocytes in cirrhosis progression and regression. **a.** Barplot showing group composition breakdown per cluster (fraction of total cell count per cluster), colored by group. **b.** UMAP showing periportal signature score of hepatocytes subpopulation in control, fibrosis, and regression group. **c.** Volcano plot showing hepatocytes differential gene expression between fibrosis and regression group. **d.** Heatmap showing representative genes expressed by periportal hepatocytes in the control, cirrhosis, and regression groups. **e.** IHC staining showing expression of E-cadherin, scale bar 50 𝜇m.

**Supplementary Fig. 3.**
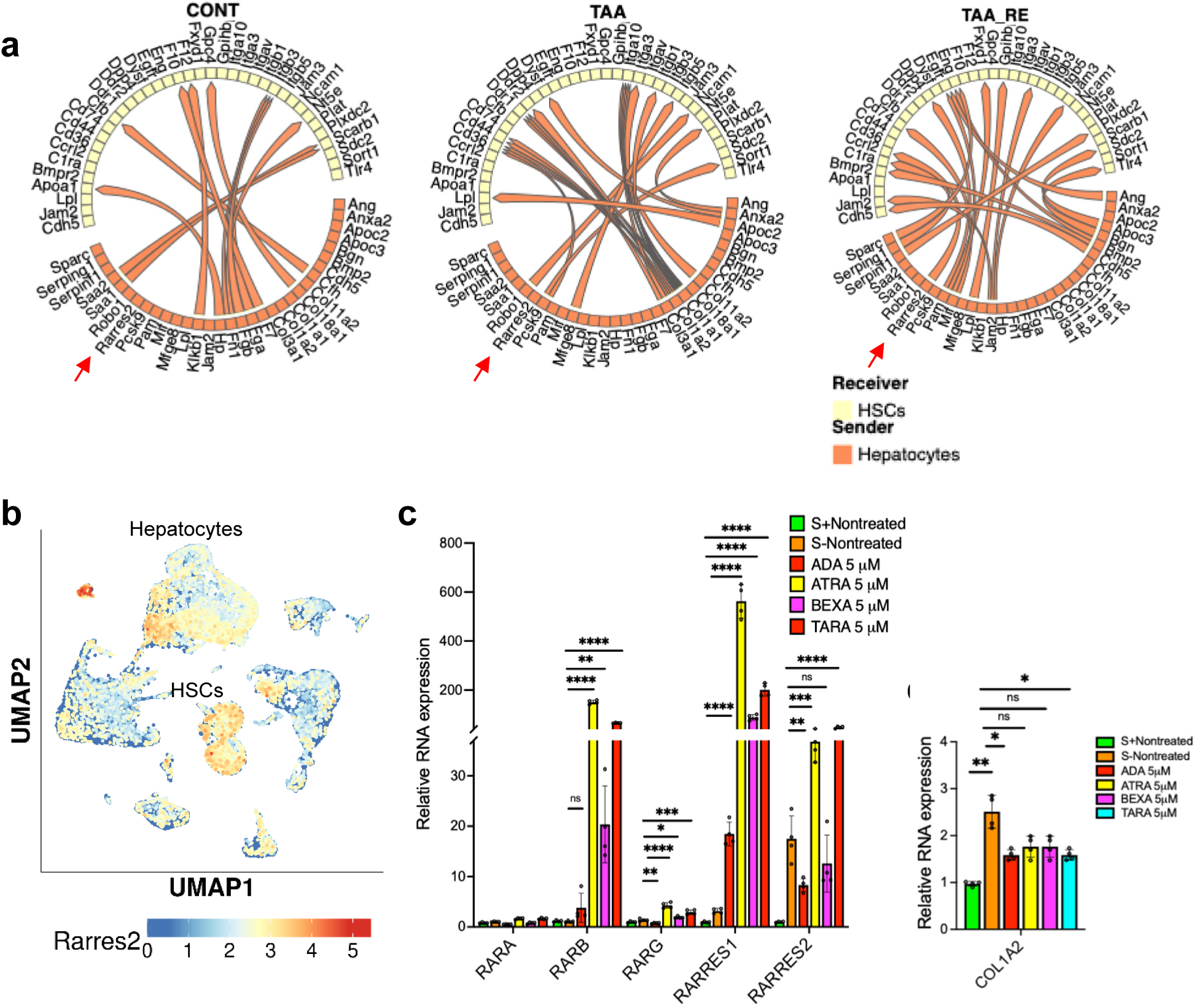
HEPs and HSCs interaction analysis. **a.** Circos plots showing interaction in hepatocytes-HSCs niche. Red arrow indicated Rarres2-Cclr2 axis. **b.** UMAP showing expression of Rarres2, colored by expression. **c.** Quantitative RT-PCR analysis of RARA, RARB, RARG, RARRES1, RARRES2 in HHSteCs treated with ADA 5 𝜇M, ATRA 5 𝜇M, BEXA 5 𝜇M and TARA 5 𝜇M. **d.** Quantitative RT-PCR analysis of COL1A2 in HHSteCs treated with ADA 5 𝜇M, ATRA 5 𝜇M, BEXA 5 𝜇M and TARA 5 𝜇M. 18S was used as internal control. Data represent the means ± SD. *p*-values were analyzed by Student’s t-test. \**p* < 0.05, \*\**p* < 0.01, \*\*\**p* < 0.001, \*\*\*\**p* < 0.0001

**Supplementary Fig. 4.**
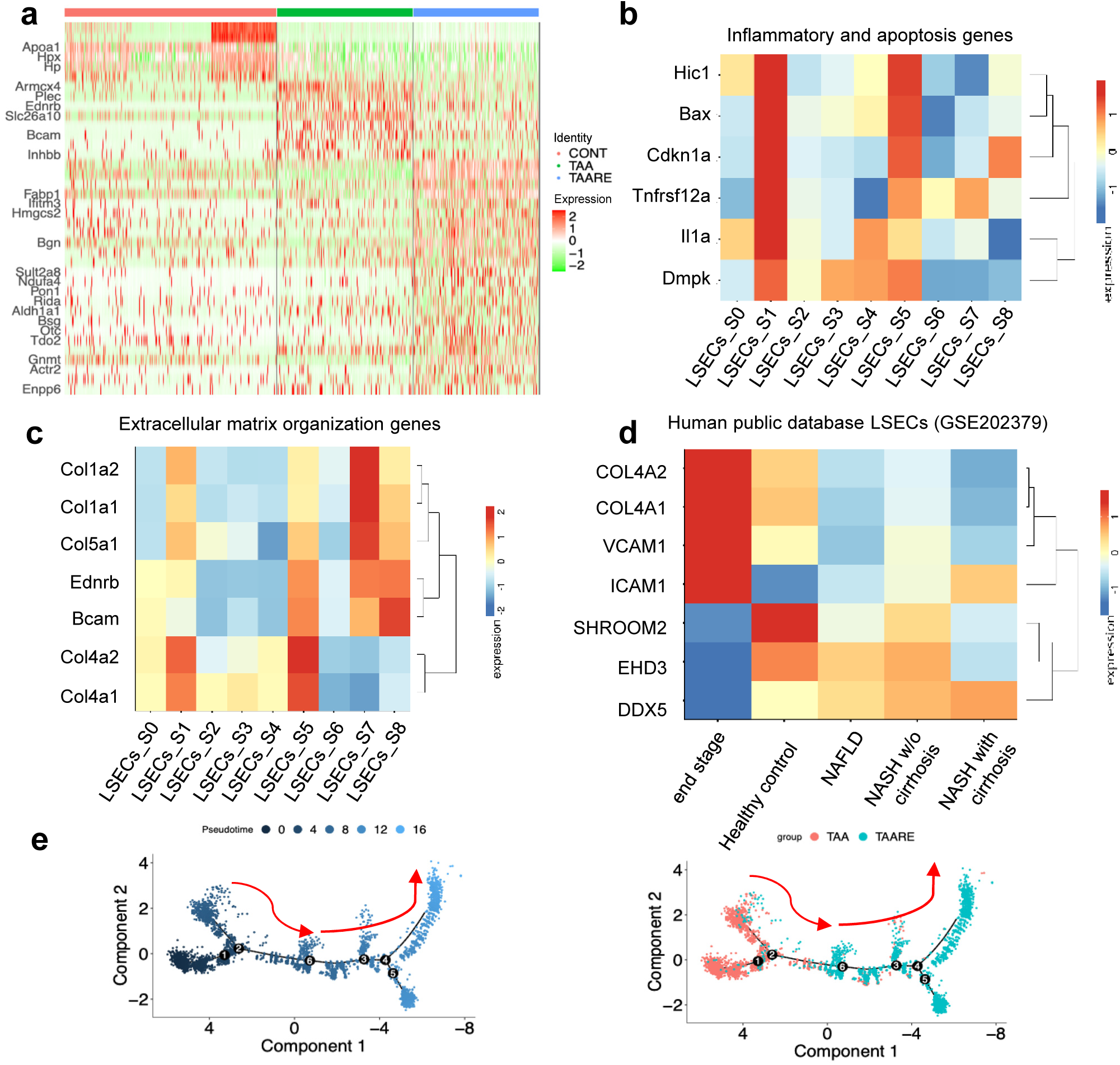
Heterogeneity of LSECs in fibrosis progression and regression. **a.** Heatmap showing top gene expression among control, fibrosis, and regression group. **b.** Heatmap showing inflammatory and apoptosis genes expression among clusters**. c.** Heatmap showing extracellular matrix organization gene expression among clusters. **d.** Heatmap showing expression of quiescent markers (SHROOM2, EHD5, and DDX5) and EMT markers (COL4A1, COL4A2, VCAM1, and ICAM1) of LSECs in human public database (GSE202739). **e.** Pseudotime trajectory mapped to LSECs UMAP coordinates. Arrows indicated predicted cell state progression from fibrosis to regression phase.

**Supplementary Fig. 5.**
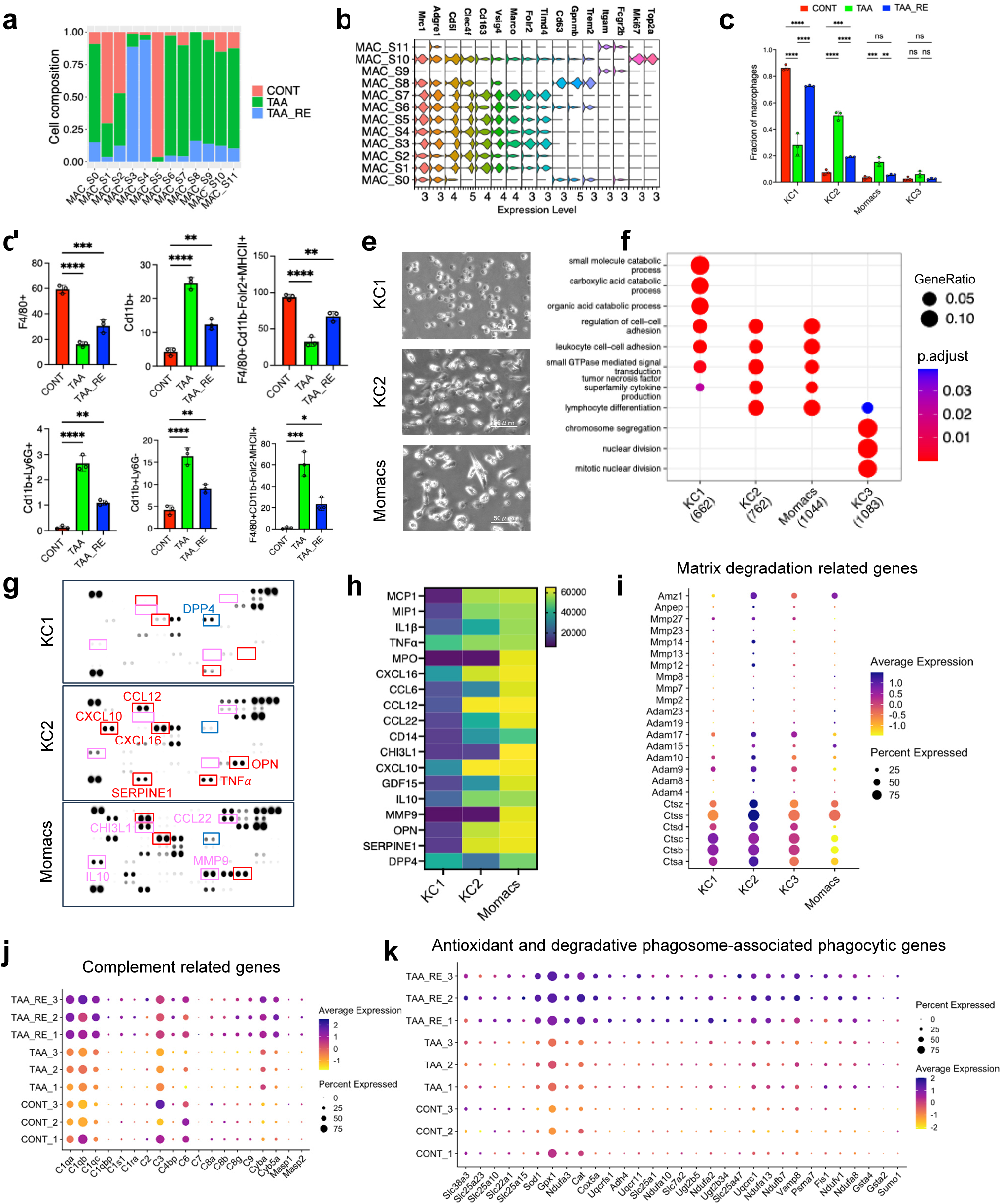
Heterogeneity of macrophages in liver fibrosis progression and regression. **a.** Barplot showing group composition breakdown per cluster (fraction of total cell count per cluster), colored by group. **b.** Stacked violin plot showing classified markers of macrophages. **c.** Barplot showing macrophages fraction from control, cirrhosis, and regression group, colored by group. **d.** Quantitative analysis of KC1, KC2, Momacs, and Neutrophils subpopulation by flow cytometry in healthy, liver fibrosis progression and regression. Data represent the means ± SD. *p*-values were analyzed by one-way ANOVA followed by Tukey’s multiple comparison test. \**p* < 0.05, \*\**p* < 0.01, \*\*\**p* < 0.001, \*\*\*\**p* < 0.0001. **e.** Contrast images showing of KC1, KC2, and Momacs morphology, scale bar 50 𝜇m. **f.** Dotplot showing the comparison of gene set enrichment analysis of KC1, KC2, Momacs, and KC3 by ClusterProfiler. **g.** Cytokine array of soluble factors in the medium of KC1, KC2, Momacs. The soluble factors with predominant changes in KC2 and Momacs was marked with red and magenta boxes, respectively. The soluble factors with predominant change in KC1 was marked with blue. **h.** Heatmap showing the cytokines quantification by mean pixel density, colored by MPI. **i.** Dotplot showing KC2 signatures. **j.** Dotplot showing complement phagocytosis related genes. **k.** Dotplot showing degradative phagosomes related genes.

**Supplementary Fig. 6.**
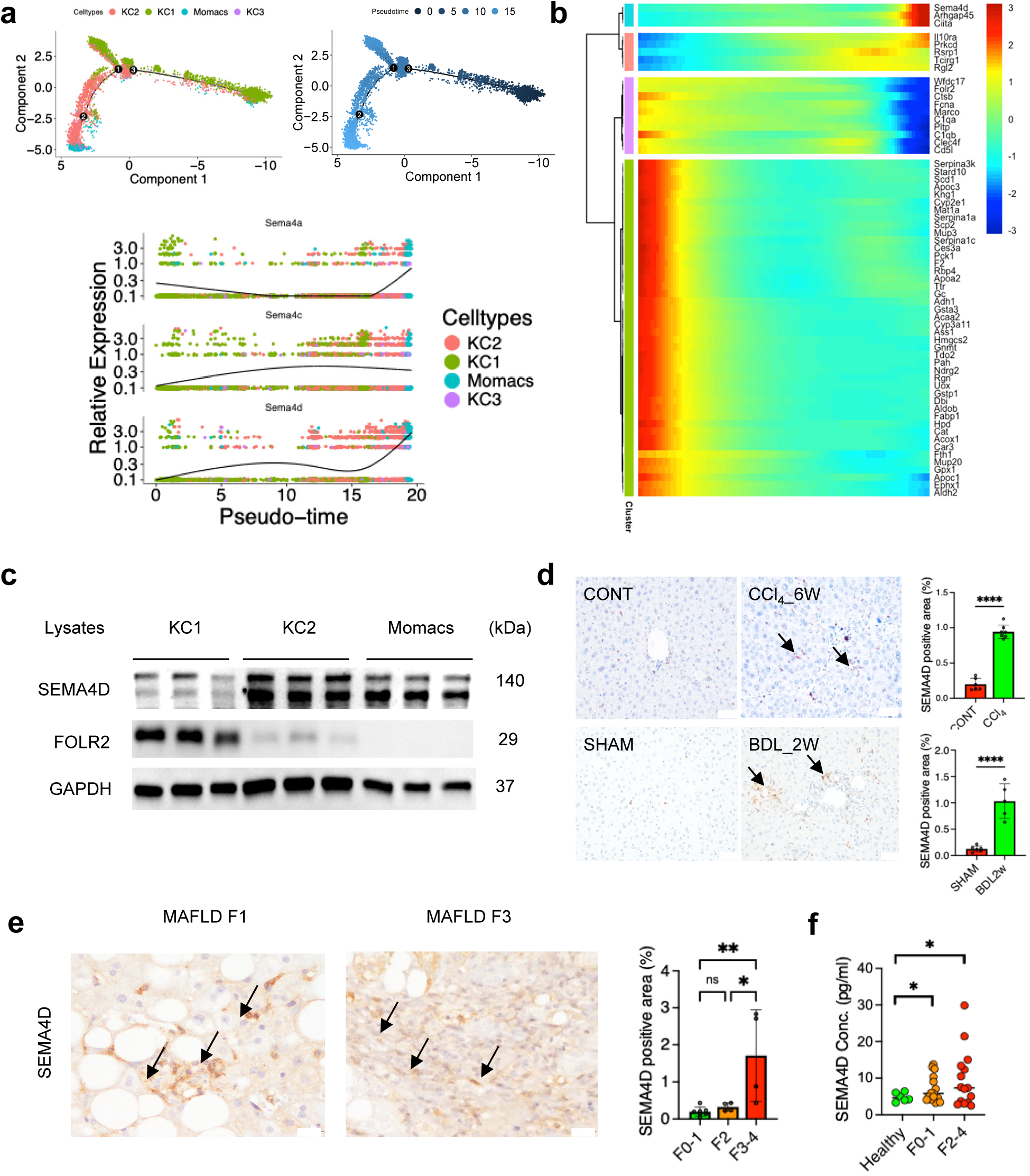
Expression of Semaphorin 4D in mouse and human fibrosis livers. **a.** Pseudotime analysis mapped to HSCs UMAP coordinates. **b.** Heatmap showing gene module expression following pseudotime and expression of Sema4a, Sema4c, and Sema4d across pseudotime **c.** Immunoblotting showing the expression of SEMA4D, FOLR2 in KC1, KC2, and Momacs. GAPDH was used as internal control. **d.** IHC staining showing expression of SEMA4D (arrow) in CCl_4_ 6 weeks and BDL 2 weeks mouse model and their quantification. Scale bar 50 𝜇m. **e.** IHC staining showing expression of SEMA4D (black arrow) in human cirrhosis (MASLD F1 and F3), scale bar 50 𝜇m **f.** Barplot showing SEMA4D expression in liver biopsy of F0-1, F2, F3-4 MASLD patients (left panel) and SEMA4D concentration in serum of healthy, F0-1, and F2-4 MASLD patients (right panel). Data represent the means ± SD. *p*-values were analyzed by one-way ANOVA followed by Tukey’s multiple comparison test. \**p* < 0.05, \*\**p* < 0.01, \*\*\**p* < 0.001, \*\*\*\**p* < 0.0001.

**Supplementary Fig. 7.**
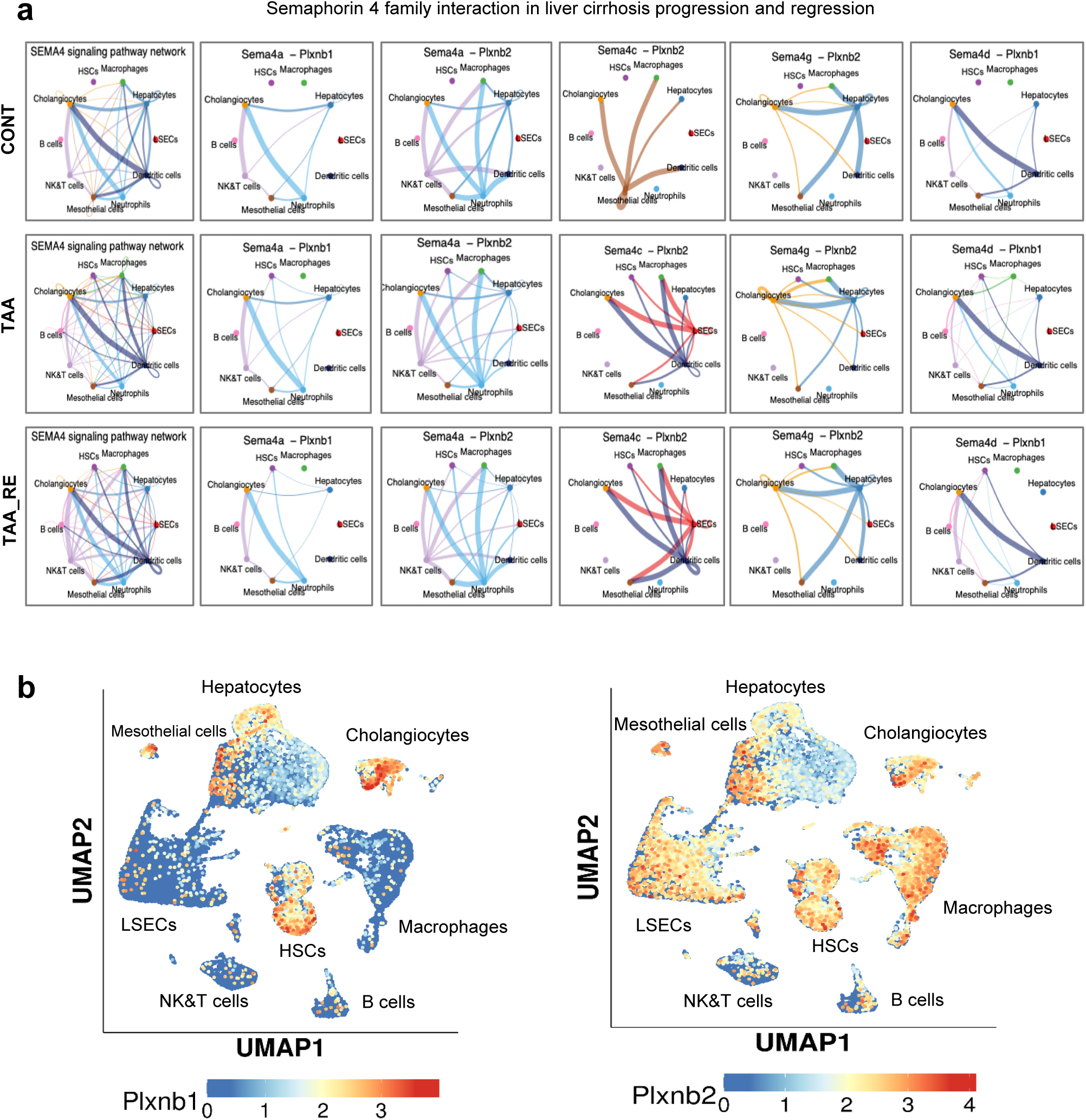
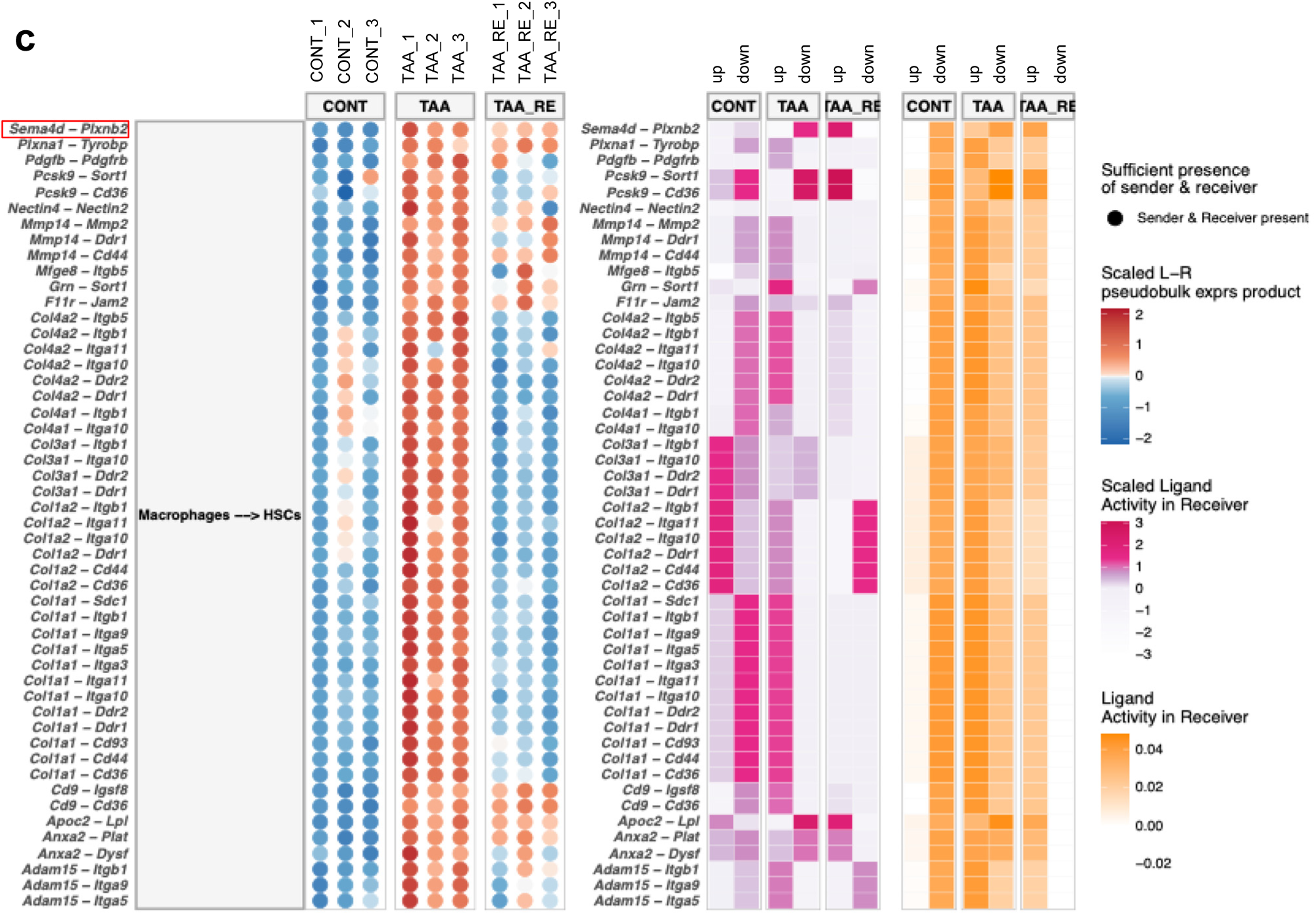
SEMA4 signaling pathways in liver fibrosis progression and regression. **a.** Circle plot showing cell-cell interaction network of Sema4 signaling pathway in healthy, liver fibrosis progression and regression. **b.** UMAP showing expression of Plxnb1 and Plxnb2 in single cell data, colored by expression level. **c.** Bubble plot showing top 50 interaction from macrophages to HSCs upregulated in fibrotic livers by Multinichenet, colored by Scaled L-R pseudobulk expression.

**Supplementary Fig. 8.**
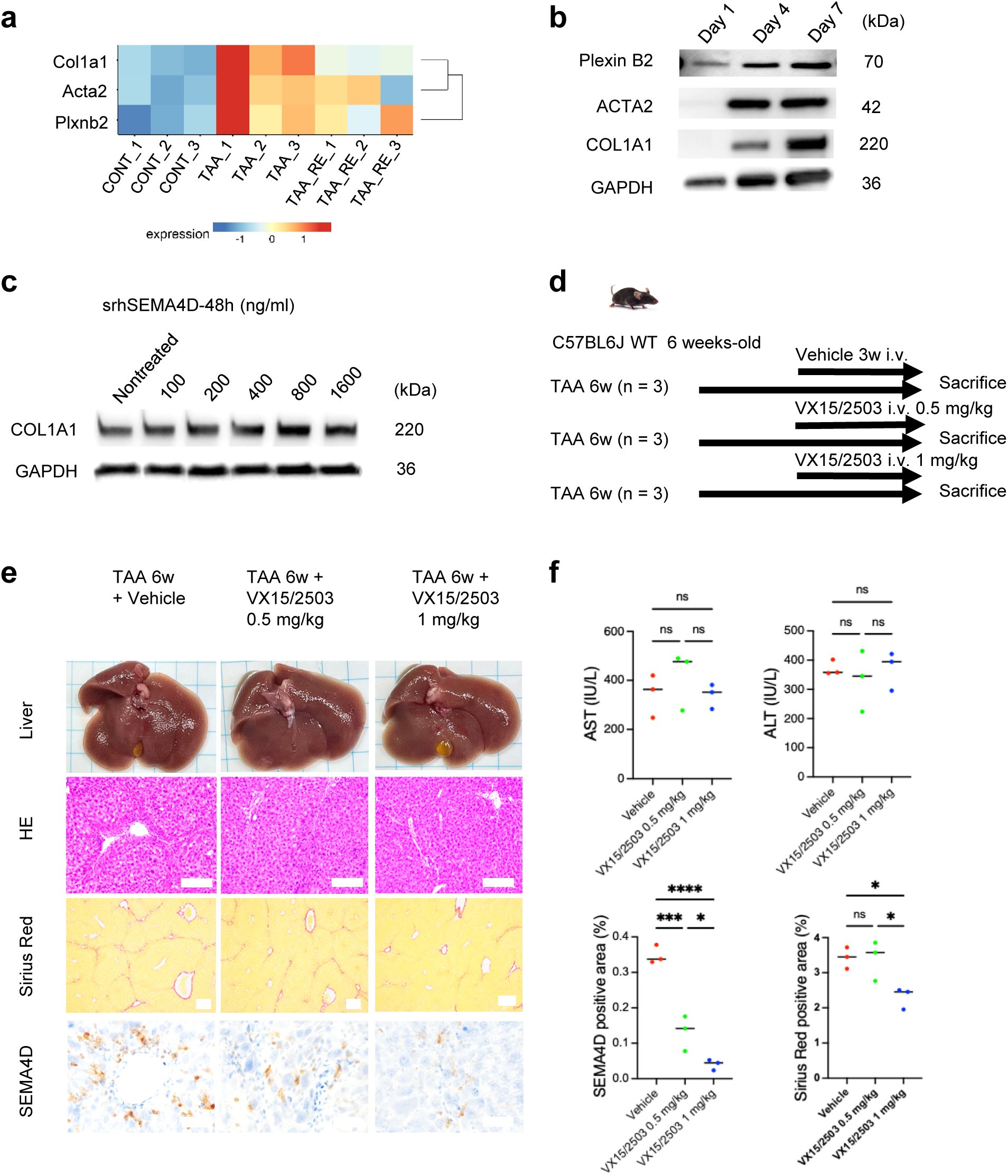
Sema4d-Plexin B2 interaction induced HSC activation and liver fibrosis development. **a.** Heatmap showing the expression of Plxnb2, Col1a1, and Acta2 in HSCs dataset from healthy, fibrosis and regressed livers. **b.** Immunoblotting showing expression of PLEXIN B2, ACTA2, and COL1A1 in mHSCs at day 1, day 4, and day 7. GAPDH was used as internal control. **c.** Immunoblotting showing expression of COL1A1 in LX2 treated with srhSEMA4D at 48 hours. GAPDH was used as internal control. **d.** Schematic overview of experimental design for **e**-**f**. **e.** Representative liver images of H&E, Sirius Red, SEMA4D IHC staining of 3 groups: TAA 6 weeks treated with vehicle, TAA 6 weeks treated with VX15/2503 at the dose of 0.5 mg/kg or 1 mg/kg (n = 3, each group), scale bar 20 𝜇 m. **f.** Measurement of AST, ALT among 3 groups (upper panel) and quantification of Sema4d and Sirius Red staining (lower panel). Data represent the means ± SD. *p*-values were analyzed by one-way ANOVA followed by Tukey’s multiple comparison test. \**p* < 0.05, \*\**p* < 0.01, \*\*\**p* < 0.001, \*\*\*\**p* < 0.0001.

**Supplementary Fig. 9.**
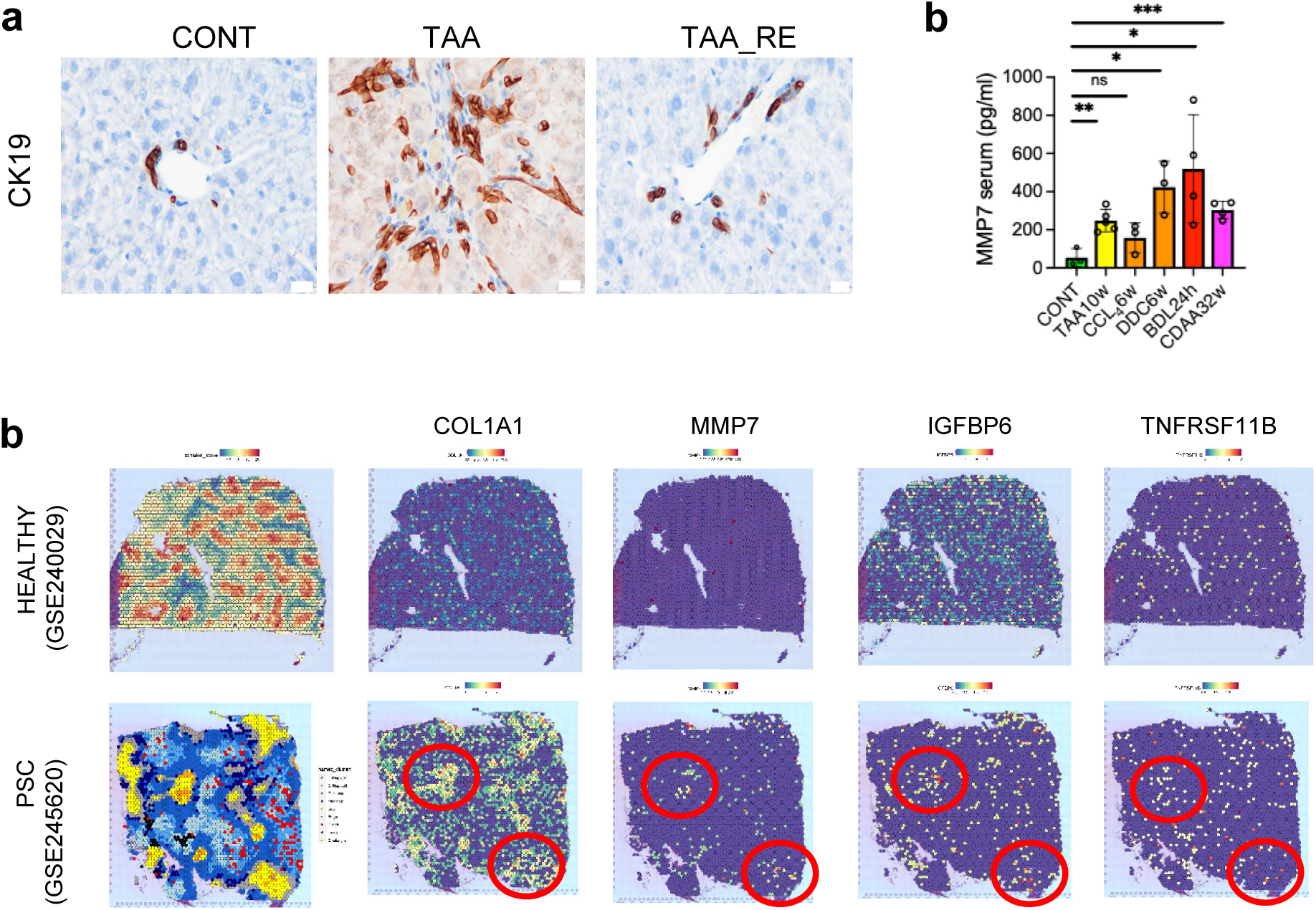
Cholangiocytes promotes HSCs activation via cytokine production. **a.** IHC staining showing CK19 expression in control, fibrosis and regressed group, scale bar 20 𝜇m. **b.** Barplot showing serum concentration of MMP7 in different mouse models. Data represent the means ± SD. *p*-values were analyzed by one-way ANOVA followed by Tukey’s multiple comparison test. \**p* < 0.05, \*\**p* < 0.01, \*\*\**p* < 0.001, \*\*\*\**p* < 0.0001. **c.** Spatial transcriptomics expression of COL1A1, MMP7, IGFBP6, and TNFRSF11B in healthy control (NDD, GSE240029) and PSC patient (PSC, GSE245620), scar area was covered with circle mark. **c.** Barplots showing quantitative data in Fig.5k

**Supplementary Fig. 10.**
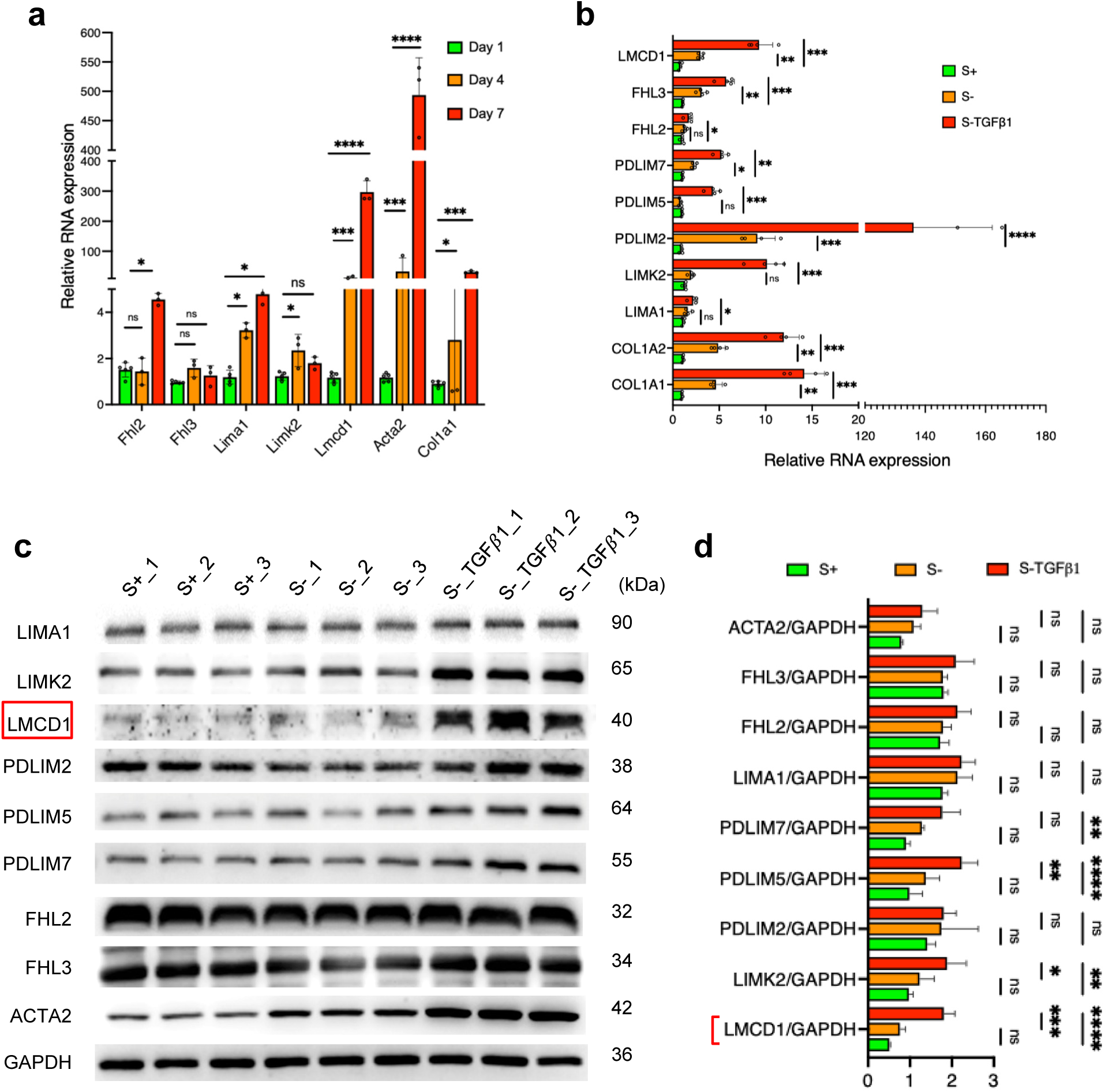
LIM family expression upregulated in mouse & human activated HSCs. **a.** Barplot showing RNA expression of Fhl2, Fhl3, Limk2, Lima1, Lmcd1, Acta2, Col1a1 in mHSCs at day 1, day 4, day 7 after isolation. Gapdh was used as internal control. **b.** Barplot showing RNA expression of LMCD1, FHL2, FHL3, PDLIM7, PDLIM5, PDLIM2, LIMK2, LIMA1, COL1A2, COL1A1 in HHSteCs cultured in medium with supplement (S+) or without supplement (S-) or without supplement and adding TGF𝛽1 (2 ng/ml). 18S was used as internal control. **c, d.** Immunoblotting showing LIMA1, LIMK2, LMCD1, PDLIM2, PDLIM5, PDLIM7, FHL2, FHL3, ACTA2 expression in HHSteCs cultured in medium with supplement (S+) or without supplement (S-) or without supplement and adding TGF𝛽 1 (2 ng/ml) and their quantification. GAPDH was used as internal control. For **a, b, d**. Data represent the means ± SD. *p*-values were analyzed by two-tailed Student’s t-test or by one-way ANOVA followed by Tukey’s multiple comparison test. \**p* < 0.05, \*\**p* < 0.01, \*\*\**p* < 0.001, \*\*\*\**p* < 0.0001

**Supplementary Fig. 11.**
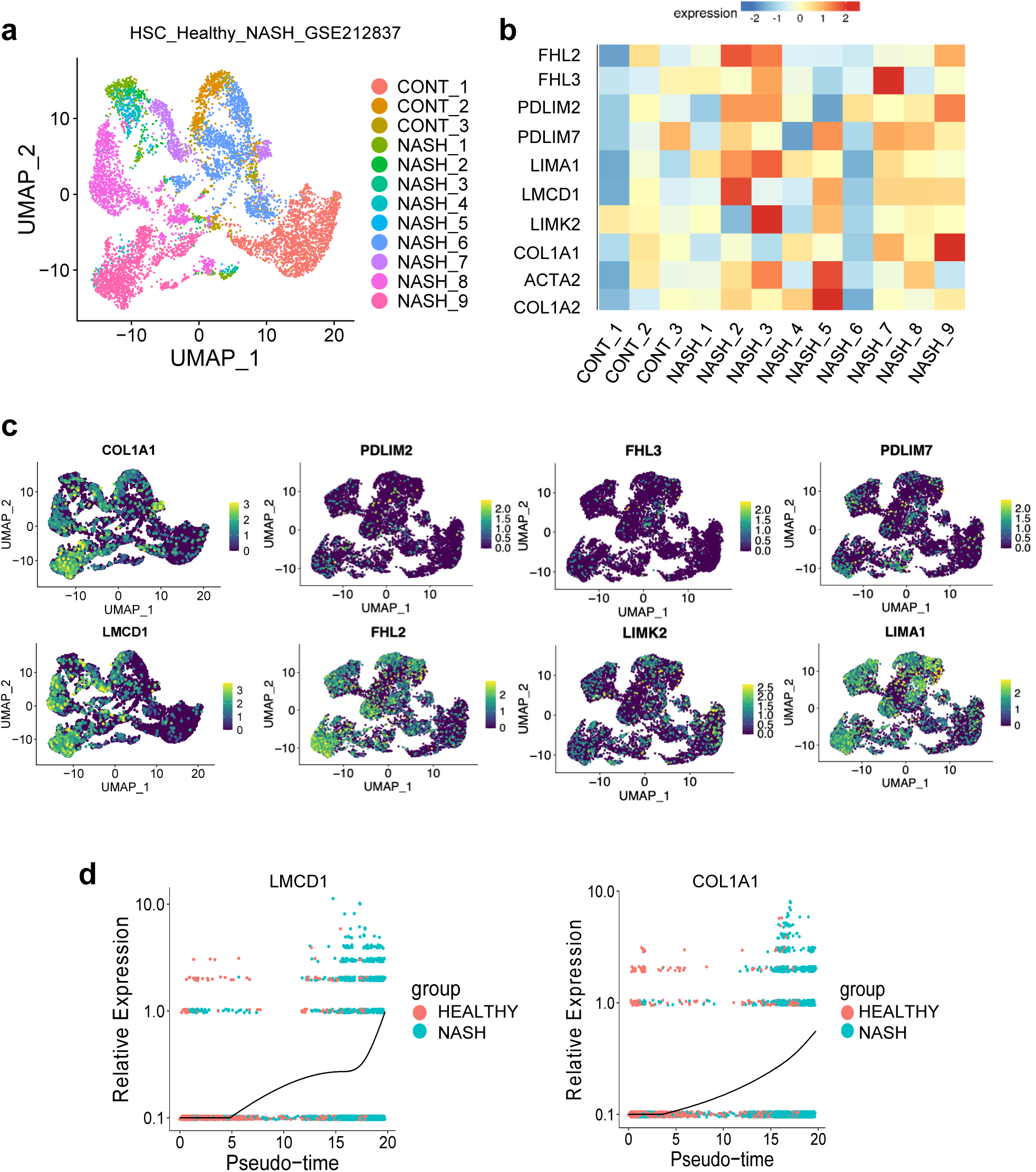
LMCD1 expressed in activated HSCs in human public dataset of NASH. **a.** UMAP showing HSCs subpopulation from public dataset (GSE212837). **b.** Heatmap showing expression of FHL2, FHL3, PDLIM2, PDLIM7, LIMA1, LMCD1, LIMK2, COL1A1, ACTA2, COL1A2 expression in healthy and NASH patients. **c.** UMAP showing expression of COL1A1, PDLIM2, FHL3, PDLIM7, LMCD1, FHL2, LIMK2, LIMA1, colored by gene expression. **d.** UMAP showing trajectory analysis of LMCD1 (left panel) and COL1A1 (right panel) alongside with pseudotime

**Supplementary Fig. 12.**
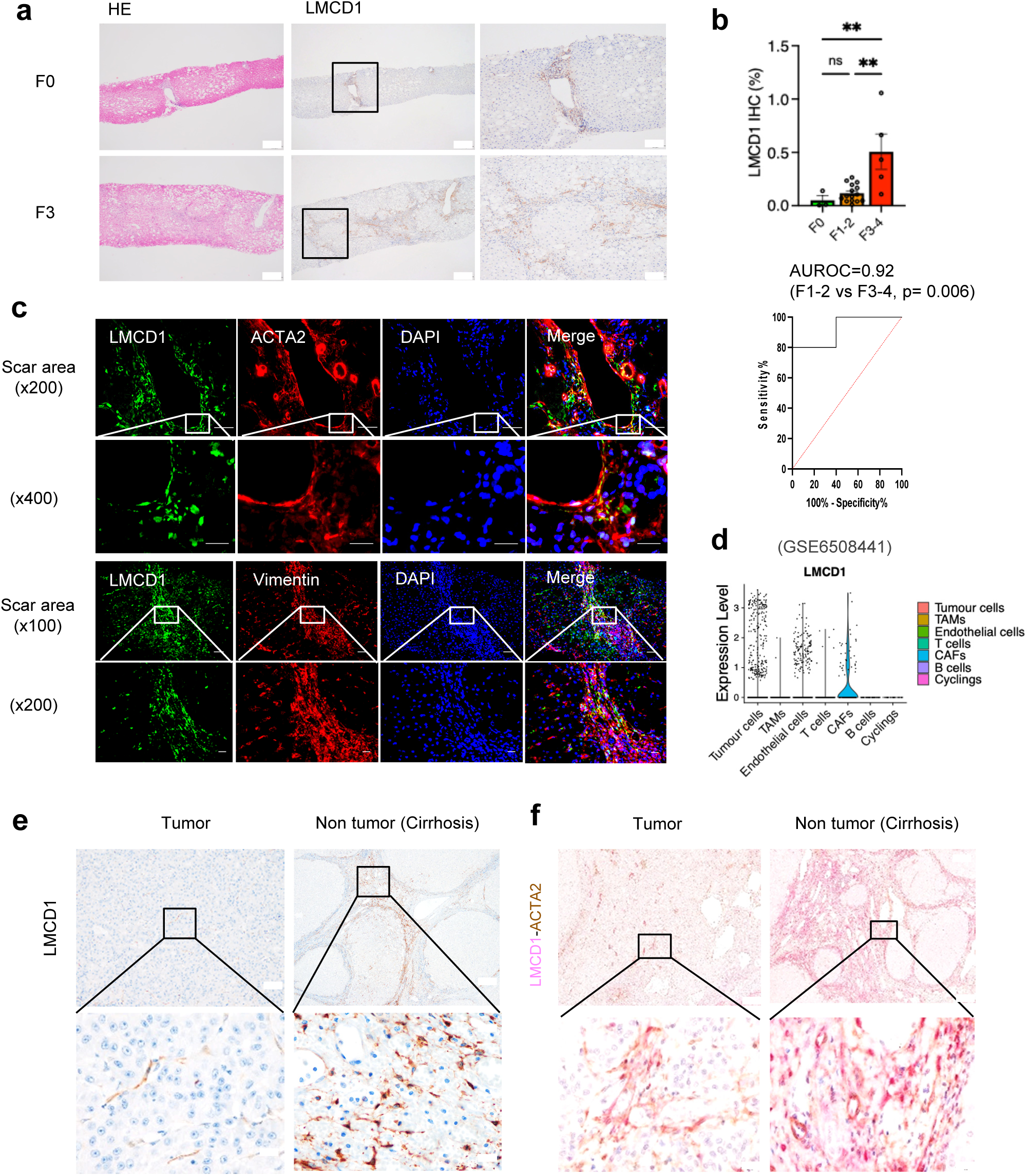
LMCD1 expression upregulated in scar area of human MASLD livers. **a.** Representative H&E and LMCD1 IHC staining in human MAFLD livers. Scale bar 200 𝜇m (left and middle panel), 100 𝜇m (right panel). **b.** Barplot showing LMCD1 positive area (%) in liver biopsy of F0, F1-2, and F3-4 MASLD patients (upper panel) and area under characteristic operating curve (AUROC) for predicting severity of fibrosis stage F3-4 compared to F1-2 in MASLD patient (lower panel). Data represent the means ± SD. *p*-values were analyzed by one-way ANOVA followed by Tukey’s multiple comparison test. \**p* < 0.05, \*\**p* < 0.01, \*\*\**p* < 0.001, \*\*\*\**p* < 0.0001. **c.** Double IF staining for LMCD1 (green) and ACTA2 (red) or Vimentin (red) in human cirrhosis livers. DAPI (blue) was used for visualizing nuclei. Scale bar 50 𝜇m (upper panel-1); 20 𝜇m (upper panel-2); 100 𝜇m (lower panel-1); 50 𝜇m (lower panel-2)**. d.** Violin plot showing expression of LMCD1 in human public database HCC (GSE6508441)**. e.** Expression of LMCD1 in tumor and non-tumor area of HCV-HCC patient. Scale bar 100 𝜇m (upper panel) and 50 𝜇m (lower panel). **f.** Double IHC showing expression of LMCD1 (Magenta) and ACTA2 (Brown) in tumor and non-tumor area of HCV-HCC patient. Scale bar 100 𝜇m (upper) and 50 𝜇m (lower).

**Supplementary Fig. 13.**
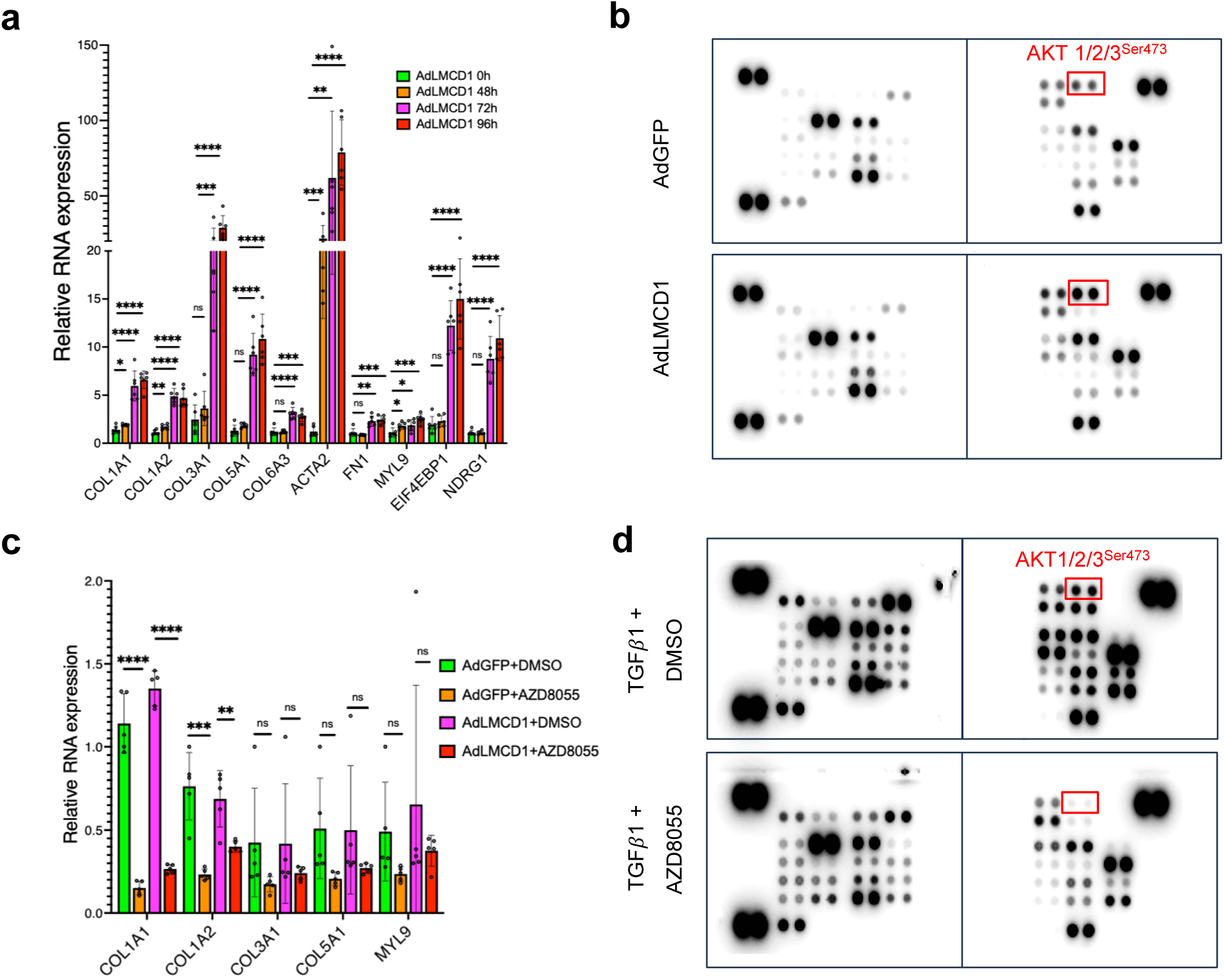
LMCD1 regulated HHSteCs activation via AKT signal. **a.** Barplot showing expression of COL1A1, COL1A2, COL3A1, COL5A1, COL6A3, ACTA2, FN1, MYL9, EIF4EBP1, NDGR1 in HHSteCs under overexpression LMCD1 time-dependently (AdLMCD1 0h, 48h, 72h, 96h). **b.** Immunoblotting showing expression of phosphoprotein alteration in HHsteCs under AdGFP or AdLMCD1. Alteration was marked with red. **c.** Barplot showing expression of COL1A1, COL1A2, COL3A1, COL5A1, MYL9 in HHSteCs under overexpression GFP or LMCD1 treated with Vehicle or AZD8055 at dose of 1 𝜇M. **d.** Immunoblotting showing expression of phosphoprotein alteration in HHSteCs under TGF𝛽1 treatment (2 ng/ml) treated with DMSO or AZD8055 at dose of 1 𝜇M. For **a, c.** Data represent the means ± SD. *p*-values were analyzed by two-tailed Student’s t-test. \**p* < 0.05, \*\**p* < 0.01, \*\*\**p* < 0.001, \*\*\*\**p* < 0.0001.

**Supplementary Fig. 14.**
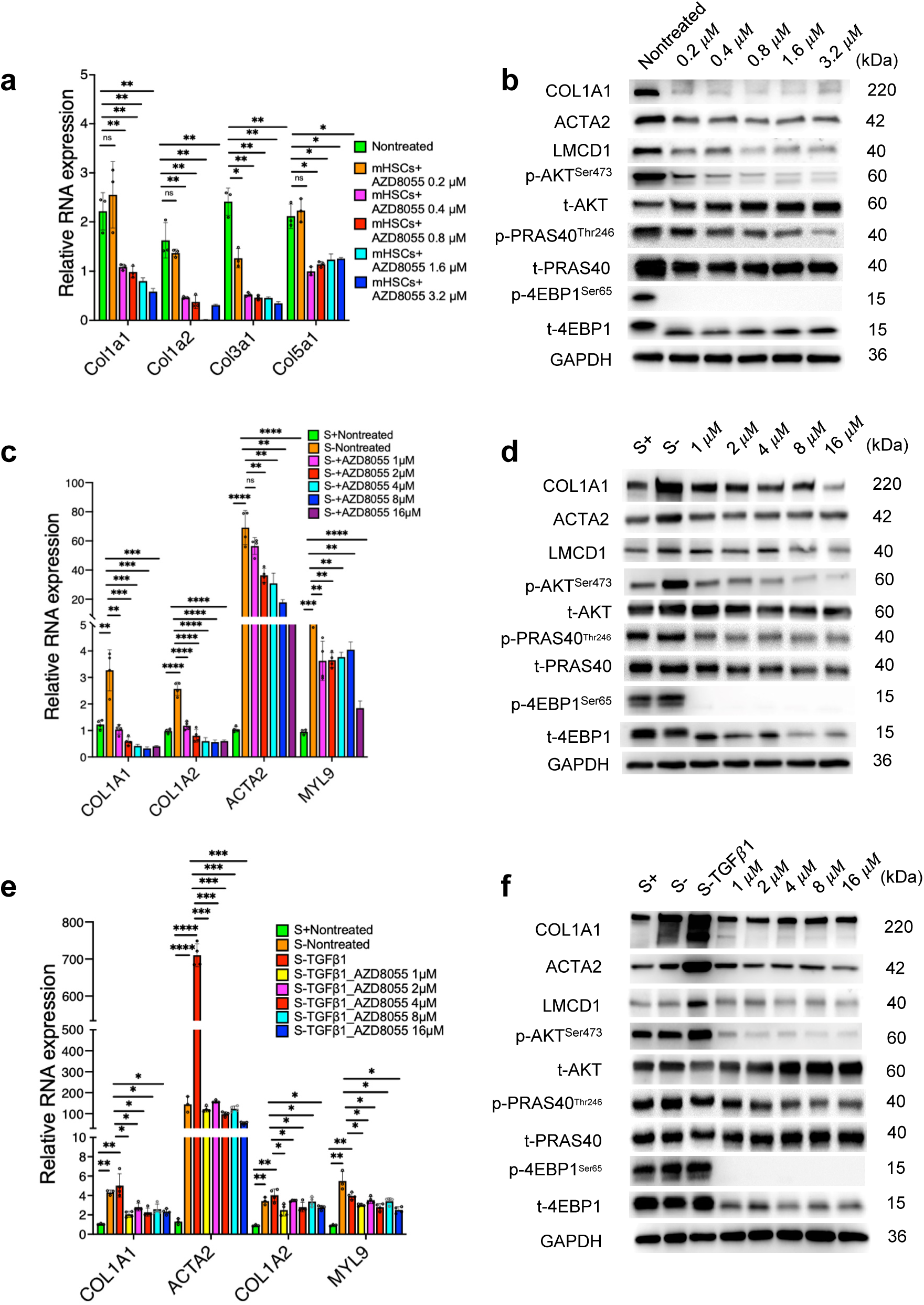
AZD8055 mitigated primary mHSCs and HHSteCs activation via LMCD1-AKT-PRAS40-4EBP1. **a.** Barplot showing Col1a1, Col1a2, Col3a1, and Col5a1 expression in mHSCs treated with AZD8055 dose-dependently**. b.** Immunoblotting showing expression of COL1A1, ACTA2, p-AKT^Ser473^, t-AKT, p-PRAS40^Thr246^, t-PRAS40, p-4EBP1^Ser65^, t-4EBP1, and LMCD1 in mHSCs treated with AZD8055 dose-dependently. GAPDH was used as internal control. **c.** Barplot showing COL1A1, COL1A2, ACTA2, and MYL9 expression in HHSteCs cultured in medium without supplement (S-) and treated with AZD8055 dose-dependently. 18S was used as internal control**. d.** Immunoblotting showing expression of COL1A1, ACTA2, p-AKT^Ser473^, t-AKT, p-PRAS40^Thr246^, t-PRAS40, p-4EBP1^Ser65^, t-4EBP1, LMCD1 in HHSteCs cultured in medium without supplement (S-) and treated with AZD8055 dose-dependently. GAPDH was used as internal control. **e.** Barplot showing COL1A1, COL1A2, ACTA2, and MYL9 expression in HHSteCs under TGF𝛽1 treatment (2 ng/ml) and treated with AZD8055 dose-dependently. **f.** Immunoblotting showing expression of COL1A1, ACTA2, p-AKT^Ser473^, t-AKT, p-PRAS40^Thr246^, t-PRAS40, p-4EBP1^Ser65^, t-4EBP1, LMCD1 in HHSteCs under TGF𝛽1 treatment (2 ng/ml) and treated with AZD8055 dose-dependently. GAPDH was used as internal control. For **a, c, e**. 18S was used as internal control. Data represent the means ± SD. *p*-values were analyzed by two-tailed Student’s t-test. \**p* < 0.05, \*\**p* < 0.01, \*\*\**p* < 0.001, \*\*\*\**p* < 0.0001

**Supplementary Fig. 15.**
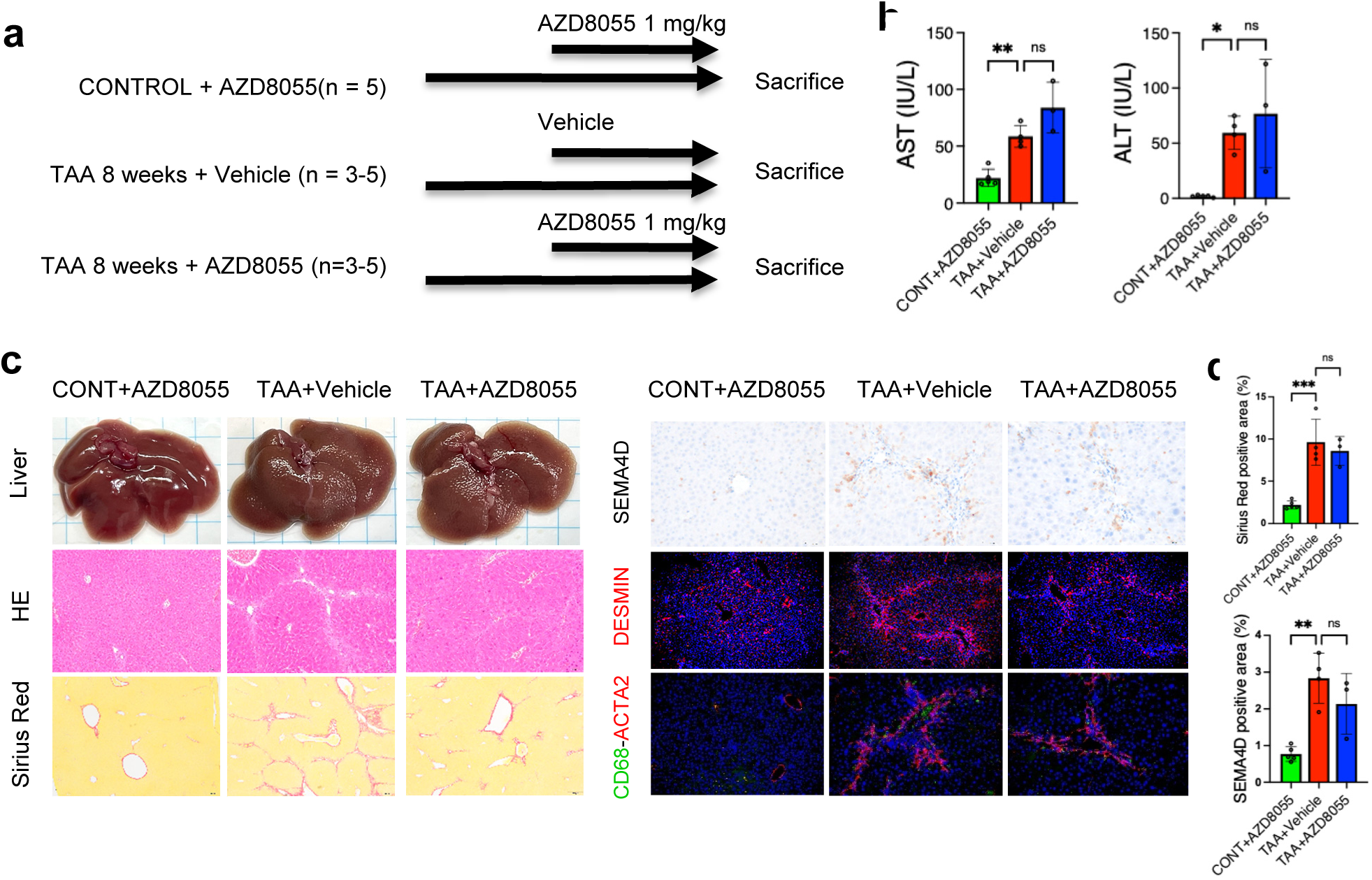
AZD8055 attenuated TAA-induced liver fibrosis in mice. **a.** Schematic overview of experimental design for **b-c. b.** Measurement serum level of AST, ALT among 3 groups (upper panel). **c.** Representative of livers, H&E, Sirius Red, IHC staining of SEMA4D, IF staining DESMIN, Double IF of CD68-ACTA2 staining of 3 groups: Control treated with AZD8055 1 mg/kg during the last 4 weeks, TAA 8 weeks treated with vehicle, and TAA 8 weeks treated with AZD8055 intraperitoneally during the last 4 weeks of TAA treatment at the dose of 1 mg/kg (n = 3-5, each group), scale bar 20 𝜇m. **d.** Quantification of Sirius Red and SEMA4D staining from panel c. Data represent the means ± SD. *p*-values were analyzed by one-way ANOVA followed by Turkey’s multiple comparison test. \**p* < 0.05, \*\**p* < 0.01, \*\*\**p* < 0.001, \*\*\*\**p* < 0.0001.

**Supplementary Fig. 16:**
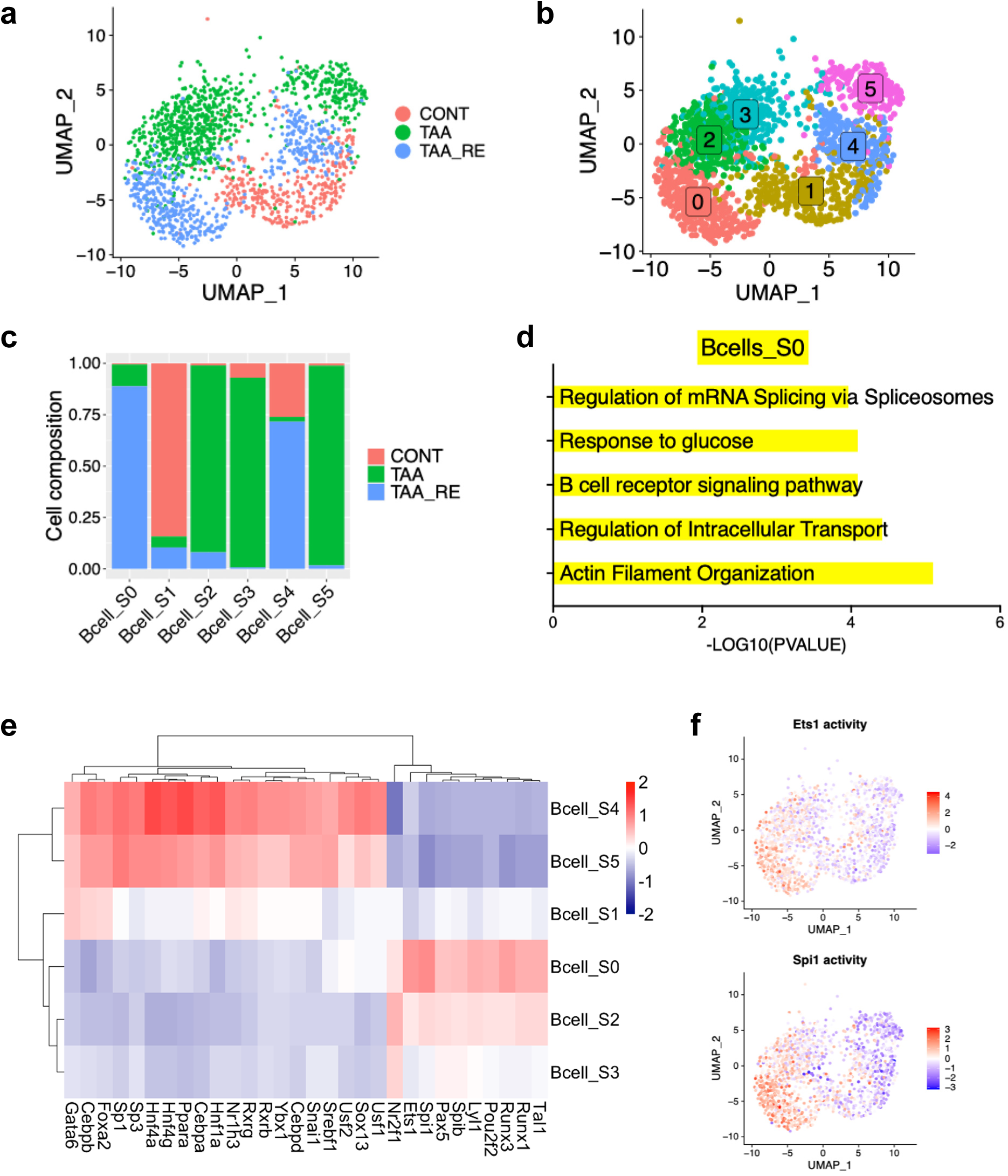
Heterogeneity of B cells in liver fibrosis progression and regression. **a.** UMAP showing B cells subpopulation from control, cirrhosis, and regression group, colored by group. **b.** UMAP showing 6 subpopulations of B cells from control, cirrhosis and the regression group, colored by cluster. **c.** Barplot showing group composition breakdown per cluster (fraction of total cell count per cluster), colored by group. **d.** Barplot showing top 5 GO pathway analyzed by EnrichR of B cell subcluster S0 compared by -log10(PVAL) of each pathway. **e.** Heatmap showing transcription factor analysis by Dorothea, colored by average expression. **f.** UMAP showing activity of transcription factor Ets1, Spi1 in B cell S0 subclusters

**Supplementary Fig. 17:**
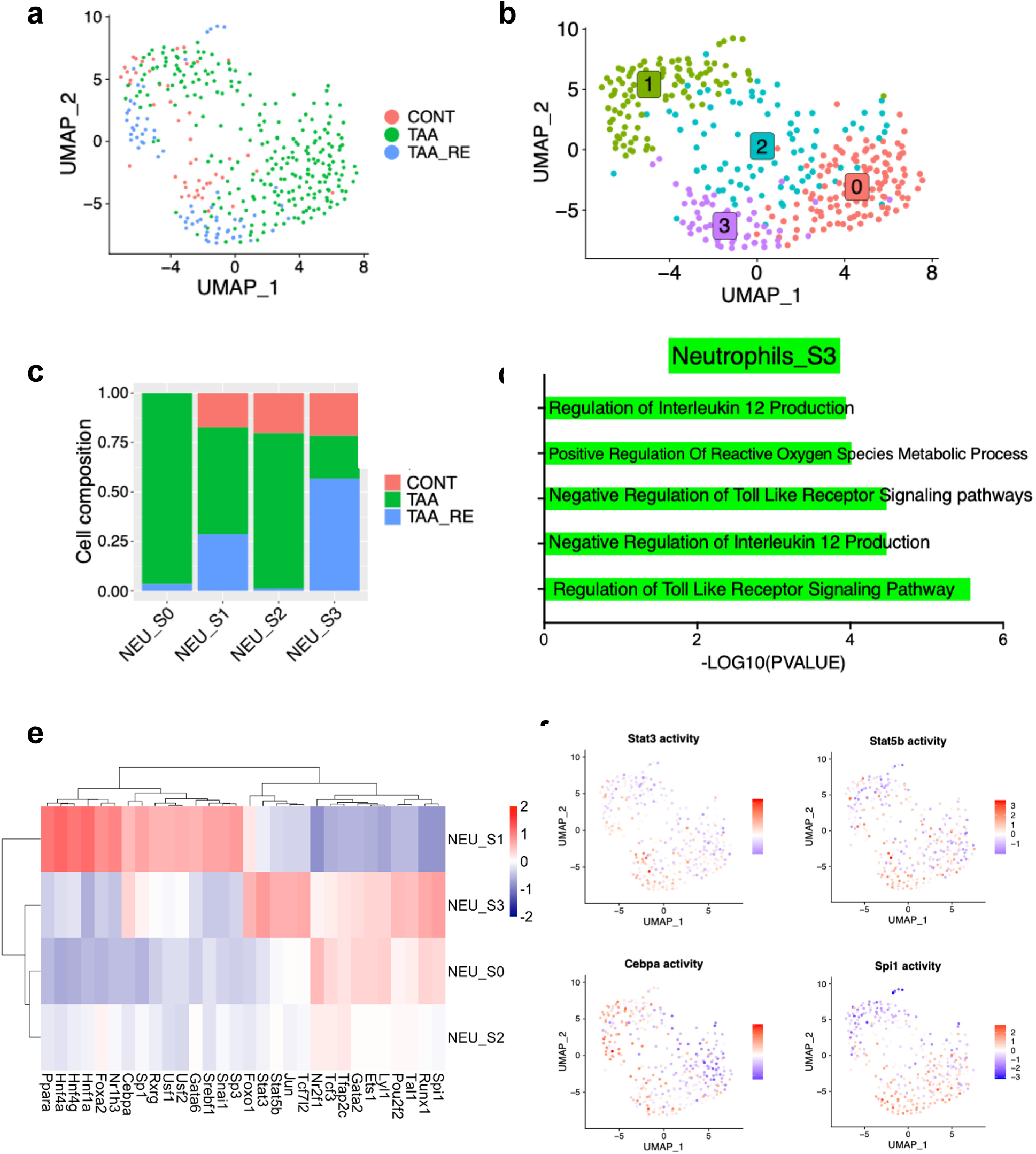
Heterogeneity of neutrophils in liver fibrosis progression and regression. **a.** UMAP showing neutrophils subpopulation from control, fibrosis, and regression group, colored by group. **b.** UMAP showing 4 subpopulations of neutrophils from control, cirrhosis and the regression group, colored by cluster. **c.** Barplot showing group composition breakdown per cluster (fraction of total cell count per cluster), colored by group. **d.** Barplot showing top 5 GO pathways analyzed by EnrichR of Neutrophils subcluster S3 compared by - log10(PVAL) of each pathway. **e.** Heatmap showing transcription factor analysis by Dorothea, colored by average expression. **f.** UMAP showing expression of transcription factor Stat3, Stat5b, Cebpa, and Spi1, colored by expression.

**Supplementary Fig. 18.**
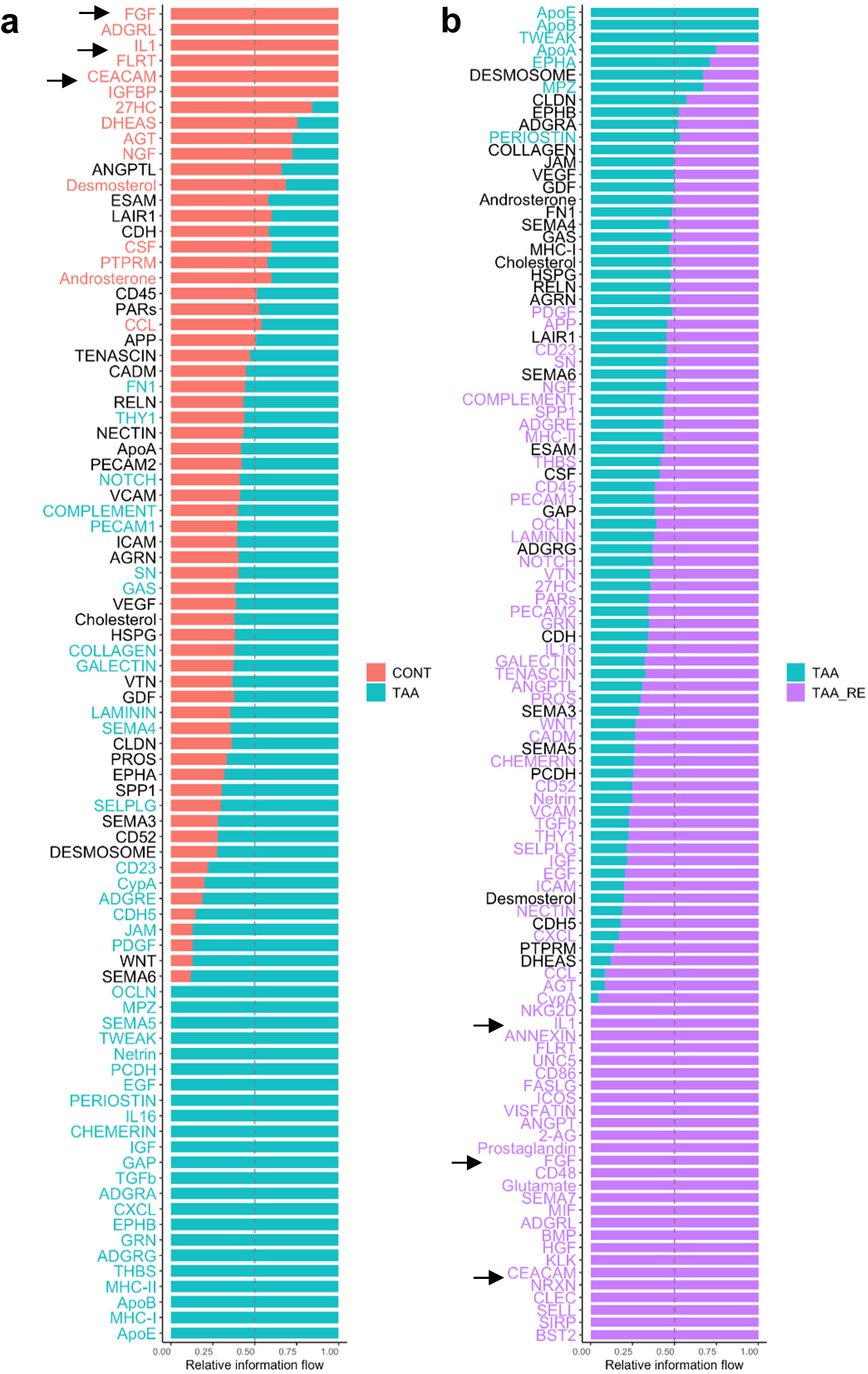
Relative information flow of pathways significantly enhanced or suppressed in healthy control, TAA, and regression group. **a.** Relative information flow in healthy control and TAA. **b.** Relative information flow in TAA and TAA_RE

**Supplementary Fig. 19.**
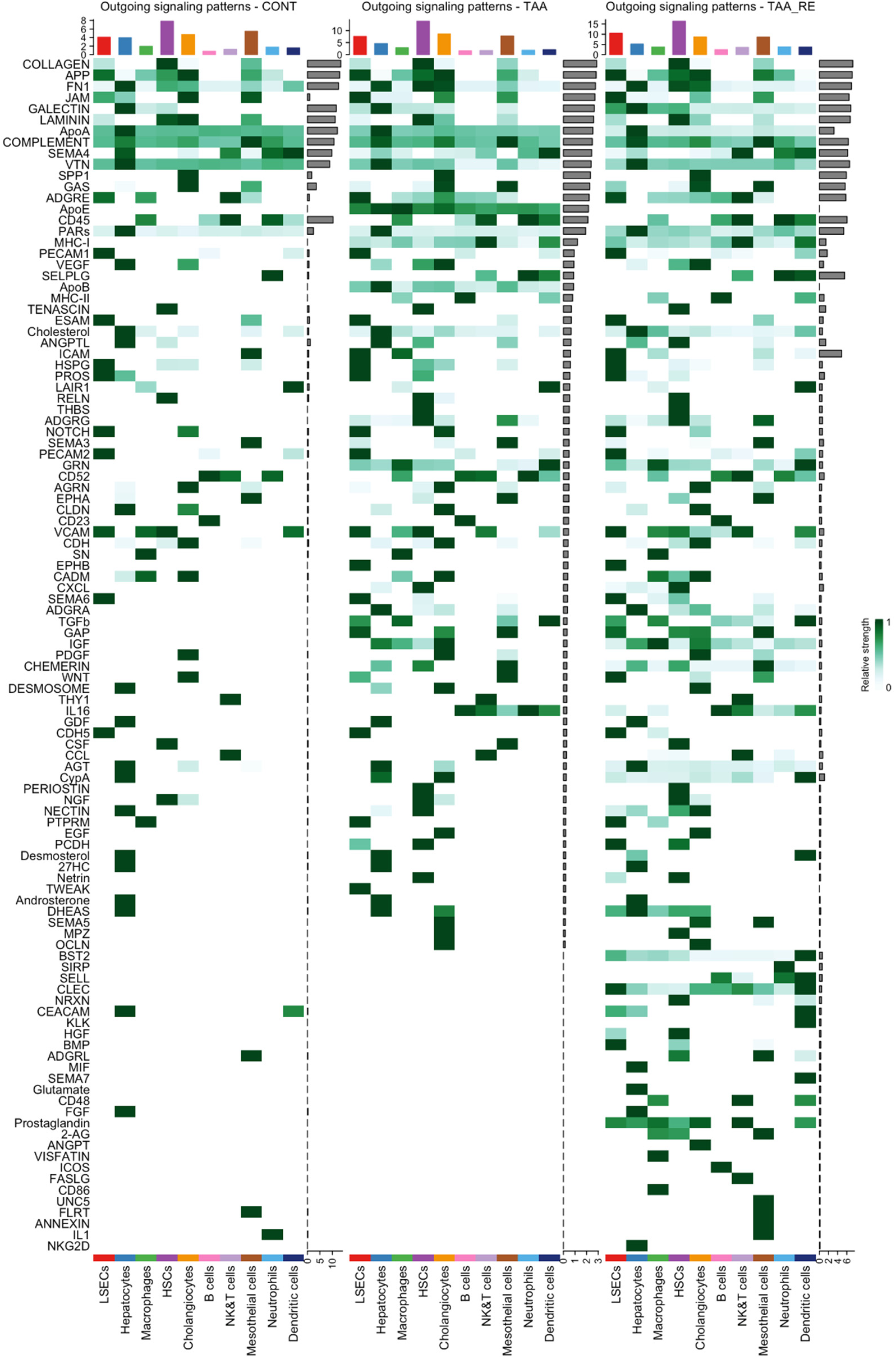
**Outgoing signaling patterns among cell types in, liver fibrosis progression (TAA) and regression (TAA-RE). healthy**

**Supplementary Fig. 20.**
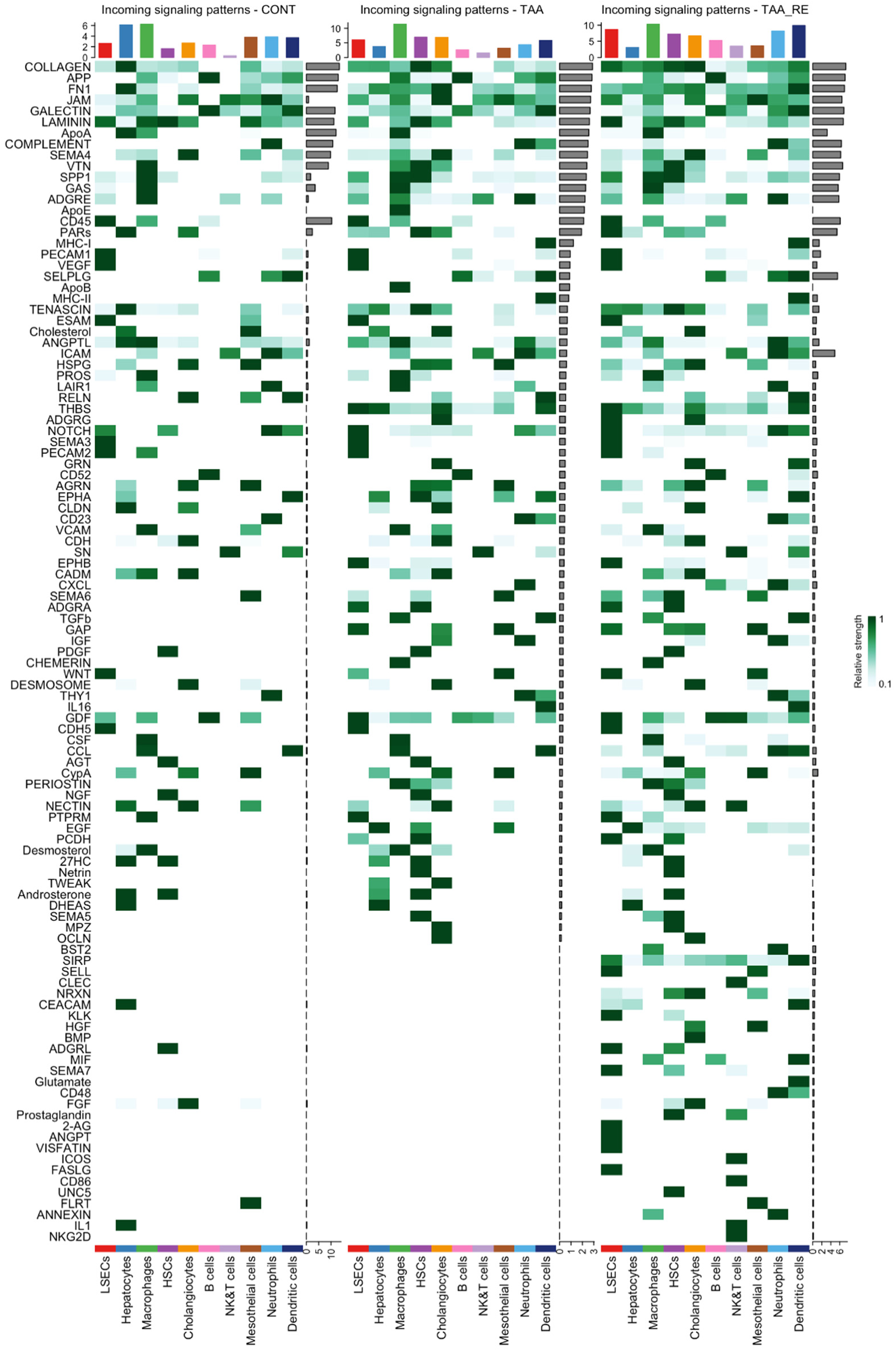
**Incoming signaling patterns among cell types in healthy, liver fibrosis progression (TAA) and regression (TAA-RE).**

