## Supplementary Data 8 for "A single-cell fixed RNA profiling of liver fibrosis progression and regression reveals SEMA4D and LMCD1 as key mediators of fibrogenesis"

**Supplementary Data 8.** List of mouse and human qRT-PCR primers used in this study

| Human primer | Sequence (5’-> 3’) |
| --- | --- |
| *18S* | Forward CGATCCGAGGGCCTCACTA  Reverse AGTCCCTGCCCTTTGTACACA |
| *α*SMA | Forward CAGCCAAGCACTGTCAGG  Reverse CCAGAGCCATTGTCACACAC |
| *COL1A1* | Forward AAGAGGAAGGCCAAGTCGAG  Reverse CACACGTCTCGGTCATGGTA |
| *COL3A1* | Forward CTGGACCCCAGGGTCTTC  Reverse CATCTGATCCAGGGTTTCCA |
| *COL4A1* | Forward TGGTGACAAAGGACAAGCAG  Reverse GGTTCACCCTTTGGACCTG |
| *MYL9* | Forward CGAGGATGTGATTCGCAACG  Reverse TGTTTGAGGATGCGGGTGAA |
| *FN1* | Forward CGGTGGCTGTCAGTCAAAG  Reverse AAACCTCGGCTTCCTTCCATAA |
| *COL1A2* | Forward GGCCCTCAAGGTTTCCAAAGG  Reverse CACCCTGTGGTCCAACAACTC |
| *COL6A3* | Forward AGGTTTGCTCAGGGGTTCATA  Reverse AGCCGCACCATTTTTGACAT |
| *COL5A1* | Forward CTTGGCCCAAAGAAAACCCG  Reverse GTAGGTGACGTTCTGGTGGG |
| *EIF4EBP1* | Forward CTATGACCGGAAATTCCTGATGG  Reverse CCCGCTTATCTTCTGGGCTA |
| *NDGR1* | Forward CTCCTGCAAGAGTTTGATGTCC  Reverse TCATGCCGATGTCATGGTAGG |
| *RARA* | Forward AAGCCCGAGTGCTCTGAGA  Reverse TTCGTAGTGTATTTGCCCAGC |
| *RARB* | Forward TCCGAAAAGCTCACCAGGAAA  Reverse GGCCAGTTCACTGAATTTGTCC |
| *RARG* | Forward TGTCACCGCGACAAAAACTGT  Reverse CGAGGGGAAAGTCTCCTGA |
| *RARRES1* | Forward AAACCCCTTGGAAATAGTCAGC  Reverse GGAAAGCCAAATCCCAGATGAG |
| *RARRES2* | Forward ACTGCCCCATAGAGACCCAA  Reverse CACCACGCATCTCAGTGCT |
| *LMCD1* | Forward ACCCCAAGAAGAGGATGGCAA  Reverse CTGGCCCAGGGACATCTTTT |
| *LIMA1* | Forward GACTCCCAGGTTAAGAGTGAGG  Reverse TTGCAGGTGCCTGAAACTTCT |
| *LIMK2* | Forward GGATTCCCTCACCAACTGGTA  Reverse AGCCACCATAAAAGGCCCTG |
| *PDLIM2* | Forward TGGGGCTTCCGTATCACAG  Reverse CTCTGGCGGATCTTGCTCT |
| *PDLIM5* | Forward CATGGCCTACAATAAGGCACC  Reverse CAGTCCAGATGCAGCGAAC |
| *PDLIM7* | Forward ATGCCCCGGACAAAACGAG  Reverse TGGTGACACACGGGAGTCT |
| *FHL2* | Forward GTACAGACTGCTATTCCAACGAG  Reverse GCACTGCATGGCATGTTGTT |
| *FHL3* | Forward TCCCGGAAGCTGGAATATGGA  Reverse CCTTGTCGGGCACAAAAGAA |

| Mouse primer | Sequence (5’-> 3’) |
| --- | --- |
| *αSma* | Forward TCCCTGGAGAAGAGCTACGAACT  Reverse AAGCGTTCGTTTCCAATGGT |
| *Col1a1* | Forward GAGCGGAGAGTACTGGATCG  Reverse GTTCGGGCTGATGTACCAGT |
| *Col1a2* | Forward TCGTGCCTAGCAACATGCC  Reverse TTTGTCAGAATACTGAGCAGCAA |
| *Col3a1* | Forward CTGTAACATGGAAACTGGGGAAA  Reverse CCATAGCTGAACTGAAAACCACC |
| *Col5a1* | Forward ATACTGGGTCGATCCCAACC  Reverse CAAGAAGTGATTCTGGCTCCCT |
| *Sema4d* | Forward TGATCCCTAGGTCAGACGGG  Reverse CTGGCTTGTGAAACTGCACC |
| *Krt19* | Forward GTTCAGTACGCATTGGGTCAG  Reverse GAGGACGAGGTCACGAAGC |
| *Fn1* | Forward ATGTGGACCCCTCCTGATAGT  Reverse GCCCAGTGATTTCAGCAAAGG |
| *Spp1* | Forward CTTTCACTCCAATCGTCCCTAC  Reverse GCTCTCTTTGGAATGCTCAAGT |
| *Scube3* | Forward TTCACTGGAACGGGAAGGATTG  Reverse GGACAGGTCAGGGCACAAGTA |
| *Thbs1* | Forward TTCTTACCCTTGACAACAACGTG  Reverse CCACAGATAGCTTGGAGGTCC |
| *Lcn2* | Forward GGGAAATATGCACAGGTATCCTC  Reverse CATGGCGAACTGGTTGTAGTC |
| *Mmp7* | Forward CTTACCTCGGATCGTAGTGGA  Reverse CCCCAACTAACCCTCTTGAAGT |
| *Lmcd1* | Forward GCCTCACTCGTGGAGGAAAAT Reverse CCAGGTCAGAGCTTAGGCAG |
| *Lima1* | Forward AGCCAGAGACGAGCGAAAAC  Reverse CCCGCACCTATTTCCCAGTC |
| *Limk2* | Forward GGGCTGTGGCACCTATGTTC  Reverse CCAGTTGGTGAGGGATTCCTG |
| *Fhl2* | Forward ATGACTGAACGCTTTGACTGC  Reverse CGATGGGTGTTCCACACTCC |
| *Fhl3* | Forward ATGAGCGAGGCATTTGACTGT  Reverse GTCATAGCACGGAACGCAGTA |
| *Rarres2* | Forward CACACAGAAAAAGGCCTCGC  Reverse AGGGCTAGGGAGATCAGCAA |
| *Gapdh* | Forward TGCACCACCAACTGCTTAG  Reverse GGATGCAGGGATGATGTTC |
