## Supplementary Data 9 for "A single-cell fixed RNA profiling of liver fibrosis progression and regression reveals SEMA4D and LMCD1 as key mediators of fibrogenesis"

**Supplementary Data 9.** List of antibodies used in this study.

| Immunohistochemical and Immunocyto/Immunofluorescent Staining | | | | | |  |  |  |  |
| --- | --- | --- | --- | --- | --- | --- | --- | --- | --- |
| Antibody | Species | | Source | Cat# | Dilution |  |  |  |  |
| Anti-$\alpha$SMA | Rabbit | Polyclonal | Abcam | Ab5694 | 1/100 |  |  |  |  |
| Anti-*α*SMA | Mouse | Monoclonal | Dako | M0851 | 1/100 |  |  |  |  |
| Anti-Ki67 | Rabbit | Polyclonal | Abcam | Ab16667 | 1/100 |  |  |  |  |
| Anti-CD68 | Rabbit | Polyclonal | Abcam | Ab125212 | 1/100 |  |  |  |  |
| Anti-CD31 | Goat | Polyclonal | R&D | AF3628 | 1/100 |  |  |  |  |
| Anti-Glutamin synthetase (GS) | Rabbit | Polyclonal | Abcam | Ab73593 | 1/100 |  |  |  |  |
| Anti-CYP2E1 | Rabbit | Monoclonal | Abcam | Ab19140 | 1/100 |  |  |  |  |
| Anti-LYVE1 | Rabbit | Polyclonal | Abcam | Ab281587 | 1/100 |  |  |  |  |
| Anti-P21 | Rabbit | Monoclonal | Abcam | Ab188224 | 1/100 |  |  |  |  |
| Anti-SEMA4D (Cd100) | Rabbit | Monoclonal | Cell Signaling | #53108 | 1/100 |  |  |  |  |
| Anti-CK19 | Rabbit | Monoclonal | Abcam | Ab52635 | 1.100 |  |  |  |  |
| Anti-DESMIN | Rabbit | Monoclonal | Invitrogen | PA5-16705 | 1/100 |  |  |  |  |
| Anti-LMCD1 | Rabbit | Polyclonal | Abcam | Ab121788 | 1/100 |  |  |  |  |
| Anti-VIMENTIN | Rat | Monoclonal | R&D | MAB2105 | 1/100 |  |  |  |  |
| Anti-E-CADHERIN | Rabbit | Polyclonal | Abcam | Ab53033 | 1/100 |  |  |  |  |
| Western Blotting analysis | | | | | |  |  |  | 1/100 |
| Anti-*α*SMA | Rabbit | Polyclonal | Abcam | Ab5694 | 1/1000 |  |  |  |  |
| Anti-*α*SMA | Mouse | Monoclonal | Dako | M0851 | 1/1000 |  |  |  |  |
| Anti-HO-1 | Rabbit | Polyclonal | Enzo | ADI-SPA-895-F | 1/1000 |  |  |  |  |
| Anti-CYP2E1 | Rabbit | Monoclonal | Abcam | Ab19140 | 1/1000 |  |  |  |  |
| Anti-GAPDH | Mouse | Monoclonal | EMD Milipore | MAB374 | 1/1000 |  |  |  |  |
| Anti-P21 | Rabbit | Monoclonal | Abcam | Ab188224 | 1/5000 |  |  |  |  |
| Anti-COL1A1 | Rabbit | Monoclonal | Abcam | Ab138492 | 1/1000 |  |  |  |  |
| Anti-COL1A1 | Rabbit | Monoclonal | Cell signaling | 72026S | 1/1000 |  |  |  |  |
| Anti-SEMA4D (CD100) | Rabbit | Monoclonal | Cell Signaling | 53108 | 1/1000 |  |  |  |  |
| Anti-PLEXIN B2 | Rabbit | Monoclonal | Abcam | Ab193355 | 1/1000 |  |  |  |  |
| Anti-LMCD1 | Mouse | Monoclonal | Santa Cruz | sc515171 | 1/1000 |  |  |  |  |
| Anti-BAX | Rabbit | Polyclonal | Cell Signaling | 2772S | 1/1000 |  |  |  |  |
| Anti-p-AKT^Ser473^ | Rabbit | Polyclonal | Cell Signaling | 9271 | 1/1000 |  |  |  |  |
| Anti-t-AKT | Rabbit | Monoclonal | Cell Signaling | 4691S | 1/1000 |  |  |  |  |
| Anti-p-PRAS40^Thr246^ | Rabbit | Monoclonal | Cell Signaling | 13175 | 1/1000 |  |  |  |  |
| Anti-t-PRAS40 | Rabbit | Monoclonal | Cell Signaling | 2691 | 1/1000 |  |  |  |  |
| Anti-p-4EBP1^Ser65^ | Rabbit | Monoclonal | Cell Signaling | 293124 | 1/1000 |  |  |  |  |
| Anti-t-4EBP1 | Rabbit | Monoclonal | Cell Signaling | 9451 | 1/1000 |  |  |  |  |
| Anti-p-SMAD3 | Rabbit | Monoclonal | Cell Signaling | 9644 | 1/1000 |  |  |  |  |
| Anti-t-SMAD3 | Rabbit | Polyclonal | Cell Signaling | 9513 | 1/1000 |  |  |  |  |
| Anti-LIMK2 | Mouse | Monoclonal | Santa Cruz | 365414 | 1/1000 |  |  |  |  |
| Anti-PDLIM2 | Mouse | Monoclonal | Santa Cruz | 515630 | 1/1000 |  |  |  |  |
| Anti-PDLIM5 | Mouse | Monoclonal | Santa Cruz | 515621 | 1/1000 |  |  |  |  |
| Anti-PDLIM7 | Mouse | Monoclonal | Santa Cruz | 98100 | 1/1000 |  |  |  |  |
| Anti-LIMA1 | Mouse | Monoclonal | Santa Cruz | 136399 | 1/1000 |  |  |  |  |
| Anti-FHL2 | Mouse | Monoclonal | Santa Cruz | 393514 | 1/1000 |  |  |  |  |
| Anti-FHL3 | Mouse | Monoclonal | Santa Cruz | 166917 | 1/1000 |  |  |  |  |
| Anti-FOLR2 | Rabbit | Monoclonal | Abcam | 302532 | 1/1000 |  |  |  |  |

List of flow cytometry antibodies used in this study.

|  | Name | Conjun  -gate | Target | Host | Source | Clone | Catalogue number | Lot number | Use | Conc  /10^6^ cell |
| --- | --- | --- | --- | --- | --- | --- | --- | --- | --- | --- |
| CD45.2 | | FITC | Mouse | Mouse | Biolegend | 104 | 109805 | B291435 | FCM |  |
| CD45.2 | | PE | Mouse | Mouse | Biolegend | 104 | 109808 | B361505 | FCM |  |
| CD45.2 | | AF700 | Mouse | Mouse | Biolegend | 104 | 109821 | B354902 | FCM |  |
| CD45.2 | | BV421 | Mouse | Mouse | Biolegend | 104 | 109831 | B368572 | FCM |  |
| CD45.2 | | BV510 | Mouse | Mouse | Biolegend | 104 | 109837 | B251593 | FCM |  |
| CD45.2 | | BV605 | Mouse | Mouse | Biolegend | 104 | 109841 | B340360 | FCM |  |
| CD45 | | BV711 | Mouse | Mouse | Biolegend | 30-F11 | 103147 | B206778 | FCM |  |
| CD11B | | FITC | Mouse | Rat | Biolegend | M1/70 | 101205 | B369619 | FCM |  |
| FOLR2 | | APC | Mouse | Rat | Biolegend | 10/FR2 | 153305 | B372041 | FCM |  |
| F4/80 | | BV510 | Mouse | Rat | Biolegend | BM8 | 123135 | B386407 | FCM |  |
| Ly6G | | BV605 | Mouse | Rat | Biolegend | 1A8 | 127639 | B393005 | FCM |  |
| MHCII | | BV711 | Mouse | Rat | Biolegend | M5/114.15.2 | 107643 | B399082 | FCM |  |
| Cd146 | | FITC | Mouse | Rat | Biolegend | ME-9F1 | 134705 | B341624 | FCM |  |
| NK1.1 | | PE | Mouse | Mouse | Biolegend | S17016D | 156503 | B375448 | FCM |  |
| Anti-mouse CD16/32 | |  | Mouse |  | InVivoMab | 2.4G2 | BE0307 |  | FCM |  |
| Zoombie NIR Fixable Viability Kit | |  |  |  | Biolegend |  | 423105 |  | FCM |  |
